# Expression and processing of polycistronic artificial microRNAs and *trans*-acting siRNAs in *Solanum lycopersicum* and *Nicotiana benthamiana*

**DOI:** 10.1101/2021.01.06.425596

**Authors:** Alice Lunardon, Samwel Muiruri Kariuki, Michael J. Axtell

**Affiliations:** Department of Biology, The Pennsylvania State University, University Park, Pennsylvania 16802, USA; Department of Molecular Biology, Princeton University, Princeton, New Jersey 08544-1014, USA; Department of Chemical Engineering, The Pennsylvania State University, University Park, Pennsylvania 16802, USA; International Institute of Tropical Agriculture, P.O. Box 30709-00100 and Department of Plant Sciences, Kenyatta University P.O. Box 43844-00100 Nairobi, Kenya; Huck Institutes of the Life Sciences, The Pennsylvania State University, University Park, Pennsylvania 16802, USA

## Abstract

Targeted gene silencing using small regulatory RNAs is a widely used technique for genetic studies in plants. Artificial microRNAs are one common approach; they have the advantage of producing just a single functional small RNA which can be designed for high target specificity and low off-target effects. Simultaneous silencing of multiple targets with artificial microRNAs can be achieved by producing polycistronic microRNA precursors. Alternatively, specialized *trans*-acting short interfering RNA (tasiRNA) precursors can be designed to produce several specific tasiRNAs at once. Here we tested several artificial microRNA- and tasiRNA-based methods for multiplexed gene silencing in *Solanum lycopersicum* (tomato) and *Nicotiana benthamiana*. Small RNA sequencing analyses revealed that many previously described approaches resulted in poor small RNA processing. The 5’-most microRNA precursor hairpins on polycistronic artificial microRNA precursors were generally processed more accurately than precursors at the 3’ end. Polycistronic artificial microRNAs where the hairpin precursors were separated by transfer RNAs had the best processing precision. Strikingly, artificial tasiRNA precursors failed to be processed in the expected phased manner in our system. These results highlight the need for further development of multiplexed artificial microRNA and tasiRNA strategies. The importance of small RNA sequencing, as opposed to single-target assays such as RNA blots or real-time PCR, is also discussed.

**Significance statement:** Several strategies for multiplexed gene silencing using artificial microRNAs or tasiRNAs have been described. We find that many result in imprecise processing, and thus low accumulation of the intended small RNAs. Our findings highlight the importance of small RNA sequencing to fully analyze gene silencing experiments, and also the need for continued methodological development of these methods.

## Introduction

Plant endogenous small RNAs (sRNAs) are a class of RNAs ranging in size between 20 and 24 nucleotides that act in RNA silencing processes, specifically targeting one or multiple RNAs with sequence complementarity, inducing their downregulation. MicroRNAs (miRNAs) and *trans*-acting small interfering RNAs (tasiRNAs) are functionally similar but have different biogenesis pathways. *MIRNA* genes are transcribed by Polymerase II (Pol II) into a long stem-loop RNA precursor that is processed by DICER-LIKE 1 (DCL1) into a 21- or 22-nucleotide duplex, formed by two strands: the mature miRNA, and the miRNA*. Once exported to the cytoplasm, the miRNA* is usually degraded and the miRNA is loaded into an ARGONAUTE (AGO) protein (Rogers and Chen, 2013). *TAS* gene transcripts are produced by Pol II and cleaved by a 22-nucleotide microRNA, leading to their conversion into double stranded RNA (dsRNA) by RNA-DEPENDENT RNA POLYMERASE6 (RDR6). The dsRNA is then sequentially processed by DCL4 into 21-nucleotide duplexes, in a phased pattern relative to the initial miRNA cut site, and one strand of the tasiRNA duplex is selectively incorporated into an AGO protein (Allen et al., 2005). MiRNAs and tasiRNAs silence their complementary target RNAs by direct AGO-mediated endonucleolytic cleavage and subsequent degradation (Jones-Rhoades et al., 2006). MiRNAs can also repress their targets by inhibiting mRNA translation (Gandikota et al., 2007).

These endogenous biogenesis and silencing machineries can be redirected by artificial miRNAs and artificial tasiRNAs (amiRNAs and atasiRNAs) to suppress specific RNAs and generating loss-of-function transient or stable transformants for genes of interest. An *aMIRNA* or *aTAS* gene is created by modifying a native precursor by replacing the miRNA/miRNA* or tasiRNA duplexes with custom sequences that are complementary to the RNA to downregulate (Schwab et al., 2006; Gutiérrez-Nava et al., 2008). The choice of the precursor backbone contributes to the success or failure of producing the artificial sRNAs (asRNAs) in a specific plant system and therefore of silencing their targets.

Many plant monocistronic *MIRNA* genes (monocistronic in this context indicates the presence of one single *MIRNA* hairpin) have been used to express amiRNAs in dicots and monocots, including *ath-MIR159* (Niu et al., 2006), *tae-MIR164* (Gasparis et al., 2017), *ath-MIR167b*, *ath-MIR171a* (Qu et al., 2007; Ai et al., 2011), *ath-MIR172a* (Schwab et al., 2006), *ath-MIR319a* (Schwab et al., 2006; Jover-Gil et al., 2014), *ath-MIR390a* (Carbonell et al., 2014), *osa-MIR390* (Carbonell et al., 2015), *ath-MIR395a* (Liang et al., 2012), and *osa-MIR528* (Warthmann et al., 2008). Amongst these, the *ath-MIR319a* and *osa-MIR528* backbones have been used most commonly because they are highly conserved across the plant kingdom (Zhang et al., 2019). Recently, the *ath-MIR390a* foldback was demonstrated to be very efficient, expressing significantly higher levels of amiRNAs than *ath-MIR319a* in *Nicotiana benthamiana* (Carbonell et al., 2014). In tomato, the following miRNA backbones have been used to express amiRNAs and obtain loss-of-function transgenic plants: *ath-MIR159a* (Zhang et al., 2011), the endogenous *sly-MIR159a* (Kravchik et al., 2014) and *ath-MIR319a* (Sharma and Prasad, 2020; Yogindran and Rajam, 2020). In a comparative study, (Vu et al., 2013) showed that in tomato, the same amiRNAs are more abundantly expressed from *sly-MIR159a* than *ath-MIR319a*, while the *sly-MIR168a* backbone was not able to produce any amiRNA.

Polycistronic constructs carry multiple *MIRNA* hairpins on the same transgene under a single promoter, to reinforce the silencing of a gene or targeting different genes simultaneously without using multiple expression cassettes. The simplest design is the tandem repeat of multiple miRNA backbones, where every hairpin carries a different amiRNA/amiRNA* duplex. The miRNA backbones can be identical, for example two or three consecutive *ath-MIR159a* (Ai et al., 2011; Kung et al., 2012), three tandem *hvu-MIR171* (Kis et al., 2016), two or three tandem *ath-MIR319a* (Park et al., 2009), or they can be different, for example the dimeric *ath-MIR319a* and *ath-MIR395a* construct used in *Arabidopsis thaliana* (Liang et al., 2012). The efficiency at which a polycistronic RNA is processed into its individual miRNA backbones directly influences the level of amiRNA expression. To increase the processing precision and the production efficiency of single RNAs from a synthetic polycistronic gene, the transfer RNA (tRNA) sequence can be placed upstream of each individual RNA, as was first demonstrated in rice for the generation of multiple Cas9-associated guide RNAs (Xie et al., 2015). The same strategy was later adopted in *A. thaliana*, where a tRNA sequence was introduced in a polycistronic *aMIRNA* gene, upstream of each of five tandem *ath-MIR319a* precursors, to efficiently express five different amiRNAs (Zhang et al., 2018). The endogenous tRNA-processing machinery is an efficient, robust cellular component that can process different types of substrates and it is conserved across monocot and dicot species, making this strategy ideal for polycistronic *aMIRNA* genes (Xie et al., 2015; Zhang et al., 2018). A second possibility for expressing multiple amiRNAs in the same gene is using plant endogenous polycistronic *MIRNA* genes, changing the native miRNA/miRNA* duplexes with amiRNA/amiRNA* sequences. Using the rice endogenous polycistronic *MIR395a-g* locus, that is 1,055bp-long and contains seven miRNA hairpins, (Fahim et al., 2012) were able to express five amiRNAs in wheat, by replacing the natural miR395 in each of the first five arms of *osa-MIRR395a-g* with the amiRNA sequences. Similarly, endogenous *TAS* genes can be engineered to express multiple atasiRNAs from the same precursor, with the advantage that atasiRNA vectors don’t have a requirement for secondary structure as amiRNA vectors. The *A. thaliana* endogenous *TAS1c* gene has been engineered to replace two or five of the native tasiRNA sequences with atasiRNAs, targeting the same or different genes (Gutiérrez-Nava et al., 2008; Carbonell et al., 2014). The *TAS1c* gene and the miRNA that triggers the production of tasiRNAs from the *TAS1c* transcript, miR173, is only present in *A. thaliana* and close relatives. Therefore, in order to use this system in other species, the *ath-MIR173* gene has to be co-introduced and co-expressed with the artificial *TAS1c* gene (*aTAS1c*). In *N. benthamiana*, the *aTAS1c* and the *ath-MIR173* genes have been co-expressed from two independent constructs that are co-infiltrated in the plant (Zhao et al., 2015; Carbonell and Daròs, 2017). Alternatively, in *A. thaliana*, tobacco and tomato, the *aTAS1c* and the *ath-MIR173* genes have been co-expressed from the same plasmid, in the same or different promoter-terminator cassettes (Baykal et al., 2016; Carbonell et al., 2019). The *TAS3* gene has also been engineered to express multiple atasiRNAs. Because both *TAS3* and miR390, the trigger miRNA for the production of functional tasiRNAs, are conserved amongst plant species, an artificial *TAS3* gene (*aTAS3*) doesn’t require the co-expression of a miRNA, reducing the experimental complexity. In *A. thaliana*, the endogenous *TAS3a* gene has been engineered to replace two or six native tasiRNAs with atasiRNAs targeting a gene of interest or the genome of a virus (Montgomery et al., 2008; Chen et al., 2016). Instead of modifying the endogenous *TAS3* gene by replacing specific atasiRNAs, a different approach has been successfully used in *A. thaliana*, tobacco and tomato to express atasiRNAs. It involves placing a miR390 cleavage site upstream and another downstream of the sequence targeted for silencing, in order to trigger the production of atasiRNAs from this sequence, for example a portion of viral genome (Felippes and Weigel, 2009; Singh et al., 2015; Singh et al., 2019).

Both endogenous and exogenous sRNA genes can be engineered to express asRNAs in a specific species, but they have to be experimentally tested to determine which strategy produces the highest asRNA levels. A *MIRNA* gene can efficiently express amiRNAs in its native host species but may not be as efficient in a different species. For example, it has been shown that the *ath-MIR159a* gene efficiently drives the expression of amiRNAs in *A. thaliana* but not in tobacco (Kung et al., 2012), the *ath-MIR319a* works efficiently in *A. thaliana* to express amiRNAs but it is not efficient in rice and the *osa-MIR528* gene works well in rice but not in *A. thaliana* (Li et al., 2013). These examples suggest that, to achieve gene silencing in a specific plant species, the miRNA precursor harboring the amiRNAs should be derived from the same species or from a closely related species, to ensure effective amiRNA processing and consequent gene silencing (Kung et al., 2012). These observations do not imply that endogenous miRNA precursors guarantee amiRNA expression, for example it has been found that in tomato plants the endogenous *sly-MIR159a* gene can drive amiRNA expression but another endogenous gene, *sly-MIR168a*, can’t (Vu et al., 2013). Furthermore, another work demonstrated that in *Brachypodium distachyon*, a chimeric *MIR390* precursor, in which the basal stem of the hairpin derives from *osa-MIR390* and the distal stem-loop derives from *ath-MIR390a*, produces higher levels of amiRNAs and is processed more accurately than the respective natural precursors (Carbonell et al., 2015).

Based on structural and sequence characteristics of natural *MIRNA* genes, there are general guidelines to design *aMIRNA* genes to increase their chance of being precisely processed into mature amiRNAs. To engineer a plant endogenous *MIRNA* gene, an important structural feature must be considered: the amiRNA* should be designed to mimic the bulges of the natural miRNA/miRNA* duplex, in order to retain the exact secondary structure of the natural miRNA precursor (Schwab et al., 2006). To design the sequence of the mature amiRNA, there are a few sequence features to follow that are determinants of plant miRNAs: 1) miRNAs tend to have a 5’ terminal uridine (U), that sort them into AGO1 complexes (Mi et al., 2008); 2) they have a bias for a cytosine at position 19 and an adenine at position 10 (Ossowski et al., 2008); 3) they have a GC content between 30 and 50% and display 5’ instability, which means higher adenine-uridine (AU) content at the 5’ and higher guanine-cytosine (GC) content at the 3’ (Ossowski et al., 2008) and 4) they have specific GC biases at positions 8-9 and 18-19 and specific AU biases at positions 5,7,10,15 (Narjala et al., 2020). Several computational methods can be used to design amiRNAs against RNAs of interest. These methods generally apply rules for target recognition and off-target avoidance and follow the above structural and sequence features, in order to identify the amiRNA candidates with the best potential for high accumulation and activity (Ossowski et al., 2008; Fahlgren et al., 2016; Mickiewicz et al., 2016). However, the sequence and structural determinants for precise processing of miRNAs are not fully understood and there are no known rules applicable to all miRNAs (Narjala et al., 2020), therefore *aMIRNA* candidates have to be tested in the specific plant system of interest to verify their efficacy.

The identification of efficient, precisely processed *asRNA* genes reduces the chances of off-targeting in the plant. An *asRNA* precursor that is not precisely processed could give rise to unwanted sRNA species that might recognize and down-regulate different RNAs other than the target. In order to assess this, high-throughput sequencing of sRNAs has been employed in plants expressing the *asRNA* genes. In contrast to Northern blot and stem-loop qRT-PCR, which are commonly used to detect the expression of the mature asRNAs, sRNA sequencing (sRNA-seq) allows measurements of the accumulation of all RNAs that are generated from an *asRNA* gene. Ideally, the majority of reads from the *asRNA* gene correspond to the correctly processed asRNA, for example in the case of the 21 nucleotide amiRNAs from the *ath-MIR390a*-based precursors in (Carbonell et al., 2014). In other cases, the *asRNA* gene produces reads corresponding to the desired asRNA but also additional sRNAs from other regions of the precursor that accumulate at high levels and have potential unintended gene targets, for example in the case of the *ath-MIR319a* precursor in petunia and the *vv-MIR319e* precursor in *N. benthamiana* (Guo et al., 2014; Castro et al., 2016).

In this work, we tested different polycistronic *asRNA* genes in tomato to determine their ability to be precisely and efficiently processed and therefore to be a good strategy for the simultaneous expression of multiple asRNAs. We screened five polycistronic *asRNA* genes by transiently expressing them in the leaves, to assess their activity in a faster way compared to the time-consuming generation of stably transformed plants. We analyzed the same *asRNA* genes in *N. benthamiana* as a control, because there is evidence that the amiRNA processing mechanisms operating in *Solanum* and *Nicotiana* are probably the same (Vu et al., 2013) and because *N. benthamiana* is easily transformed. As a case study, we decided to test the polycistronic *asRNA* genes by engineering them to express asRNAs designed to target genes involved in the drought stress response, because drought is amongst the most severe abiotic stresses affecting the productivity of most field crops including tomato. We report a detailed analysis of the expression and processing of four polycistronic *aMIRNA* genes and one *aTAS* gene in tomato and *N. benthamiana*, to identify the best working construct to express multiple asRNAs in these plant species.

## Results

### Design of monocistronic and polycistronic artificial microRNA and *trans*-acting siRNA genes

The mRNAs of eight tomato genes belonging to four gene families were chosen as targets of the amiRNAs (Table 1), because there is direct or indirect evidence in the literature that their downregulation enhances drought stress resistance in tomato. These genes encode for: protein phosphatase type 2Cs (*SlPP2C1*, *SlPP2C2*, *SlPP2C3*, (Gonzalez-Guzman et al., 2014), S-adenosylhomocysteine hydrolases (*SlSAHH1*, *SlSAHH2*, *SlSAHH3*, (Li et al., 2015), hybrid proline-rich protein 1 (*SlHyPRP1*, (Li et al., 2016) and ABA uridine diphosphate glucosyltransferase 75C1 (*SlUGT75C1*, (Sun et al., 2017).

**Table 1.**
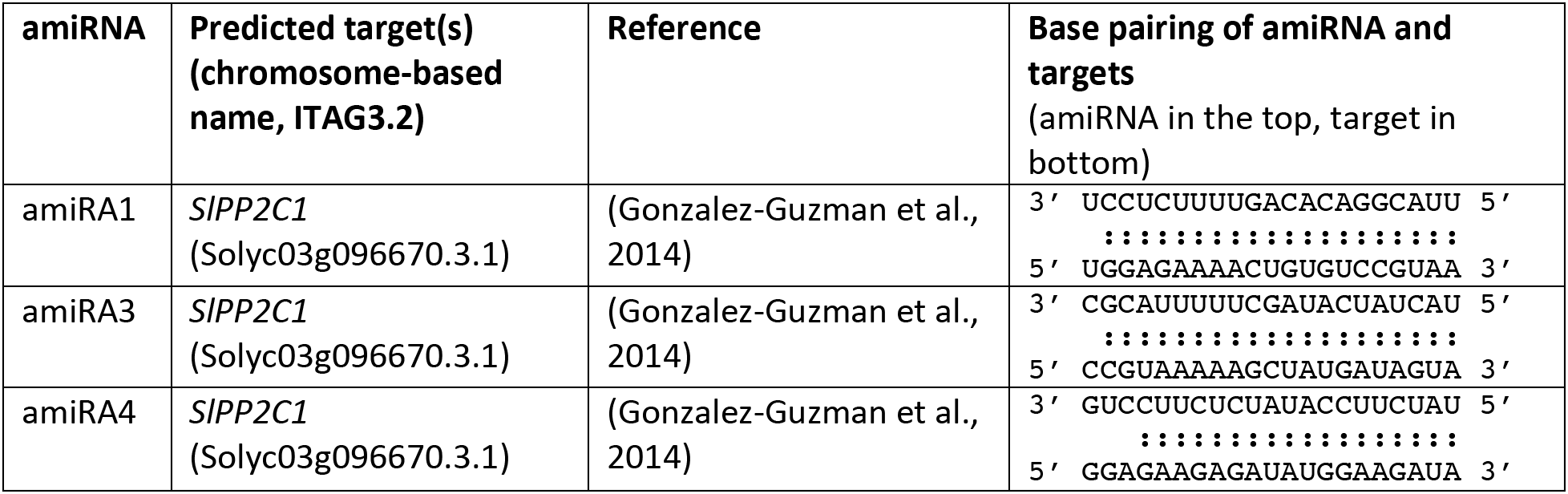
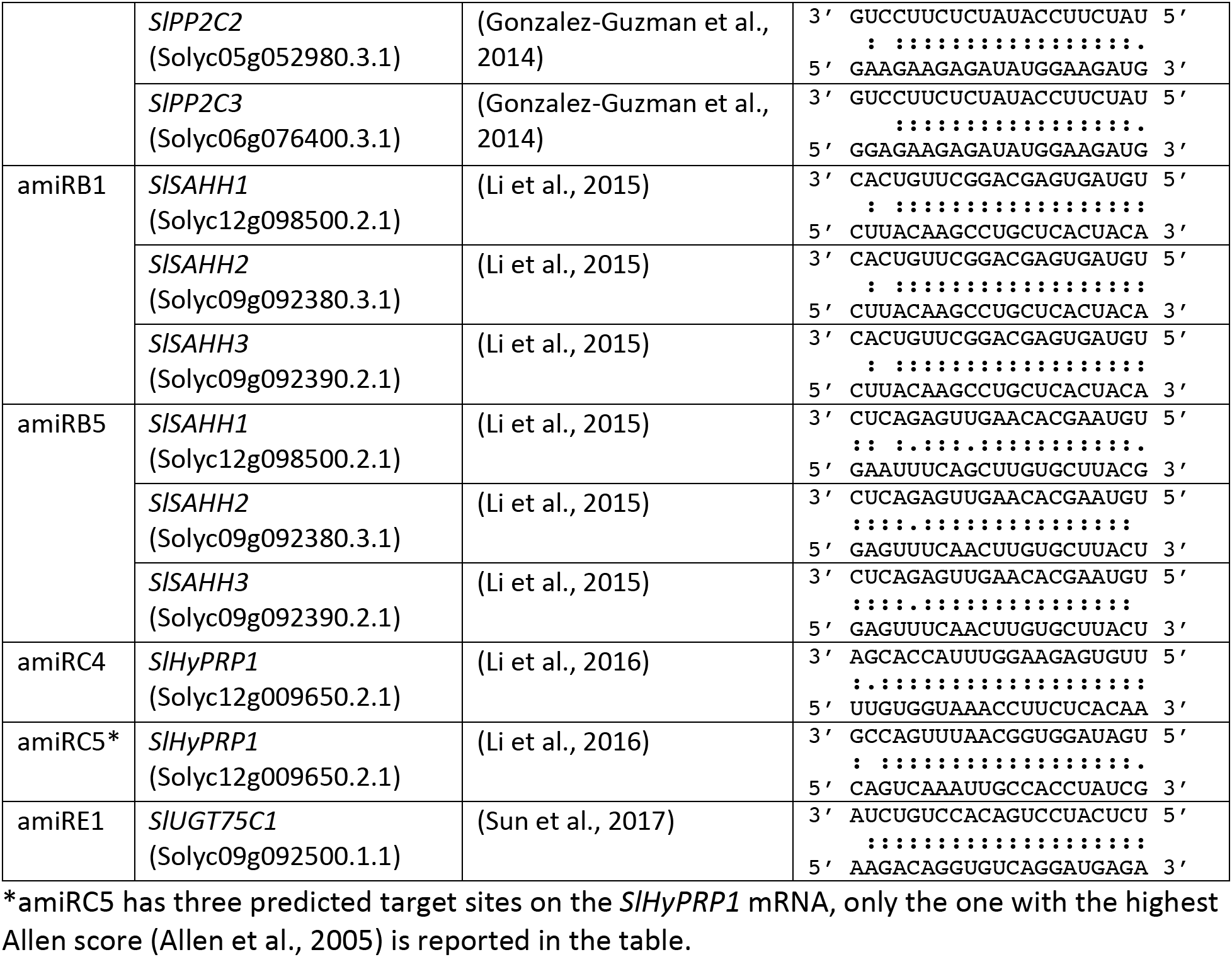
Base pairing of artificial microRNAs and target mRNAs.

Several amiRNAs were designed against the tomato target mRNAs using P-SAMS (Fahlgren et al., 2016) and WMD3 (Schwab et al., 2006) and the top eight candidates with the smallest number of possible off-target mRNAs in tomato were selected (Table 1). Each predicted amiRNA/mRNA base pairing was manually inspected to verify that it contained extensive complementarity, at most one mismatch between positions 2-13 and no mismatch between positions 9-11 (relative to the 5’ of the amiRNA), which are all critical features for plant miRNA-mediated target repression (Table 1, (Axtell, 2013a). Three amiRNAs were designed to target multiple mRNAs of the same gene family, by choosing a common region (amiRA4 against *SlPP2C1, SlPP2C2* and *SlPP2C3;* amiRB1 and amiRB5 against *SlSAHH1*, *SlSAHH2* and *SlSAHH3*), because of the possible functional redundancy of these genes in the response to drought stress (Gonzalez-Guzman et al., 2014; Li et al., 2015).

The *ath-MIR390a* hairpin was used to express each amiRNA individually, as monocistronic genes, in order to test their expression and provide a control for the expression of the polycistronic genes. This precursor was chosen because it has a short distal stem loop, is conserved amongst plants, it is processed accurately and it has been previously used in other species like *A. thaliana* and *N. benthamiana* to express 21-nt amiRNAs that can associate with AGO1 (Montgomery et al., 2008; Carbonell et al., 2014). The *ath-MIR390a* hairpins were designed according to (Carbonell et al., 2014), to contain one of the amiRNAs and its corresponding amiRNA* in place of the mature ath-miR390a/miR390a*, maintaining the pattern of matches and mismatches of the original *ath-MIR390a* (Figure 1A, Supplemental Figure 1, Supplemental Table 1).

**Figure 1.**
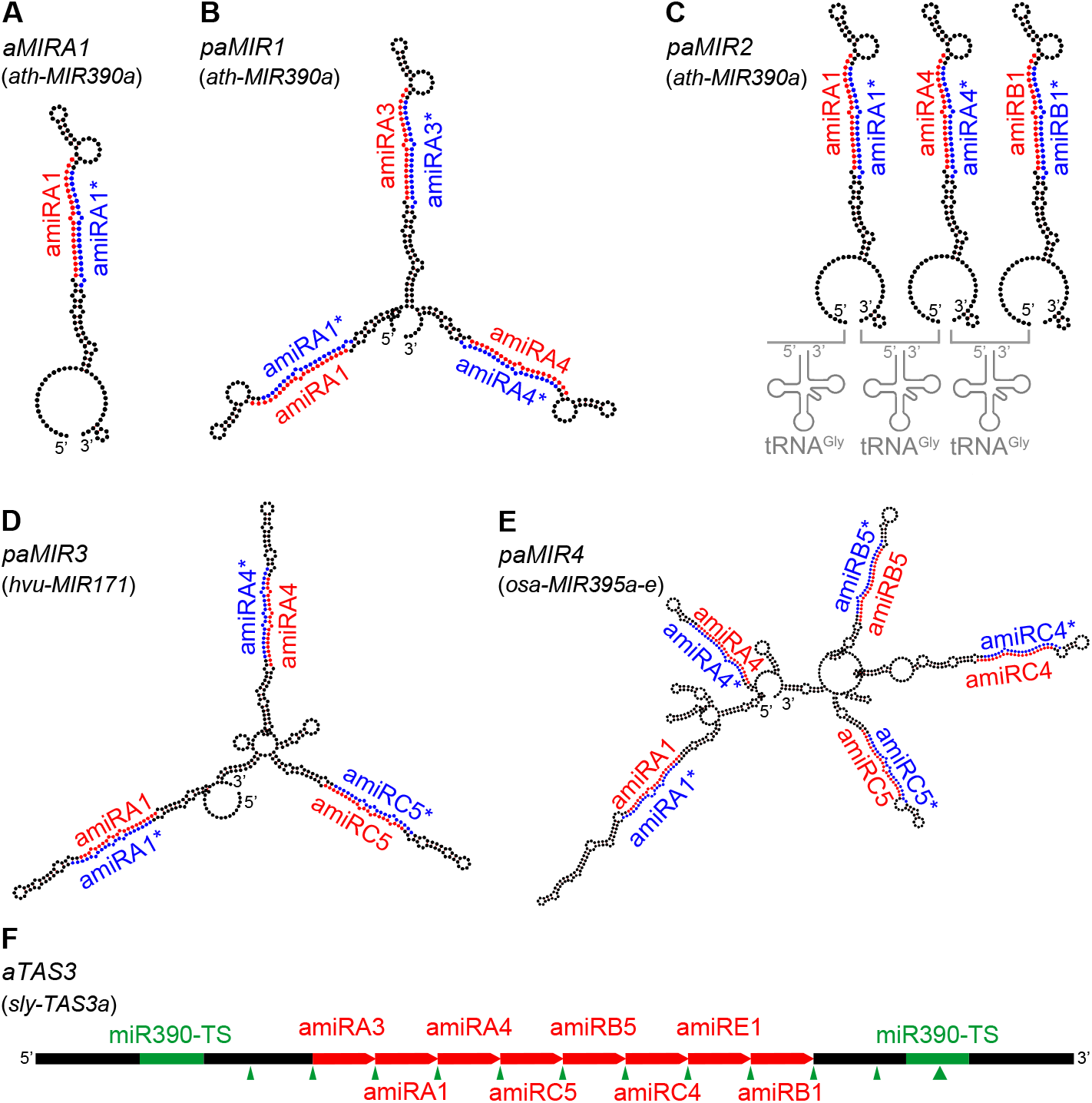
Monocistronic and polycistronic artificial microRNA and *trans*-acting siRNA genes. A-E) Predicted secondary structure of *aMIRNA* genes. A) *aMIRA1*: *ath-MIR390a* hairpin with amiRA1/amiRA1* in place of ath-miR390a/miR390a*; see Supplemental Figure 1 for the other monocistronic *aMIRNA* genes. B) *paMIR1*: tandem repeat of three *ath-MIR390a* hairpins. C) *paMIR2*: tandem repeat of three *ath-MIR390a* hairpins with the tRNA^Gly^ sequence upstream of each hairpin. D) *paMIR3*: tandem repeat of three *hvu-MIR171* hairpins. E) *paMIR4*: first five hairpins (a-e) of the rice endogenous polycistronic *osa-MIR395a-g* gene. Black: amiRNA backbone. Red: mature amiRNA. Blue: amiRNA*. Grey: secondary cloverleaf structure of tRNA^Gly^. F) Schematic of *aTAS3*: tomato endogenous *sly-TAS3a* gene with the endogenous tasiRNAs replaced by the amiRNA sequences. Black: *sly-TAS3a* gene. Green: miR390 target sites (mir390-TS). Thick green arrow: miR390-triggered cleavage site. Thin green arrows: positions where the *sly-TAS3a* transcript is subsequently cleaved in phase relative to the first cut site. Red: amiRNAs replacing the tasiRNAs.

Different strategies were tested to express the amiRNAs as polycistronic genes. The tomato genome was first analyzed to find endogenous polycistronic miRNA genes that could be engineered to express amiRNAs. Good candidates have two characteristics: 1) they are compact in size, smaller than 5kb, for easier in vitro manipulation and easier *Agrobacterium*-mediated in transient expression in the leaf; 2) they show evidence of accumulation of their mature miRNAs and they express uniquely mapping reads in tomato wild type sample, as a proof that their multiple hairpins are all successfully processed into miRNAs. No endogenous polycistronic miRNA gene with at least three hairpins was found to satisfy these features.

Five alternative strategies were chosen to express multiple amiRNAs as cluster of primary transcripts with a shared promoter. The first strategy, polycistronic *aMIR1* or *paMIR1*, was a tandem repeat of three *ath-MIR390a* hairpins, each carrying an amiRNA/amiRNA* duplex in place of the mature ath-miR390a/miR390a*, as described above (Figure 1B). The second strategy, *paMIR2*, was adapted from (Zhang et al., 2018) and consisted in placing a tRNA sequence upstream of each amiRNA hairpin in tandem. Specifically, a tRNA^Gly^ sequence was placed upstream of each of three *ath-MIR390a* hairpins in tandem, each carrying an amiRNA/amiRNA* duplex as described above (Figure 1C). The third strategy, *paMIR3*, was adapted from (Kis et al., 2016): a tandem repeat of three barley *hvu-MIR171* precursors, each carrying an amiRNA/amiRNA* duplex in place of the mature hvu-miR171/miR171* (Figure 1D). The fourth strategy, *paMIR4*, engineered the rice endogenous polycistronic miRNA gene *osa-MIR395a-g*, as in (Fahim et al., 2012). The first five hairpins of the gene were used and engineered to change the miR395/miR395* duplexes with the amiRNA/amiRNA* sequences (Figure 1E). After replacing the original miRNAs/miRNA*s with the amiRNAs/amiRNA*s in *paMIR1*, *paMIR2*, *paMIR3* and *paMIR4*, their backbone sequences were all modified when necessary to respect the same identical pattern of matches and mismatches of the original miRNA backbones. The last strategy, *aTAS3*, consisted in modifying the tomato endogenous *sly-TAS3a* gene, by replacing the endogenous tasiRNAs with the amiRNA sequences. Specifically, *sly-TAS3a* gene has 12 tasiRNA sequences that are generated between the two miR390 target sites, only the central eight tasiRNA sequences were replaced by amiRNAs (Figure 1F). Each artificial polycistronic sRNA gene contained a different combination of amiRNAs, targeting the same or different gene families (Figure 1, Supplemental Table 1).

### Processing precision of monocistronic and polycistronic artificial microRNA and *trans*-acting siRNA genes

All monocistronic and polycistronic asRNA genes were cloned into the Ti vector pLSU-1.1, a modified version of pLSU-1 (Lee et al., 2012), under the control of a short version of the 35S promoter and the NOS terminator (Supplemental Data 1). The Ti vectors were introduced into *A. tumefaciens* 1D1249, a strain that does not elicit necrosis when agroinfiltrated into tomato leaves (Wroblewski et al., 2005). Transient syringe agroinfiltration assays were performed in *N. benthamiana* and tomato leaves. For each construct, three replicates were obtained by agroinfiltrating three different leaves. *N. benthamiana* was studied for two reasons: i) to compare the results obtained in tomato with a different species that belongs to the same family (Solanaceae); ii) to serve as a control, because the transient agroinfiltration protocol is well established and works efficiently in *N. benthamiana*.

sRNA-seq was performed on each leaf sample, collected three days after the agroinfiltration (Supplemental Table 2). To determine how precisely the asRNA genes were processed to form mature asRNAs, the sRNA reads were mapped to the genes. The processing precision was calculated as the sum of reads aligned to the amiRNA and amiRNA*, divided by the number of reads mapping to the entire precursor (Figure 2). For comparison, the processing precision of 15 endogenous *MIRNA* genes was also calculated in control samples (leaves agroinfiltrated with the empty vector and non-agroinfiltrated leaves). Only two endogenous *MIRNA* genes had low levels of processing precision, *nbe-MIR156g* and *nbe-MIR169a*, likely due to a starting non-accurate annotation of their mature miRNA/miRNA* sequences. Amongst the monocistronic *aMIRNA* genes, *aMIRA1*, *aMIRA3*, *aMIRA4* and aMIRB5 showed the highest precisions in both tomato and *N. benthamiana*, with > 79% and > 89% of total precursor reads aligned to the amiRNA/amiRNA* duplex, respectively. *aMIRC5* and *aMIRE1* had slightly lower precisions, between 0.67-0.72 and 0.74-0.87 in tomato and *N. benthamiana*, respectively. *aMIRB1* and *aMIRC4* were the two genes with the lowest values: in both species, their processing precision was lower than the majority of endogenous *MIRNA* genes. These results suggest that the same amiRNA precursor can be processed with different levels of precision, depending on the sequence of the amiRNA/amiRNA* duplex.

**Figure 2.**
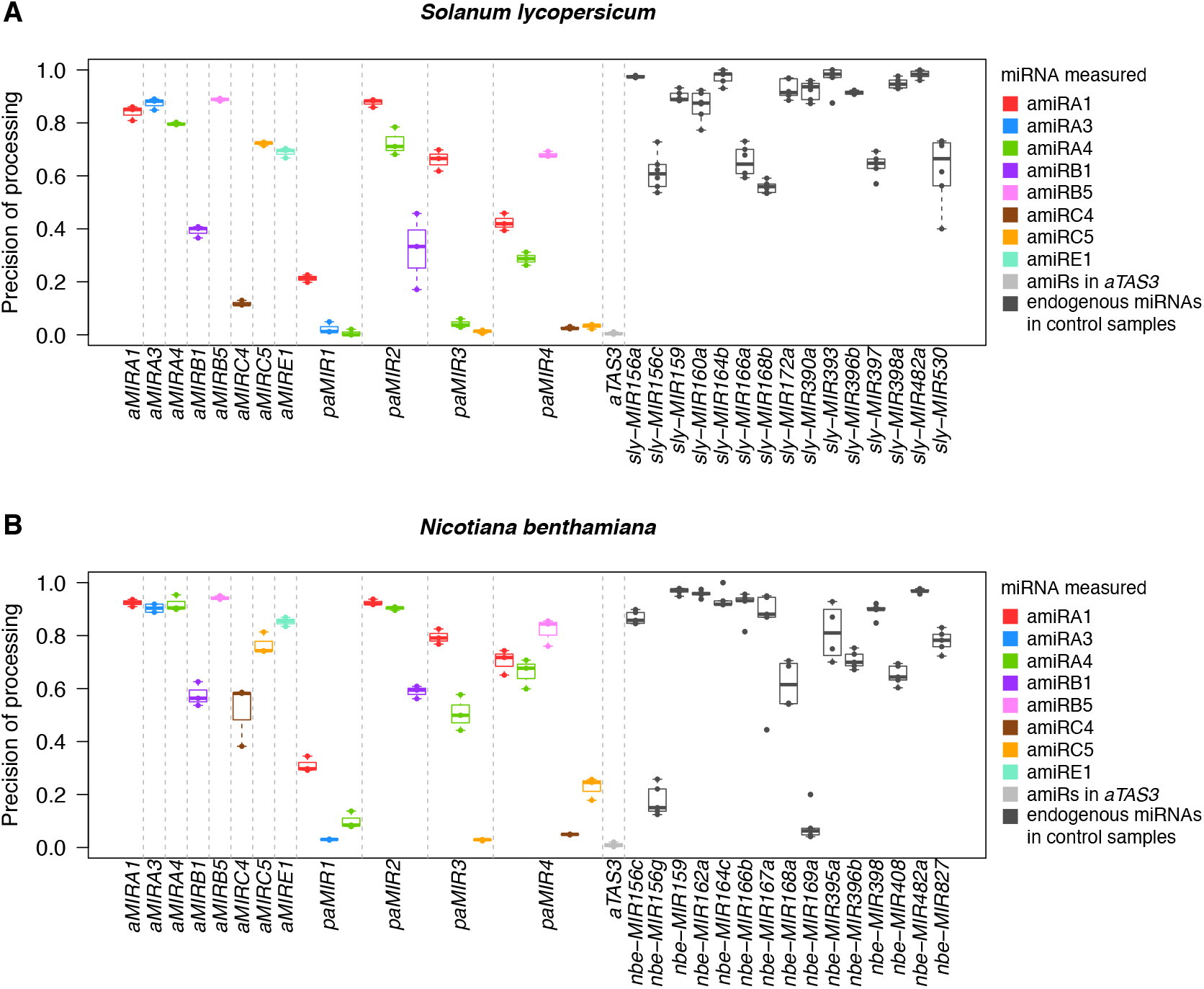
Precision of processing of the artificial microRNA and *trans*-acting siRNA genes. See Supplemental Code 1 for generating these plots. A) Precision of processing in tomato, defined as: [reads aligned to miRNA + reads aligned to miRNA*] / [total reads aligned to miRNA hairpin]. 0: none of the reads mapped to the precursor corresponds to the mature miRNA or miRNA*. 1: all the reads mapped to the precursor align to the miRNA/miRNA* duplex. The x axis shows which *asRNA* gene was agroinfiltrated. The color corresponds to the specific amiRNA measured with sRNA-seq. Light grey indicates the processing precision of *aTAS3*, considering all its amiRNA sequences together. Dark grey indicates the processing precision of a set of 15 endogenous *MIRNA* genes, measured in leaflets agroinfiltrated with the empty vector and in non-agroinfiltrated leaflets. B) Precision of processing in *N. benthamiana*.

A general trend was observed amongst the polycistronic *aMIRNA* genes in both species: the processing precision of the hairpin precursors was higher for those located at the 5’ end of the gene and decreased for those located at the 3’ end (Figure 2). In contrast, the *aTAS3* gene had very few reads aligned to the designed amiRNAs (replacing the endogenous tasiRNAs), independently of their position in the precursor, resulting in a total processing precision < 0.02 in both species. Both *paMIR1* and *paMIR2* are composed by three *ath-MIR390a* hairpins in tandem, therefore the processing precision of their hairpins can be directly compared with the monocistronic *aMIRNA* genes carrying the same amiRNA/amiRNA* duplex. In *paMIR1*, all the hairpins were processed less accurately into mature amiRNAs compared to their monocistronic version. In contrast, in *paMIR2*, the three hairpins containing amiRA1, amiRA4 and amiRB1 were processed with similar precisions of their corresponding monocistronic genes. These results were reproducible in tomato and *N. benthamiana*, suggesting that placing three *ath-MIR390a* hairpins in tandem under the same promoter reduced the precision at which the hairpins were processed and that the addition of tRNA^Gly^ upstream of each hairpin increased the processing precision, restoring the level of the individual *aMIRNA* genes (Zhang et al., 2018). In *paMIR3*, only the hairpin at the 5’ end of the gene had > 50% of reads aligned to the amiRNA/amiRNA* duplex. The low processing precision of the middle and 3’ end hairpins indicates that *paMIR3*, developed in the monocot species barley and based on the *hvu-MIR171* precursor (Kis et al., 2016), is not an efficient candidate polycistronic *aMIRNA* gene for the two dicot species studied. The last polycistronic *aMIRNA* tested, *paMIR4*, was also developed in a monocot species, wheat, and was based on the rice *MIR395* polycistronic gene (Fahim et al., 2012). In *N. benthamiana*, *paMIR4* showed high processing precision for the first three hairpins at the 5’ and low for the two hairpins at the 3’ end. In tomato, the relative processing precision of the five hairpins was similar to *N. benthamiana* but only the *aMIRB5* hairpin was > 0.5, suggesting that this polycistronic *aMIRNA* strategy is a better candidate for *N. benthamiana* expression than tomato.

In *N. benthamiana* the processing precision scores were generally higher than in tomato, but the relative distribution of these values between the *aMIRNA* genes was similar in both species (Figure 2). These results indicate that the different polycistronic amiRNA strategies tested behaved similarly in the two Solanaceae species studied but with higher efficiency in *N. benthamiana*.

### sRNA expression profile of monocistronic and polycistronic artificial microRNA and *trans*-acting siRNA genes

The profile of sRNA expression from the entire length of the *aMIRNA* and *aTAS* genes was investigated to further understand how efficiently the different precursors were processed in the two species. The sRNA coverage of the monocistronic *aMIRNA* genes was plotted as a stacked histogram based on the length of the sequenced sRNAs, at a single nucleotide resolution (Figure 3). sRNA-seq data from the three replicates were merged because they were highly consistent (Supplemental Figure 2). The sRNA coverage of each *aMIRNA* gene was measured in the leaves agroinfiltrated with the same amiRNA construct. Control non-agroinfiltrated leaves showed no sRNA reads mapping to any of the *aMIRNA* genes (Supplemental Figure 2). Both in tomato and *N. benthamiana*, *aMIRA1*, *aMIRA3*, *amiRA4* and *aMIRB5* showed specific expression of sRNAs from the amiRNA/amiRNA* duplex position of the hairpin and the vast majority of the reads corresponded to the mature amiRNA (Figure 3). Compared to these precursors, *aMIRC5* and *aMIRE1* were also precisely processed into mature amiRNAs but with higher counts of amiRNA* reads, which in *aMIRE1* were more abundant than the mature amiRNA. Both *aMIRB1* and *aMIRC4* precursors were processed heterogeneously, with no specific expression of the mature amiRNA. However, they both showed accumulation of reads in the amiRNA* position, which was more evident in *N. benthamiana* than tomato for amiRC4. These results confirm the previous observation (Figure 2) that the same amiRNA precursor can be processed differently depending on the sequence of the amiRNA/amiRNA* duplex.

**Figure 3.**
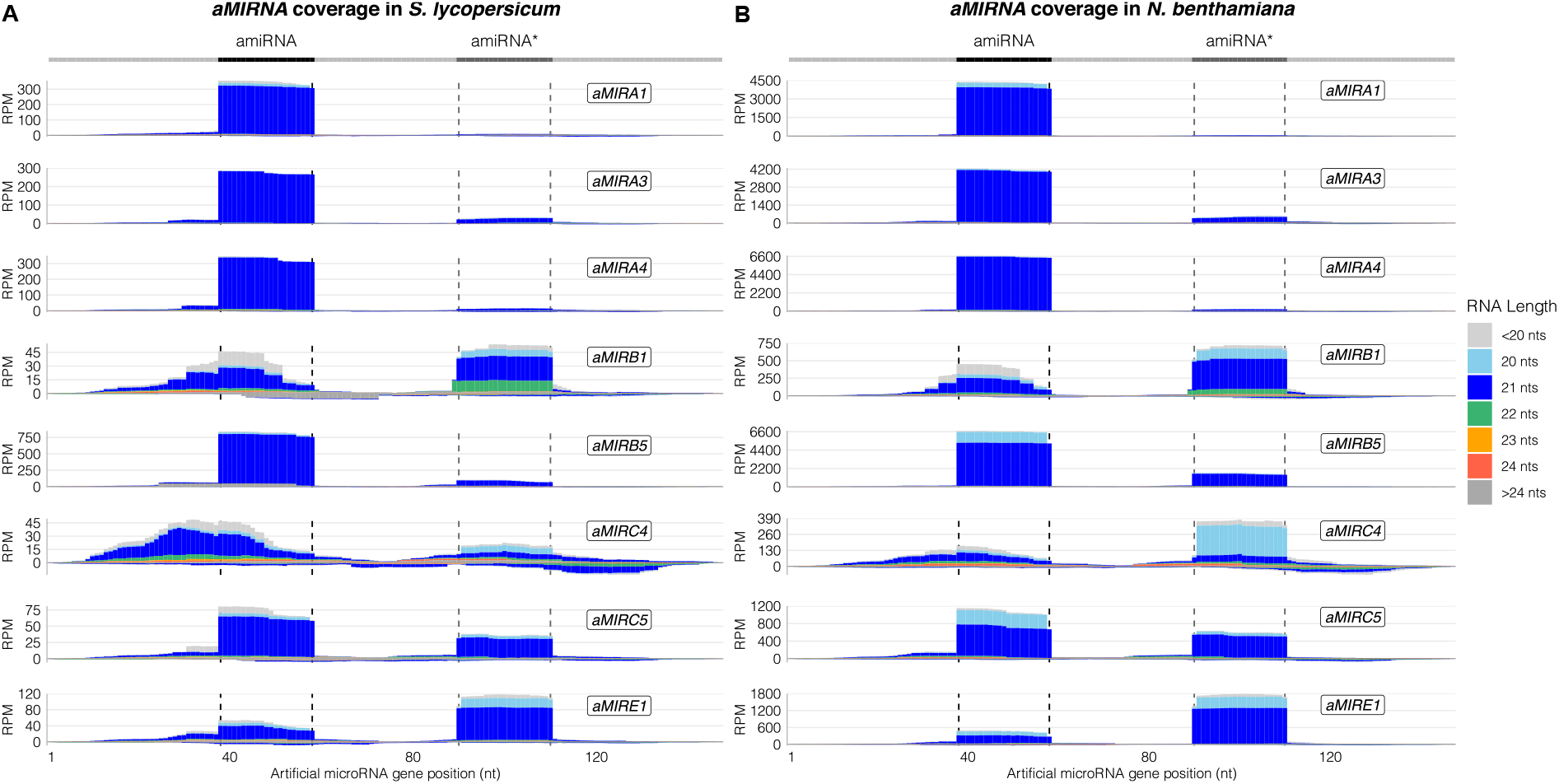
sRNA coverage of the artificial monocistronic microRNA genes. See Supplemental Code 2 for generating this plot. A) Tomato sRNA coverage across each *aMIRNA* gene. The x axis indicates the position on the gene in nucleotides, from 5’ to 3’. At the top, the light grey line corresponds to the precursors’ backbone; the position of the amiRNA and amiRNA* in the precursors are indicated in black and dark grey respectively. The y axis is the sRNA coverage in Reads Per Million (RPM) for each nucleotide. Positive coverage means the reads align to the + DNA strand, negative coverage means the reads align to the - DNA strand. The coverage of reads with different lengths is represented separately with different colors, stacked from bottom to top according to the legend on the right. Coverages of all reads shorter than 20 nucleotides and longer than 24 nucleotides are represented in light grey and dark grey respectively. B) *N. benthamiana* sRNA coverage across each *aMIRNA* gene.

The amiRNAs were screened for a few sequence features that are important for increasing the chance of precise processing of miRNA precursors (Schwab et al., 2006; Mi et al., 2008; Ossowski et al., 2008; Narjala et al., 2020). All the amiRNA sequences had a uridine at the 5’ position, a cytosine at position 19, and GC content between 30-50% (Supplemental Table 3). Amongst the other sequence features, none were specifically biased in amiRB1, amiRC4, and amiRE1 in ways that could explain why these amiRNAs were processed less precisely than the others (Supplemental Table 3).

To determine if the accumulation of sRNA reads in the amiRNA and amiRNA* positions of the hairpins (Figure 3) corresponded exactly to the mature amiRNA positions, the coverage of the 5’ nucleotide of each read was plotted at the 5’ of the amiRNAs and amiRNA*s and five nucleotides upstream and downstream (Supplemental Figure 3). This close up view of the sRNA coverage confirmed that the *aMIRA1*, *aMIRA3*, *aMIRA4*, *aMIRB5*, *aMIRC5* and *aMIRE1* hairpins were processed exactly at the predicted 5’ of both the amiRNAs and amiRNA*s, while in *aMIRB1* only the amiRNA* had significant accumulation at the exact predicted position. Finally, the two *aMIRNA* genes that had the lowest processing precisions, *aMIRB1* and *aMIRC4*, and the two *aMIRNA* genes with the highest proportions of amiRNA* reads, *aMIRC5* and *aMIRE1*, all had in common the accumulation, at different levels, of a 20 nucleotide sRNA processed one nucleotide downstream of the amiRNA* 5’ end.

As a control, the sRNA coverage and sRNA 5’ coverage plots were also generated for the same set of endogenous *MIRNA* genes of Figure 2 (Supplemental Figure 4, Supplemental Figure 5). The sRNA profiles of the endogenous *MIRNA* genes showed an enriched accumulation of the predicted mature miRNA, confirming that amongst the tested *aMIRNA* genes, those that had sRNA profiles similar to the endogenous *MIRNA* genes (*aMIRA1*, *aMIRA3*, *aMIRA4*, *aMIRB5*, *aMIRC5* and *aMIRE1*) are the best candidates for amiRNA expression.

The sRNA expression profile was then analyzed for the polycistronic *aMIRNA* and *aTAS* genes. In *paMIR1*, both tomato and *N. benthamiana* showed accumulation of reads corresponding to the mature amiRA1 (Figure 4, Supplemental Figure 6). In contrast, both the *aMIRA3* and *aMIRA4* hairpins showed expression of reads outside the amiRNA/amiRNA* duplex, with some differences between tomato and *N. benthamiana*. This suggests that the two hairpins at the 3’ end of *paMIR1* are processed differently from the prediction, possibly because in vivo they don’t form the predicted hairpin secondary structures (Figure 1B). The sRNA 5’ coverage plot revealed that the *aMIRA1* hairpin was also not processed exactly as expected, because the most abundant 21 nucleotide sRNA originated from one nucleotide upstream amiRA1 (Figure 5).

**Figure 4.**
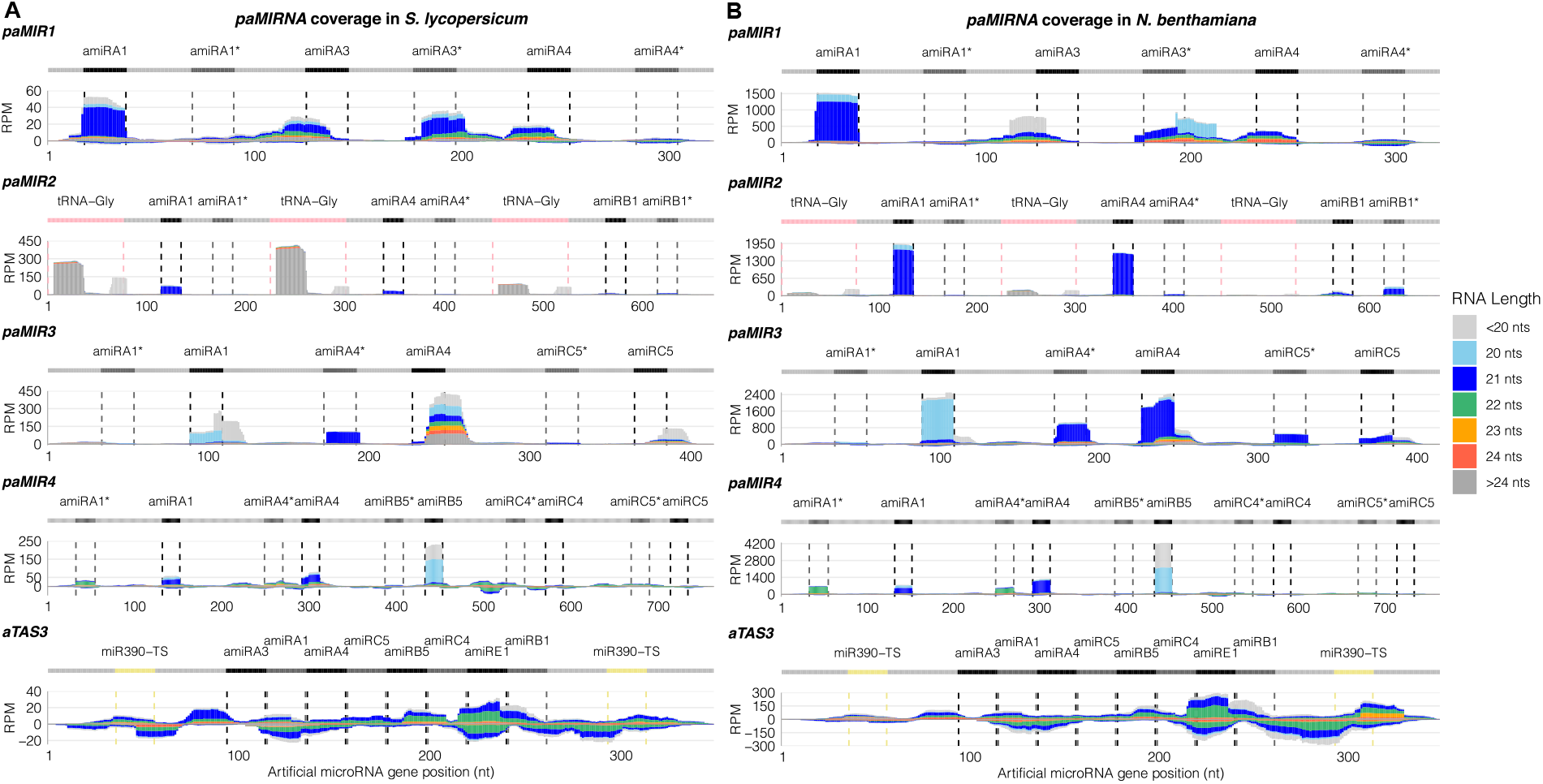
sRNA coverage of the artificial polycistronic microRNA and *trans*-acting siRNA genes. See Supplemental Code 3 for generating this plot. A) Tomato sRNA coverage across each polycistronic *aMIRNA,* or *paMIRNA*, and *aTAS* gene. The x axis indicates the position on the gene in nucleotides, from 5’ to 3’. At the top of each plot, the light grey line corresponds to the precursor backbone; the position of the amiRNA and amiRNA* in the precursors are indicated in black and dark grey respectively, except for *aTAS3* where these colors are used to distinguish adjacent amiRNAs; pink indicates the tRNA^Gly^ sequences; yellow indicates the miR390 target sites. The y axis is the sRNA coverage in RPM for each nucleotide. Positive coverage means the reads align to the + DNA strand, negative coverage means the reads align to the - DNA strand. The coverage of reads with different lengths is represented separately with different colors, stacked from bottom to top according to the legend on the right. Coverages of all reads shorter than 20 nucleotides and longer than 24 nucleotides are represented in light grey and dark grey respectively. B) *N. benthamiana* sRNA coverage across each *aMIRNA* and *aTAS* gene.

**Figure 5.**
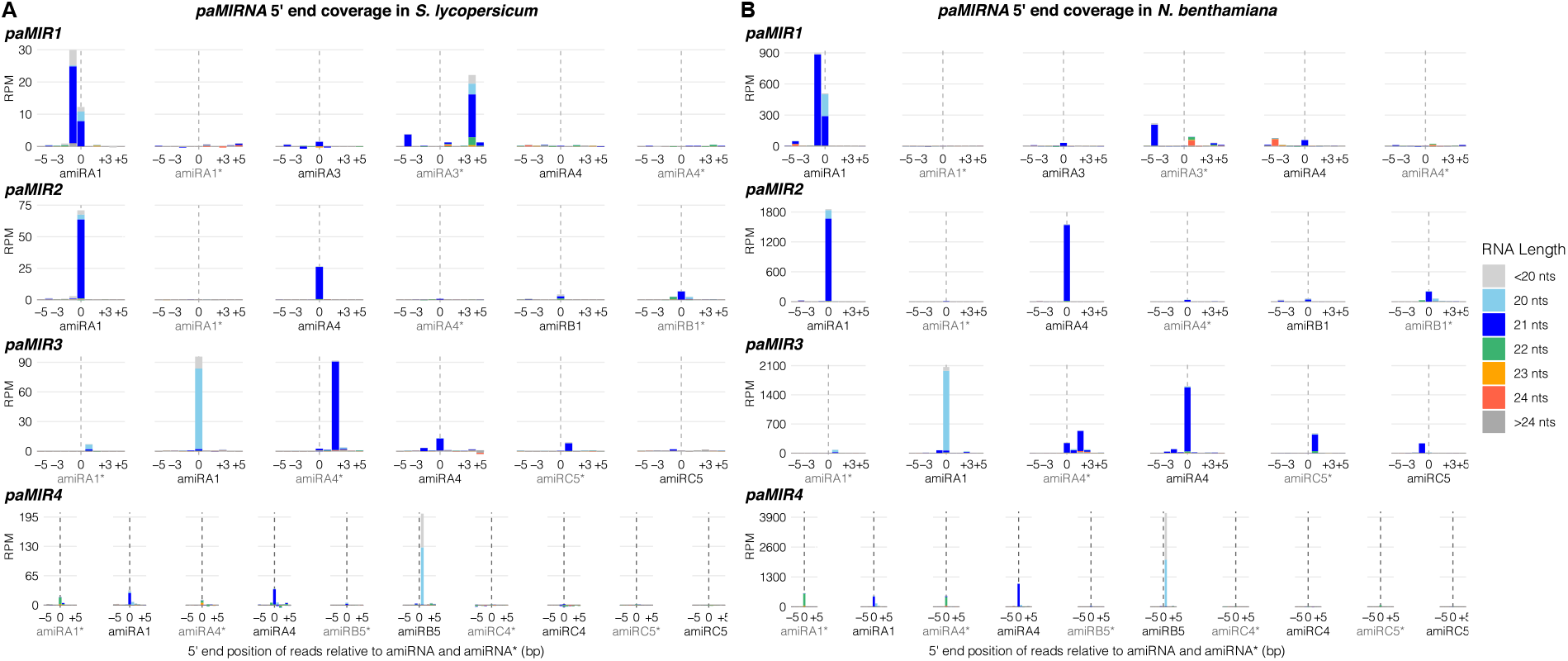
sRNA 5’ coverage around the artificial miRNA and miRNA* 5’ ends in the artificial polycistronic microRNA genes. See Supplemental Code 4 for generating this plot. A) Tomato sRNA 5’ coverage. In the x axis, 0 indicates the 5’ end of the amiRNAs and amiRNA*s, −5 and +5 indicate five nucleotides upstream and downstream of them. The y axis is the sRNA 5’ coverage in RPM. Positive coverage means the reads align to the + DNA strand, negative coverage means the reads align to the - DNA strand. The sRNA 5’ coverage of reads with different lengths is represented separately with different colors, stacked from bottom to top according to the legend on the right. Coverages of all reads shorter than 20 nucleotides and longer than 24 nucleotides are represented in light grey and dark grey respectively. B) *N. benthamiana* sRNA 5’ coverage.

The sRNA coverage of the three hairpins of *paMIR2* (Figure 4, Supplemental Figure 6) corresponded to the sRNA profile of the same hairpins expressed as monocistronic genes (Figure 3): *aMIRA1* and *aMIRA4* generated predominantly the mature amiRA1 and amiRA4 sequences respectively, instead *aMIRB1* was only accurately processed at the amiRB1*. The sRNA 5’ coverage profile of *paMIR2* (Figure 5) confirmed that this polycistronic amiRNA construct was processed as its corresponding individual hairpins. For example, *aMIRB1* showed similar accumulation of a 22 nucleotide sRNA one nucleotide upstream of amiRB1* and of a 20 nucleotide sRNA one nucleotide downstream of amiRB1* in both the monocistronic *aMIRB1* gene (Supplemental Figure 3) and the polycistronic *paMIR2* gene (Figure 5). This result indicates that the presence of a tRNA sequence upstream of each hairpin enhances the *in vivo* processing precision of the polycistronic RNA into single hairpins (Zhang et al., 2018) also in tomato and *N. benthamiana*. Finally, reads outside the 20-24 nucleotides range that aligned to two regions of the tRNA^Gly^ (Figure 4) were likely derived from genomic tRNA sequences, because they were also observed in the control, non-agroinfiltrated leaves in both species (Supplemental Figure 6).

The sRNA coverage of *paMIR3* was different between tomato and *N. benthamiana* (Figure 4, Supplemental Figure 6). In both species *paMIR3* generated many sequences outside the 20-24 nucleotides range, but in tomato the fraction of these reads was higher than *N. benthamiana*. In both species, the hairpin at the 5’ of the gene generated a 20 nucleotides version of the mature amiRA1 instead of the predicted 21 nucleotides, one nucleotide shorter at the 3’ (Figure 5). The hairpin in the center of the gene was correctly processed into the mature amiRA4 and amiRA4* sequences in *N. benthamiana*, with amiRA4 more abundant than amiRA4*. Instead in tomato, the same hairpin was processed into a mix of different sRNA sizes partially overlapping amiRA4 (Figure 4). The mature amiRA1, amiRA4 and amiRC5 were all highly precisely processed in the monocistronic genes with the *ath-MIR390* background (Figure 3) and in *paMIR3* they were all inserted in the same *hvu-MIR171* hairpin, suggesting similar processing precision in *paMIR3*. In contrast, in both species the hairpin at the 3’ of *paMIR3* was the only one where both the amiRNA and amiRNA* sequences (amiRC5/amiRC5*) were generated one nucleotide away from their predicted 5’ ends (Figure 5). This confirmed the previous observation that the processing precision of the polycistronic *aMIRNA* genes is lower towards the 3’ of the genes.

Amongst the five hairpins of *paMIR4*, only the first three at the 5’ of the gene expressed sRNAs corresponding to the amiRNAs and amiRNA*s in both species (Figure 4, Supplemental Figure 6). This was another example of the difference in processing precision between hairpins located at the 5’ and at 3’ of a polycistronic *aMIRNA* gene. The difference between the two species was that in tomato there were more reads coming from regions of the hairpins outside the amiRNA/amiRNA* duplex compared to *N. benthamiana*. The two hairpins at the 5’ of *paMIR4* were both processed like the rice *MIR395a* and *MIR395b* into 21 nucleotides mature amiRNAs and 22 nucleotides amiRNA*s. Instead in the center of *paMIR4*, amiRB5 was diced one nucleotide downstream the annotated 5’ of osa-miR395c, into highly abundant 20 nucleotides and shorter RNAs (Figure 5).

In both species, *aTAS3* generated a mix of sRNAs of different sizes that did not correspond to the amiRNA sequences that in *aTAS3* replaced the endogenous tasiRNAs of *sly-TAS3a* (Figure 4, Supplemental Figure 6). The reads mapped to *aTAS3* were not even in phase, a defining feature of *TAS* genes. For example, in tomato the phase score of *TAS3a* and *TAS3b* is 541 and 344.6, respectively, according to the data analyzed in plantsmallrnagenes.science.psu.edu (Lunardon et al., 2020). Using the same calculator (Axtell, 2013b), the phase score of *aTAS3* in tomato was between 6.9 and 9.7 in three replicates. The *aTAS3* construct is therefore not a good strategy for the expression of multiple artificial sRNAs in tomato and *N. benthamiana*.

Finally, a general difference was observed between tomato and *N. benthamiana*, both for the monocistronic and the polycistronic *aMIRNA* genes. In *N. benthamiana*, in addition to higher processing precision scores (Figure 2), higher expression levels were also measured for the *aMIRNA* genes compared to tomato (Figure 3, Figure 4), suggesting that the *Agrobacterium*-mediated transient expression of amiRNAs is more efficient in *N. benthamiana*.

### Comparison of artificial microRNA levels measured through sRNA sequencing and stem-loop qRT-PCR

In order to confirm the abundance of the mature amiRNAs detected with sRNA sequencing, stem-loop qRT-PCR was performed on the same RNA samples of agroinfiltrated leaves that were used for sequencing (Supplemental Figure 7). For each amiRNA, its accumulation level measured through sRNA-seq was plotted versus its accumulation level measured through stem-loop qRT-PCR (Figure 6). Only amiRNAs that were detected with both techniques were reported in the plot. In the majority of cases, when an amiRNA was not reported it was because it was not sequenced (RPM = 0) and the stem-loop qRT-PCR did not detect it or produced a false positive (Supplemental Figure 7). Only in a few cases, an amiRNA was not sequenced but had a valid low stem-loop qRT-PCR signal or vice versa, it was not detected in stem-loop qRT-PCR but was sequenced at low RPMs, indicating possible false positives or very lowly abundant RNAs. In both species, there was a positive correlation between the two techniques, meaning that in general, the stem-loop qRT-PCR confirmed the sRNA-seq data (Figure 6). This positive correlation was not perfect, and many biases were observed, for example amiRC4 and amiRC5 from the monocistronic genes were overestimated in the stem-loop qRT-PCR or underestimated in sRNA-seq. Theses biases were conserved in the two species, that showed similar relative distributions, and therefore depended on the specific sequence of the measured RNA. If examining only a single target RNA sequence, the correlation between stem-loop qRT-PCR and sRNA sequencing was noticeably consistent. This indicates that RNA-sequence-specific effects likely made a significant contribution to the discrepancies.

**Figure 6.**
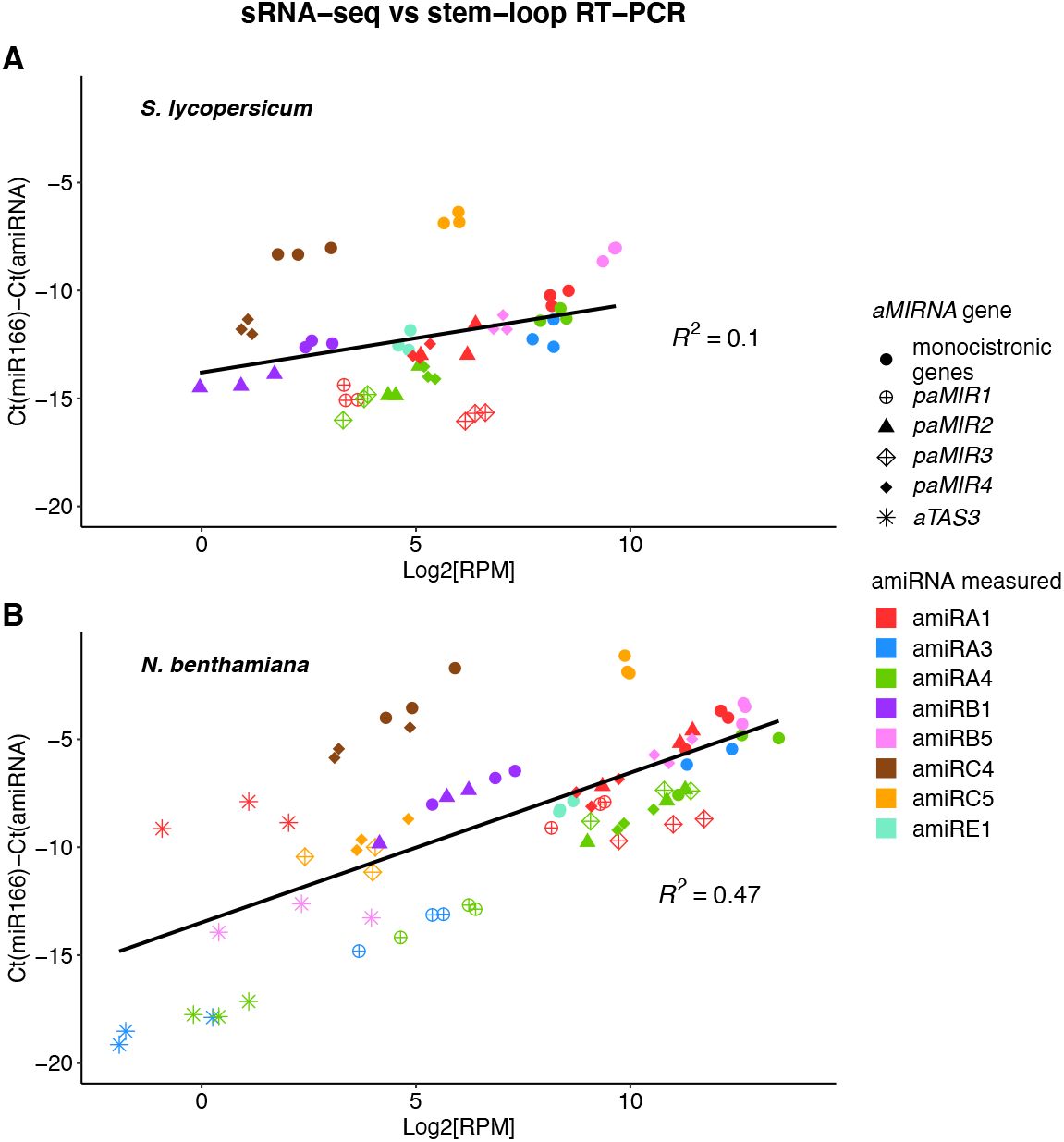
Correlation between sRNA-seq and stem-loop qRT-PCR artificial microRNA levels. See Supplemental Code 5 for generating this plot. A) Correlation in tomato. The x axis shows the abundance of the mature amiRNAs measured through sRNA-seq, as the binary logarithm of their RPM value. The y axis shows the abundance of the mature amiRNAs measured through stem-loop qRT-PCR, as the difference between the cycle threshold (Ct) of the loading control miR166 and the amiRNA. The color of the data points indicates the mature amiRNA sequence that was measured, the shape indicates the sample in which it was measured. Data points with same color and shape are replicates. B) Correlation in *N. benthamiana*.

The main difference between these two amiRNA quantification methods was that a number of amiRNAs that were detected at very low levels in sRNA-seq compared to other amiRNAs, in stem-loop qRT-PCR they showed similar abundance levels compared to other amiRNAs. These amiRNAs were: amiRB1 and amiRC4 from the monocistronic genes, amiRB1 from *paMIR2*, amiRA4 from *paMIR3* in tomato, amiRC5 from *paMIR3* in *N. benthamiana*, amiRC4 and amiRC5 from *paMIR4* and all the sequences from *aTAS3* (Figure 6). These amiRNAs corresponded to those hairpins with low processing precisions measured by sRNA-seq (Figure 2). One possibility is that their accumulation was overestimated in stem-loop qRT-PCR; the alternative possibility is that sRNA-seq under-estimated the true accumulations. Both methods rely on steps that can introduce sequence-specific bias including, reverse transcription, PCR, and specifically for sRNA-seq, RNA ligation.

### Target gene accumulation in tomato

The amiRNAs were designed to target specific genes in tomato (Table 1), therefore the accumulation of the target genes was investigated in tomato but not in *N. benthamiana*. To minimize possible gene expression variations dependent on specific leaves and leaflets, the same tomato leaflet was agroinfiltrated with an *aMIRNA* or *aTAS* gene on the left side and with the control empty vector on the right side and the gene accumulation was compared between the two sides. Multiple replicates were performed in different leaflets, leaves and plants. The accumulation of the gene mRNAs was measured with qRT-PCR (Figure 7). Overall, there was no correlation between the accumulation of the amiRNAs and the accumulation of their target genes. For example, in *paMIR2*-agroinfiltrated leaflets, amiRA1 and amiRA4 were detected at higher levels than amiRB1 according to sRNA-seq data (Figure 5) but their target genes all showed no significant up- or down-regulation (Figure 7).

**Figure 7.**
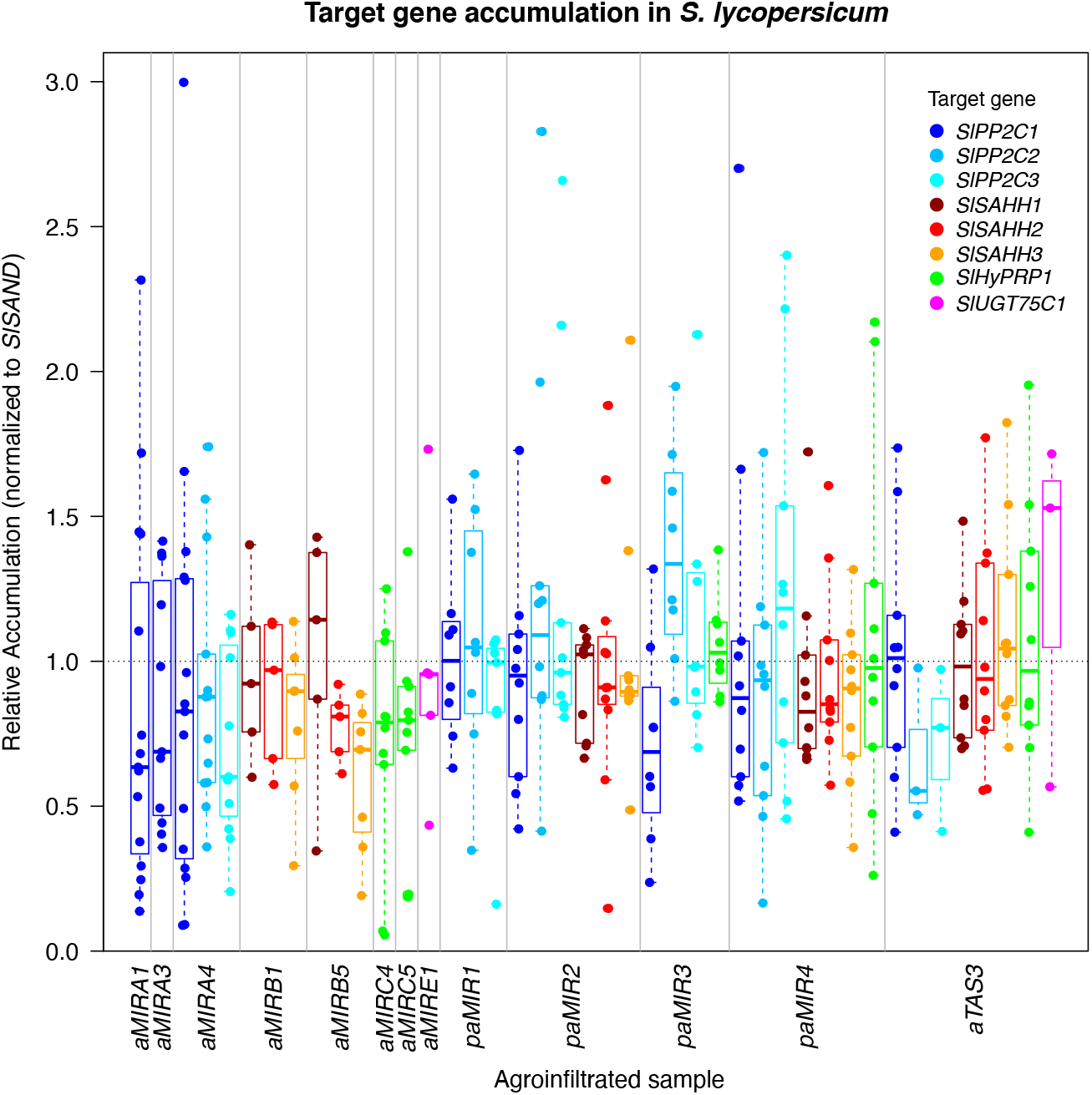
qRT-PCR detection of artificial miRNA targets in tomato. See Supplemental Code 6 for generating this plot. The y axis shows the accumulation of the target genes in the side of the tomato leaflet agroinfiltrated with an *aMIRNA* or *aTAS* construct, relative to their accumulation level in the control side of the leaflet, agroinfiltrated with the empty vector construct. Values < 1 mean down-regulation and values > 1 mean up-regulation of the mRNAs. Accumulation data are normalized on the reference gene *SlSAND*. The x axis shows in which sample the data are measured and the color of the data points which mRNA is measured. Different data points in the same column are replicates.

The vast majority of target genes didn’t show any consistent effect as a consequence of the amiRNA accumulation, with a few exceptions. Amongst the polycistronic *aMIRNA* genes, *paMIR3* produced the highest abundant amiRNA processed with a precise 5’ end: a one nucleotide shorter version of amiRA1 (70-96 RPM; Figure 5), which was associated with down-regulation of its target *SlPP2C1* in six out of eight total replicates (Figure 7). Amongst the monocistronic *aMIRNA* genes, *aMIRB5* produced the amiRNA with the highest accumulation level (638-784 RPM, Figure 3) and its targets *SlSAHH2* and *SlSAHH3* showed down-regulation in all replicates of *aMIRB5* agroinfiltrations (Figure 7). This suggests that there might be a threshold of amiRNA expression to reach in *Agrobacterium*-mediated transient assays in order to detect significant effects.

## Discussion

Our analysis of polycistronic *aMIRNA* and *atasiRNA* processing revealed several surprising results. The use of small RNA-sequencing, as opposed to single-target assays such as RNA blots or stem-loop qRT-PCR, was critical in revealing deficiencies of various approaches. We were surprised, and dismayed, that so many of the approaches failed to accumulate precisely processed miRNAs or siRNAs. Previous studies have focused on amiRNA-target mRNA interactions as a key determinant of function (Deveson et al., 2013; Li et al., 2013). Our data suggest that another confounding variable is the precise (or imprecise) processing of precursors. The same *aMIRNA* hairpin backbone could have very large variation in the precision of processing depending only on the sequences of the amiRNA and corresponding amiRNA*. This occurred despite the fact that the overall secondary structures were maintained during design. This variation also was not obviously correlated with previously described sequence features (Schwab et al., 2006; Mi et al., 2008; Ossowski et al., 2008; Narjala et al., 2020) that are known to affect plant *MIRNA* processing.

There are multiple possible reasons for variability in *aMIRNA* processing: A) The hairpin secondary structure actually formed *in vivo* might be different than the predicted secondary structure, and negatively affect processing. This has been suggested as an explanation for the failure of engineered *sly-MIR168a* in tomato (Vu et al., 2013). B) The tertiary hairpin RNA structure might be defective and thus interfere with the processing of the hairpins forming the complex RNA structure; C) Unknown, yet important, sequence determinants of *MIRNA* hairpin processing could have been inadvertently altered by the insertion of the amiRNA and amiRNA* sequences. Our study did not examine AGO loading, although all amiRNAs were designed to possess a 5’-U, which is required for loading into the major miRNA and siRNA AGO, AGO1 (Mi et al., 2008). It is possible that there are other, as-yet discovered sequence contexts that encourage or discourage AGO loading. In this scenario, poorly loaded amiRNAs would be expected to accumulate to lower levels because of more rapid turnover.

Our attempts to use the *aTAS3* strategy were wholly unsuccessful in *N. benthamiana* and tomato. sRNA sequencing showed that the *aTAS3* RNA was not processed as expected. The same strategy was previously successful in *Arabidopsis*, where the atasiRNAs were successfully detected and the mutants showed a phenotype (Chen et al., 2016). Perhaps alternative *TAS3-* based strategies, such as using only the miR390 target sites and excluding any other *Arabidopsis*-derived sequences (Singh et al., 2015; Singh et al., 2019) may improve *aTAS* performance in the Solanaceae.

One noteworthy finding was that the processing precision for most polycistronic *aMIRNA* hairpin precursors was higher for the 5’-most hairpins and decreased for the 3’-most hairpins. Interestingly, this contrasts with observations of polycistronic *aMIRNA* processing in human cells, where the position of the hairpins made no difference (Hu et al., 2010). One possible explanation for this polar effect in plants, but not animals, could be the contrasting methods of hairpin processing. Plant *MIRNA* hairpins are processed directly into miRNA/miRNA* duplexes by a Pol-II associated complex involving Dicer-Like 1 (Fang and Spector, 2007; Fang et al., 2015). The coupling between hairpin transcription and DCL1 processing could be rapid enough that the 5’-most hairpins of a polycistronic RNA are diced before the 3’ hairpins are transcribed. This in turn could destabilize the nascent transcript in much the same way as typical Pol II transcription is terminated by cleavage and polyadenylation. In contrast, animals use Drosha to produce short hairpins that are then exported to the cytoplasm for final processing into miRNA/miRNA* duplexes; this may be less directly coupled to transcription and thus not trigger premature termination of the polycistronic precursor. Clearly direct experiments are required in the future to test this hypothesis. Nonetheless, the polar effect of plant *aMIRNA* polycistronic precursors is plainly evident from our data.

Overall the best approach in our system for producing multiple functional amiRNAs (transient expression in *N. benthamiana* and tomato) were the tRNA-based constructs (‘*paMIR2’;* Figure 1C; (Zhang et al., 2018). This judgement is based primarily on the higher processing precisions observed by small RNA-sequencing (Figure 4). Despite this, *paMIR2* was not particularly effective at actually silencing its target mRNAs in tomato (Figure 7). This is a stark reminder that the efficacy of any particular amiRNA *in vivo* remains difficult to predict *a priori*. We see three explanations for this. A) Even with precise processing, the resulting amiRNAs may not be efficiently loaded into AGO proteins. If there are as yet unknown determinants for miRNA/AGO association, certain amiRNAs may not function well. B) It is now well-established that local mRNA secondary structure at and surrounding miRNA target sites can exert strong effects on miRNA function (Li et al., 2014; Yang et al., 2020). However neither the prediction of *in vivo* local mRNA secondary structures, nor the quantitative details of structure’s effects on miRNA function, are robust enough to make this easily predictable; thus target site selection remains hit or miss. C) Plant miRNAs can sometimes exert much stronger effects on protein accumulation as opposed to mRNA accumulation; this is often attributed to translational repression occurring independently of mRNA degradation. In our experiments we were not able to directly assess protein accumulation of our native tomato proteins. D) In these experiments, the amiRNAs were expressed transiently through *Agrobacterium* leaf infiltrations. Compared to stable transgenic lines, transient expression assays might be associated with higher expression variability, which might have caused high variability on the accumulation of the target genes.

This study highlights the need to consider variable precursor processing as a confounding variable when using amiRNAs or asiRNAs. These difficulties in maturation become especially pronounced when attempting multiple gene silencing. The use of small RNA sequencing, which captures all small RNA species, as opposed to single-target assays such as RNA blots or real-time PCR, is critical to detect these problems. Future studies to discern exactly what mechanisms explain highly variable precursor processing would be most useful for improving multiplex gene silencing by amiRNAs.

## Methods

### Design and selection of artificial microRNAs

An initial set of amiRNA candidates were designed for each tomato mRNA target, with the online tools: P-SAMS (Fahlgren et al., 2016), allowing the software to automatically filter the results based on target specificity, and WMD3 (Schwab et al., 2006), with default parameters. The top three candidates from each software were then tested for possible off-targets using the sRNA-transcriptome aligner GSTAR (https://github.com/MikeAxtell/GSTAr) (GSTAr.pl -t -a *amiRNA_input_file.fasta tomato_transcriptome_input_file.fasta* > *output_file.txt*). The top amiRNA sequences with the smallest number of possible off-target mRNAs with Allen score < 3 were selected.

### Analysis of tomato endogenous polycistronic microRNA genes

The tomato *MIRNA* gene annotations for the genome version v2.5 from Ensembl Plants (Cunningham et al., 2019) were retrieved from 1) the tomato gene annotation file v2.5 from Ensembl Plants (Cunningham et al., 2019), selecting for ‘pre_miRNA’ feature type; 2) the tomato small RNA locus annotation file from https://plantsmallrnagenes.science.psu.edu (Lunardon et al., 2020), selecting for ‘MIRNA’ and ‘nearMIRNA’ feature types. The above annotations were merged, the duplicates were manually removed and the unique *MIRNA* genes were clustered with the BEDTools function ‘bedtools cluster’ (Quinlan and Hall, 2010) to find *MIRNA* genes closer than 5kb (bedtools cluster -i *input_file.gff3* -d 5000 > *output_file.gff3*). Clustered *MIRNA* genes were manually screened in the Plant small RNA genes genome browser (Lunardon et al., 2020) for predominant RNA size produced, expression level and proportion of uniquely aligned reads.

### Annotation of endogenous miRNAs and miRNA*s in tomato and *N. benthamiana*

Fifteen endogenous *MIRNA* genes were manually annotated for their miRNA and miRNA* sequences in the tomato genome version SL4.0 and the *N. benthamiana* genome version v1.0.1 from The Sol Genomics Network (Fernandez-Pozo et al., 2015). *MIRNA* precursors were retrieved from the miRBase22.1 database (Kozomara and Griffiths-Jones, 2014) for tomato (SL2.50 genome version) and *N. tabacum* (genome version not available). Their homologous sequences in the tomato and *N. benthamiana* genome versions mentioned above were found with BLAST (Camacho et al., 2009) using the best results. Fifteen *MIRNA* genes belonging to conserved, highly confident miRNA families were then randomly selected for each species. Their secondary structures were predicted with mfold (Zuker, 2003) and were used to manually annotate miRNAs and miRNA*s, following the 2 nucleotides 3’ overhangs criteria for plant miRNA/miRNA* duplex annotation (Axtell and Meyers, 2018). The coordinates of the *MIRNA* genes were finally extended of a variable number of nucleotides so that in all *MIRNA* genes 35 bases upstream and 35 bases downstream of the miRNA/miRNA* duplex were included, consistently with the amiRNA genes.

### Design of monocistronic and polycistronic artificial sRNA genes

The final selected amiRNA sequences were separately introduced in the *ath-MIR390a* precursor in place of miR390a, designing the amiRNA* according to (Carbonell et al., 2014). Compared to (Carbonell et al., 2014), the sequence of *ath-MIR390a* was extended, including 20 extra nucleotides both at the 5’ and 3’ of the precursor from the genomic sequence of *ath-MIR390a*. See Supplemental Table 1 for the complete sequences of the monocistronic *aMIRNA* genes. The DNA sequences of the single *aMIRNA* genes were chemically synthesized by GenScript® (GenParts™ Synthesis service) in linearized, double-stranded DNA (dsDNA) fragments.

The *paMIR1* gene was designed by adding in tandem three *ath-MIR390a* sequences, without the extra flanking 20 nucleotides, containing the amiRA1, amiRA3 and amiRA4 mature sequences. The *paMIR2* gene was designed by adding in tandem three *ath-MIR390a* sequences, with the extra flanking 20 nucleotides, containing the amiRA1, amiRA4 and amiRB1 mature sequences, and introducing upstream of each precursor the tRNA^Gly^ sequence, according to (Zhang et al., 2018). The *paMIR3* gene was designed as a tandem repeat of three *hvu-MIR171* precursors, where the mature miR171s were replaced with amiRA1, amiRA4 and amiRC5 and the amiRNA*s were designed according to (Kis et al., 2016). Following the strategy developed by (Fahim et al., 2012), the *paMIR4* gene consisted of a truncated version of the *osa-MIR395a-g* backbone, containing the first five miRNA hairpins. The miR395 sequences were replaced with amiRA1, amiRA4, amiRB5, amiRC4 and amiRC5 and the amiRNA*s were designed to mimic the pattern of matches and mismatches of the original miR395/miR395* duplexes. 20 nucleotides were added both upstream of the first hairpin and downstream of the last hairpin from the genomic sequence of *osa-MIR395a-e*. The *aTAS3* gene was designed starting from the tomato endogenous *TAS3a* gene, including the sequence between the two miR390 target sites and 35 extra nucleotides both upstream of the 5’ target site and downstream of the 3’ target site. The endogenous tasiRNA sequences in the positions 5’D3(+), 5’D4(+), 5’D5(+), 5’D6(+), 5’D7(+), 5’D8(+), 5’D9(+) and 5’D10(+) were replaced with amiRB1, amiRE1, amiRC4, amiRB5, amiRC5, amiRA4, amiRA1 and amiRA3, respectively. See Supplemental Table 1 for the complete sequences of the polycistronic *asRNA* genes. The DNA sequences of the polycistronic *asRNA* genes were chemically synthesized and inserted into the pMA-RQ or pMA-T plasmids by Invitrogen by Thermo Fisher Scientific (GeneArt Gene Synthesis service). Predicted secondary structures were obtained with RNAfold (option “view in forna”, http://rna.tbi.univie.ac.at/cgi-bin/RNAWebSuite/RNAfold.cgi).

### Cloning of artificial sRNA genes into pLSU-1.1 and transformation into *A. tumefaciens* 1D1249

A modified version of the binary Ti vector pLSU-1 (Lee et al., 2012), with a 315bp 35S promoter, multi cloning site and NOS terminator, here called pLSU-1.1, was kindly provided by Dr. Wayne Curtis (The Pennsylvania State University, PA, USA) (see Supplemental Data 1 for the complete sequence of the pLSU-1.1 plasmid). Some *asRNA* genes were amplified before thee cloning to introduce the correct restriction enzyme sites, using Q5® High-Fidelity DNA Polymerase (see Supplemental Table 4 for primers). The PCR products were purified with the E.Z.N.A.® Cycle Pure Kit V-spin (Omega Bio-tek). The purified PCR products and the pLSU-1.1 plasmid were digested with KpnI-HF® and BamHI-HF® (New England Biolabs^®^ Inc.). With the same restriction enzymes, the other *asRNA* genes were directly digested from the pMA-RQ/pMA-T plasmids. The digestion products were run on agarose gel and purified with Monarch^®^ DNA Gel Extraction Kit (New England Biolabs^®^ Inc.). The *asRNA* genes were ligated into pLSU-1.1 with T4 DNA Ligase (New England Biolabs^®^ Inc.) and the ligation products were transformed into *Escherichia coli* for colony PCR screening. Positive clones were further confirmed with Sanger sequencing and transformed by electroporation into *A. tumefaciens* strain 1D1249, kindly provided by Dr. Gregory B. Martin (Boyce Thompson Institute for Plant Research, NY, USA), because it does not elicit necrosis when agroinfiltrated into tomato leaves (Wroblewski et al., 2005). Colony PCR was used to find *A. tumefaciens* 1D1249 positive transformants, further confirmed by Sanger sequencing of the purified colony PCR products.

### Leaf transient agroinfiltrations and RNA extraction

Seeds of *S. lycopersicum* variety Lanai were kindly provided by Dr. Wayne Curtis (The Pennsylvania State University, PA, USA). Tomato Lanai and *N. benthamiana* were grown in a growth room at 22°C, 16 h light 8 h dark regime. Tomato plants with seven or eight leaves and *N. benthamiana* plants with five to nine leaves were used for transient agroinfiltrations for sRNA-seq. Additional tomato plants with five to nine leaves were agroinfiltrated for the analysis of tomato target mRNA levels. At day one, *A. tumefaciens* 1D1249 was inoculated overnight at 28°C in 3mL of YEP broth with kanamycin 50μL/mL. At day two, 2μL of the overnight culture were inoculated in 15mL YEP broth with kanamycin 50μL/mL, MES 10mM and acetosyringone 20μM. At day three, the bacteria were precipitated by centrifugation at 3,000 rpm for 6 minutes and resuspended in the inoculation buffer, containing MgCl_2_ 10mM, MES 10mM and acetosyringone 100μM, to different final concentrations: OD_600_ of 0.2 for agroinfiltration of tomato and *N. benthamiana* leaves used in sRNA-seq, and OD_600_ ranging from 0.2 to 1 for agroinfiltration of additional tomato plants for the analysis of target mRNA levels. The bacteria were left on the bench for 3 hours and then were agroinfiltrated with a 1mL syringe on the underside of the leaves. For sRNA-seq, three biological replicates were obtained for each *asRNA* gene, by agroinfiltrating the same bacteria into three separate primary leaflets in the 3^rd^ and 4^th^ compound leaves of the same tomato plant, and into three separate leaves in different *N. benthamiana* plants. For the analysis of tomato target mRNA levels, the same tomato leaflet was agroinfiltrated with bacteria carrying an *asRNA* gene on the left side of the leaflet and with bacteria carrying the control empty vector on the right side. At day six, three days after the agroinfiltrations were performed, the areas of the leaves that were agroinfiltrated were cut with a scalpel blade and flash frozen in liquid nitrogen, stored at −80°C, and then ground in the Mini-Beadbeater-16 (BioSpec Products). Total RNA was extracted with TRI Reagent (Sigma-Aldrich) per manufacturer instructions, adding a second sodium-acetate–ethanol precipitation and ethanol wash step.

### sRNA sequencing

The sRNA-seq libraries were prepared using dual indexing for multiplexing, according to (Persson et al., 2017). The following modification was applied: instead of using a pre-adenylated 3’ adapter, a non-adenylated 3’ adapter was ordered for chemical synthesis from IDT™ and adenylated with the 5’ DNA Adenylation Kit (New England Biolabs^®^ Inc.) (see Supplemental Table 4 for primers and adapters used). For the library PCR amplification step, a unique combination of forward and reverse primers was used for each sample. The four forward primers used were taken from (Persson et al., 2017). The 24 reverse primers used were the NEBNext Index 1-12 Primers for Illumina from NEBNext^®^ Multiplex Oligos for Illumina^®^ (Index Primers Set 1, E7335) and NEBNext Index 13-16, 18-23, 25 and 27 Primers for Illumina from NEBNext^®^ Multiplex Oligos for Illumina^®^ (Index Primers Set 2, E7500). A small aliquot for each library was run on a 6% acrylamide gel to verify the presence of the expected bands at 159-162bp corresponding to sRNA insert sizes of 21-24 nucleotides. All the samples were pooled together, mixing different volumes of the individual amplified libraries based on the intensity of their 159bp band on gel: 1μL for libraries with darker bands and 1.3μL for libraries with lighter bands. The pooled PCR reactions were purified with the Monarch^®^ PCR & DNA Cleanup Kit (5 μg) (New England Biolabs^®^ Inc.) and run on 6% acrylamide gel. A band between approximately 147-180bp, corresponding to RNA insert sizes of 9-42 nucleotides, was extracted, the gel pulverized and mixed with 3 volumes of Elution Buffer (300mM sodium acetate and 1mM EDTA pH8) and eluted overnight in a rotator at room temperature. The pooled libraries were then ethanol-precipitated and resuspended in TE buffer. Extracted bands were quantified by qPCR and quality-controlled by high-sensitivity DNA chip (Agilent). Sequencing was performed on a NextSeq (Illumina) in single read, 75 nucleotides in length, run mode by the Penn State genomics core.

### sRNA-seq data processing

The 3ʹ adapter was removed from the sRNA-seq FASTQ files (Supplemental Table 2) with cutadapt (Martin, 2011) (cutadapt -a *3_adapter_sequence* --discard-untrimmed -m 15 -o *output_file.fastq input_file.fastq*). The quality of reads was checked with FastQC (http://www.bioinformatics.babraham.ac.uk/projects/fastqc). The reads were aligned to the *asRNA* genes with ShortStack v3.8.1 (Axtell, 2013b; Johnson et al., 2016) (ShortStack --align_only --keep_quals --readfile *input.fastq* --outdir *output_directory* --genomefile *asRNAgene_sequence.fa*). The reads were also aligned to the genome sequences of *S. lycopersicum* SL4.0 and *N. benthamiana* v1.0.1, downloaded from The Sol Genomics Network (Fernandez-Pozo et al., 2015), with ShortStack v3.8.1 with default parameters.

### sRNA precision of processing and coverage

The precision of processing was defined as: [reads aligned to miRNA + reads aligned to miRNA*] / [total reads aligned to miRNA hairpin], both for amiRNAs and endogenous miRNAs. Reads mapped to the miRNA and miRNA* positions were extracted from the alignment files with the BEDTools function ‘intersectBed’, restricting for reads wholly contained inside the miRNA and miRNA* positions and derived from the same strand of the miRNA (Quinlan and Hall, 2010) (intersectBed -wao -F 1 -s -a *miRNA_and_miRNA*_coordinates.gff3* -b *input.bam* > *output.bam*). Only reads with length between 20 and 24 nucleotides long were considered in this analysis.

To obtain the sRNA coverage on the *asRNA* genes, a custom Perl script was developed that uses as input an alignment file (Supplemental Code 7). To obtain the sRNA 5’ end coverage around the amiRNA and amiRNA* start positions, the alignment files were first selected for mapped reads with the SAMtools function ‘samtools view’ (Li et al., 2009) (samtools view -b -F 4 *input.bam* > *mapped.bam*) and then converted to BED format with the BEDTools function ‘bamtobed’ (Quinlan and Hall, 2010) (bedtools bamtobed -i *mapped.bam* > *mapped.bed*). The 5’ end position of each read was extracted with a custom Perl script and the sRNA 5’ end coverage was then obtained with the same script used above (Supplemental Code 7).

### Stem-loop quantitative real time PCR (qRT-PCR) of artificial microRNAs and real time qRT-PCR of target genes

The stem-loop qRT-PCRs were performed according to (Varkonyi-Gasic et al., 2007), starting from total RNA samples that were all diluted to the same final concentration of 125ng/μL, and using the ProtoScript^®^ II Reverse Transcriptase (New England Biolabs^®^ Inc.) for the pulsed reverse transcription. miR166 was used as internal control (see Supplemental Table 4 for primer sequences). The amiRNA accumulation was calculated by subtracting the C_t_ value of the amiRNA of interest from the C_t_ value of miR166. False positives were amplification products with wrong band size in a 4% agarose gel electrophoresis and they were considered as undetected.

To perform qRT-PCR, total RNA samples were first treated with DNase I (RNase-free New England Biolabs^®^ Inc.), per the manufacturer’s instructions, ethanol precipitated and resuspended. The treated total RNA (1 μg) was used for cDNA synthesis with the MultiScribe™ Reverse Transcriptase (Thermo Fisher Scientific) according to the manufacturer’s instructions. PCR reactions used PerfeCTa^®^ SYBR^®^ Green FastMix^®^ (Quantabio) on a StepONE-Plus quantitative PCR system (Applied Biosystems) per the manufacturer’s instructions. Specific primers were designed using Primer BLAST (https://www.ncbi.nlm.nih.gov/tools/primer-blast/) or were selected from published papers (Supplemental Table 4). The housekeeping gene *SlSAND* was used as internal control (Expósito-Rodríguez et al., 2008). The relative accumulation of mRNAs was calculated as: 2^(dC_ta_-dC_tc_), where dC_ta_ = (C_t_^*SlSAND*^ – C_t_^gene of interest^), measured in dthe left side of the leaflet transiently expressing an artificial sRNA gene, and dC_tc_ = (C_t_^*SlSAND*^ – C_t_^gene of interest^), measured in the right part of the leaflet agroinfiltrated with the control empty vector.

## Accession numbers

The sRNA-seq data were deposited in the SRA database with the accession numbers reported in Supplemental Table 2.

## Acknowledgements

We thank Dr. Wayne Curtis (The Pennsylvania State University, PA, USA) for kindly providing the modified pLSU-1 vector, and Dr. Gregory B. Martin (Boyce Thompson Institute for Plant Research, NY, USA) for kindly providing *A. tumefaciens* strain 1D1249. This work was supported by an award from the United States Defense Advanced Research Projects Agency (Agreement Number HR0011-17-2-0055).

## Supplemental Figures

**Supplemental Figure 1.**
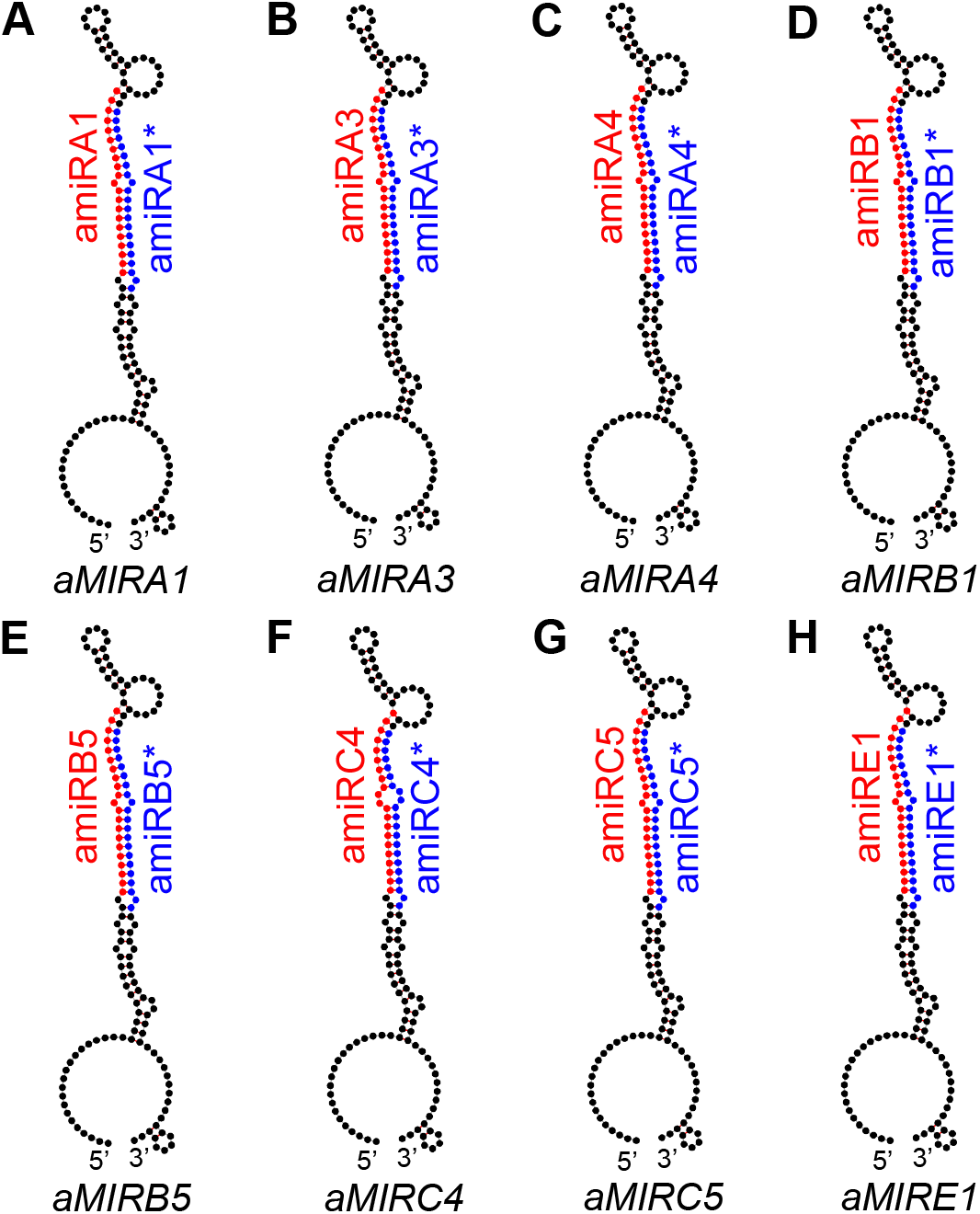
Predicted secondary structure of the monocistronic artificial microRNA genes. A) *aMIRA1*: *ath-MIR390a* hairpin with amiRA1/amiRA1* in place of ath-miR390a/miR390a*. B-H) Same structure of A) but for amiRA3/amiRA3*, amiRA4/amiRA4*, amiRB1/amiRB1*, amiRB5/amiRB5*, amiRC4/amiRC4*, amiRC5/amiRC5* and amiRE1/amiRE1*. Black: *aMIRNA* backbone. Red: mature amiRNA. Blue: amiRNA*.

**Supplemental Figure 2.**
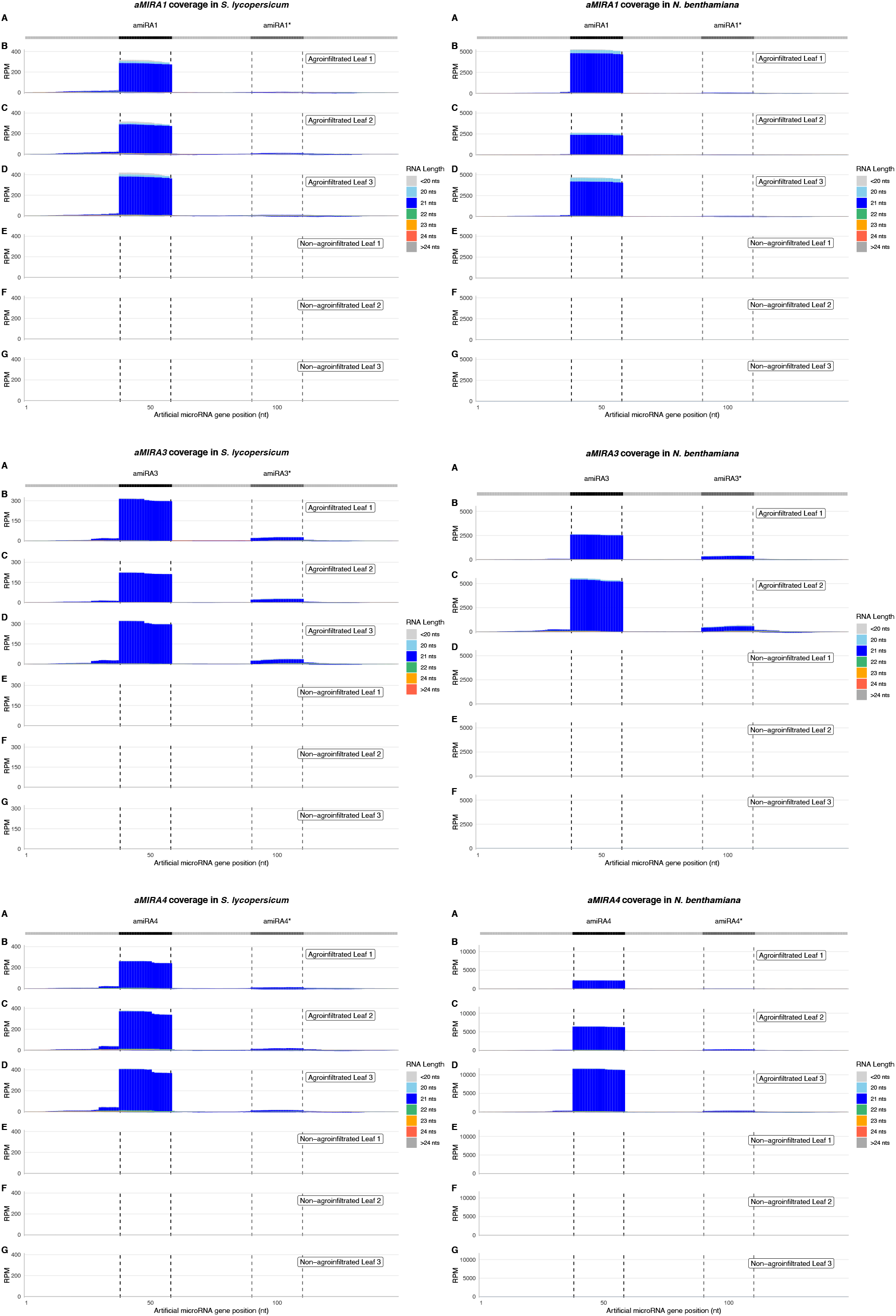
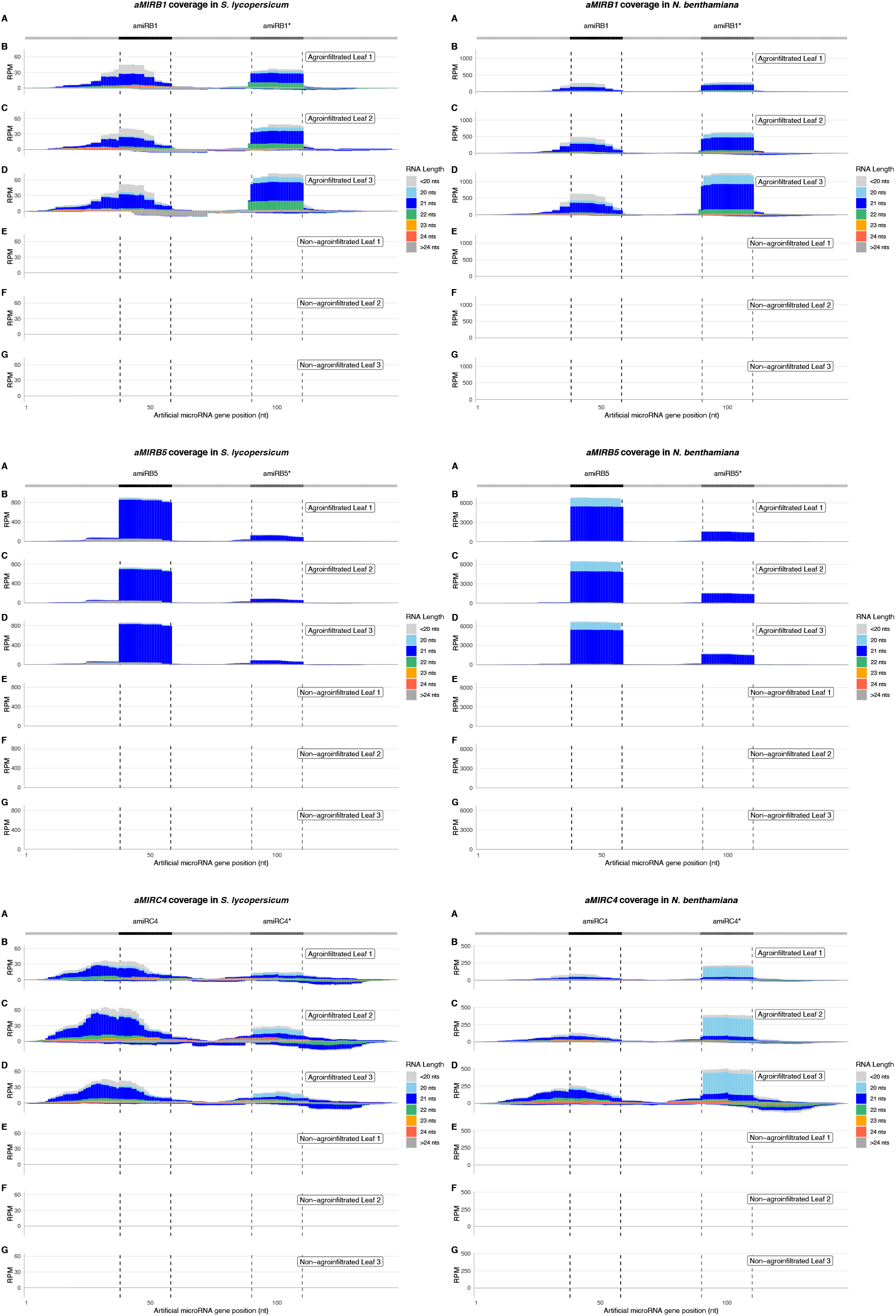
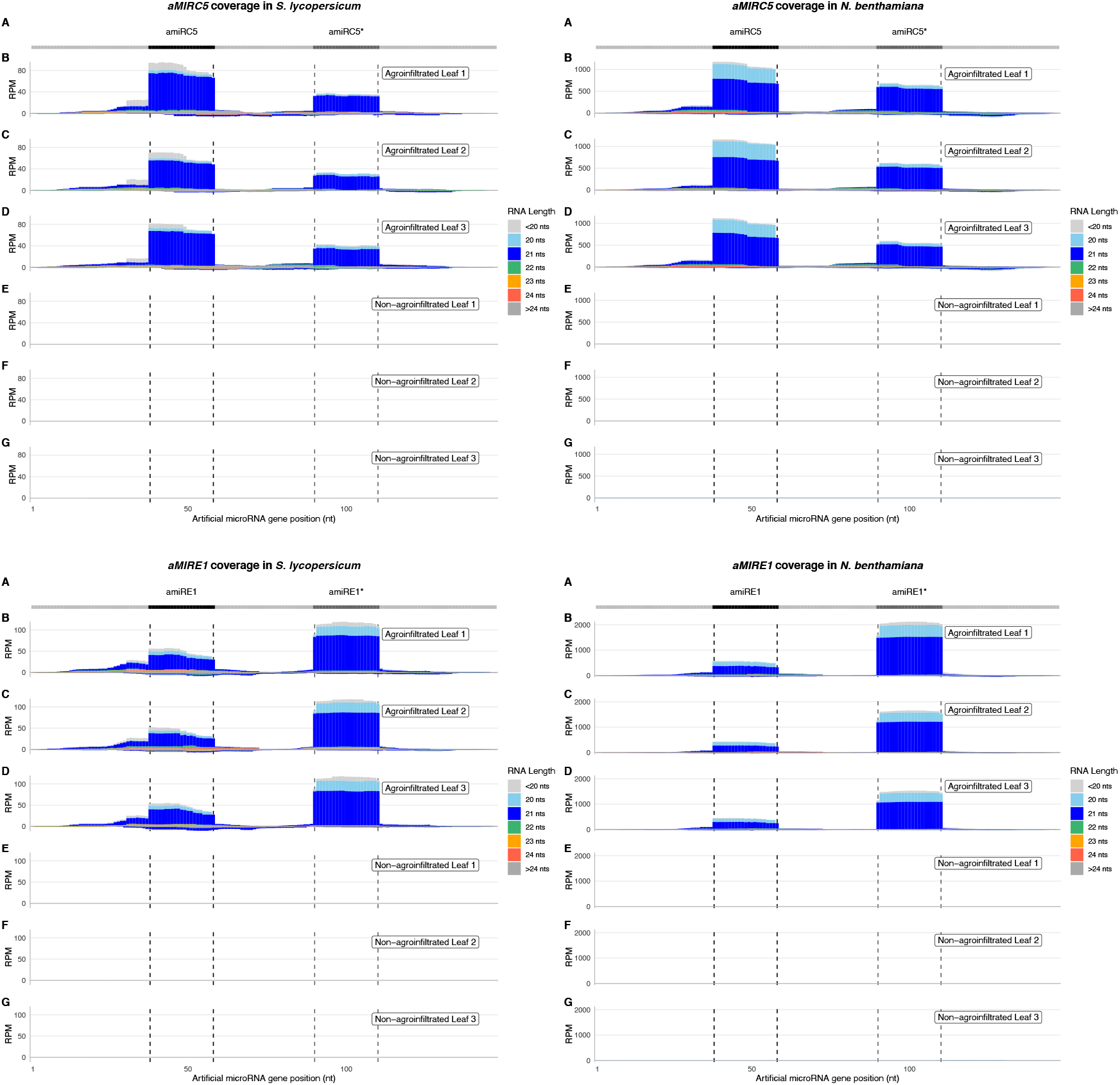
sRNA coverage of the artificial monocistronic microRNA genes in each individual sample. Each plot represents the sRNA coverage of a specific *aMIRNA* gene in tomato, on the left, and *N. benthamiana*, on the right. At the top (A), the light grey line corresponds to the precursors’ backbone; the position of the amiRNA and amiRNA* in the precursors are indicated in black and dark grey respectively. The three top plots (B, C, D) show the coverage in the three leaves agroinfiltrated with the corresponding *aMIRNA* construct; the three bottom plots (E, F, G) show the coverage in the three control, non-agroinfiltrated leaves. The x axis indicates the position on the gene in nucleotides, from 5’ to 3’. The y axis is the sRNA coverage in RPM for each nucleotide. Positive coverage means the reads align to the + DNA strand, negative coverage means the reads align to the - DNA strand. The coverage of reads with different lengths is represented separately with different colors, stacked from bottom to top according to the legend on the right. Coverages of all reads shorter than 20 nucleotides and longer than 24 nucleotides are represented in light grey and dark grey respectively.

**Supplemental Figure 3.**
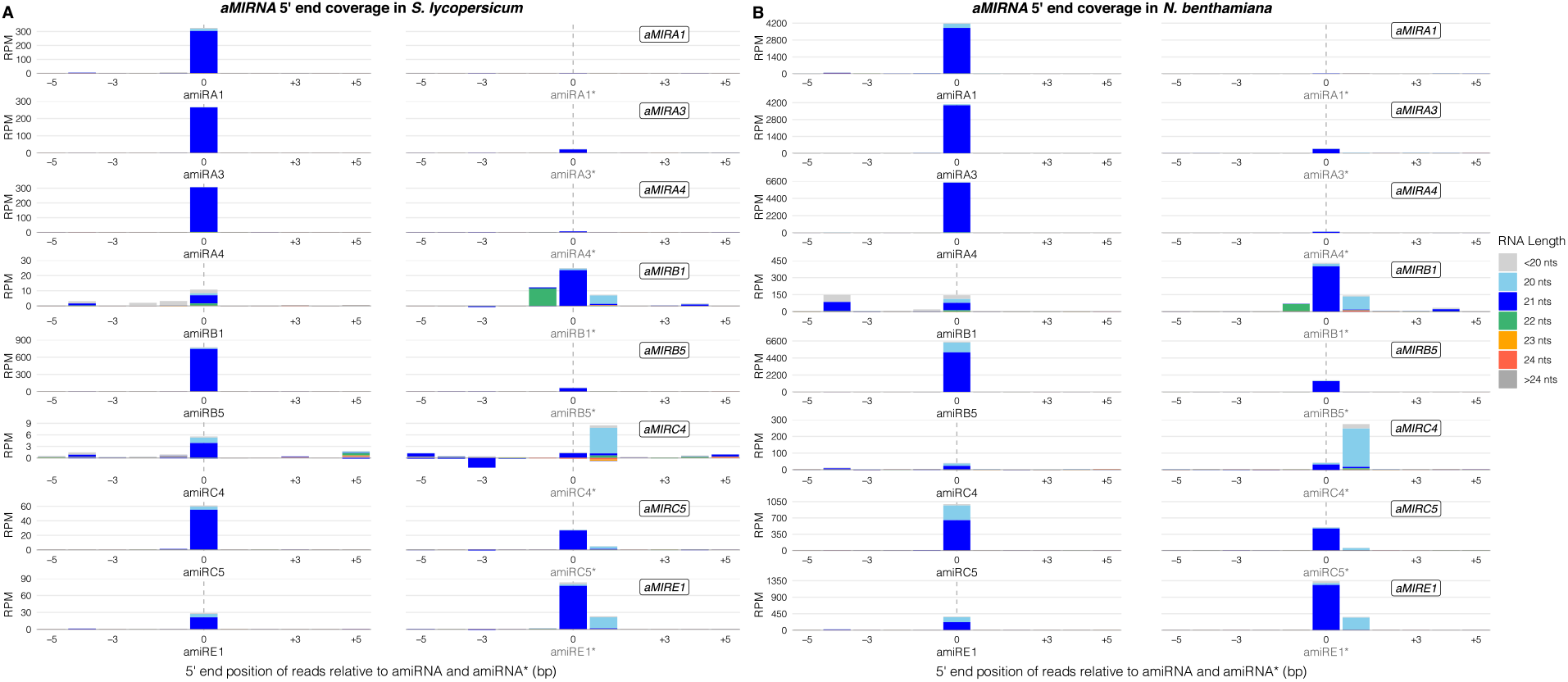
sRNA 5’ coverage around the artificial microRNA and microRNA* 5’ ends in the artificial monocistronic microRNA genes. A) Tomato sRNA 5’ coverage. In the x axis, 0 indicates the 5’ end of the amiRNAs and amiRNA*s, −5 and +5 indicate five nucleotides upstream and downstream of them. The y axis is the sRNA 5’ coverage in RPM. Positive coverage means the reads align to the + DNA strand, negative coverage means the reads align to the - DNA strand. The sRNA 5’ coverage of reads with different lengths is represented separately with different colors, stacked from bottom to top according to the legend on the right. Coverages of all reads shorter than 20 nucleotides and longer than 24 nucleotides are represented in light grey and dark grey respectively. B) *N. benthamiana* sRNA 5’ coverage.

**Supplemental Figure 4.**
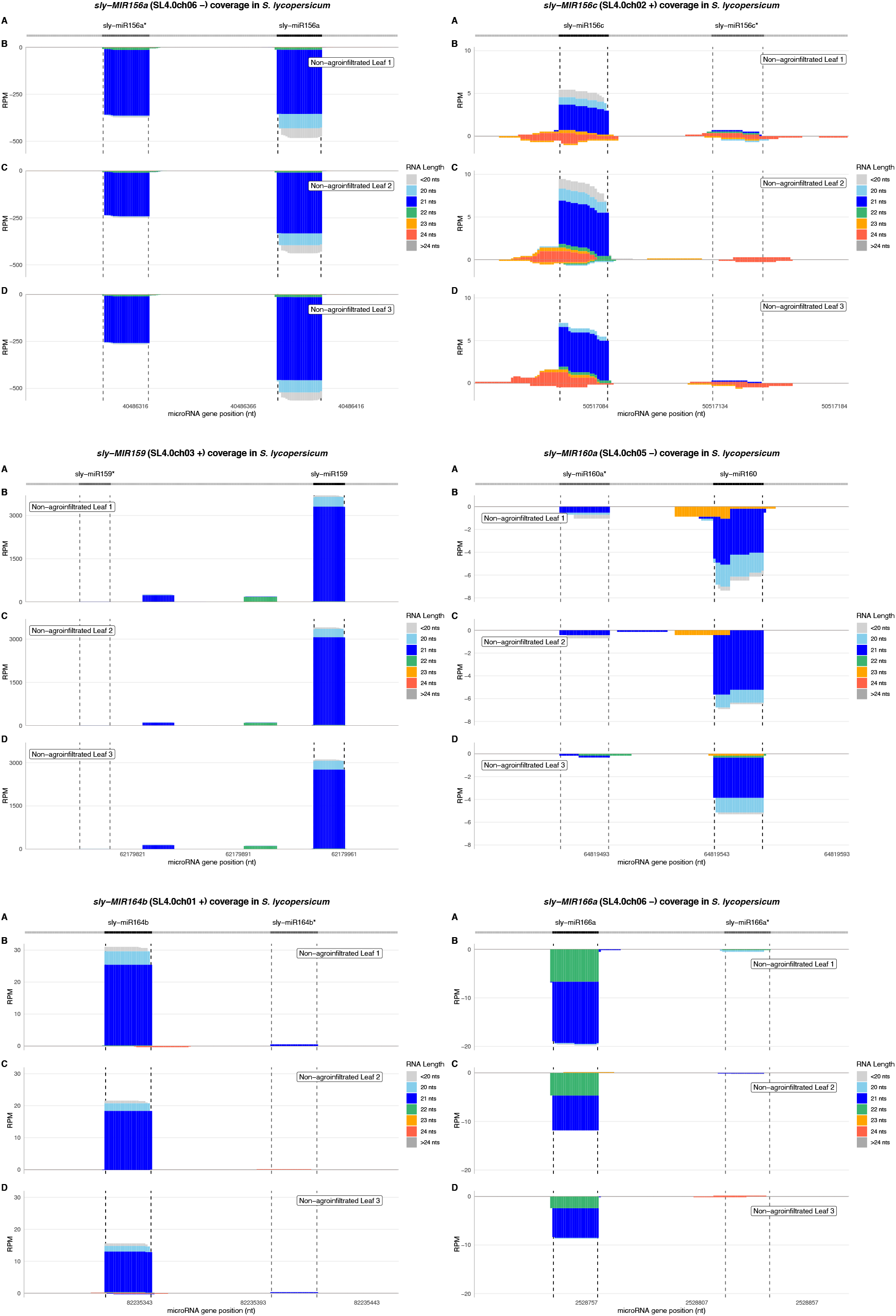
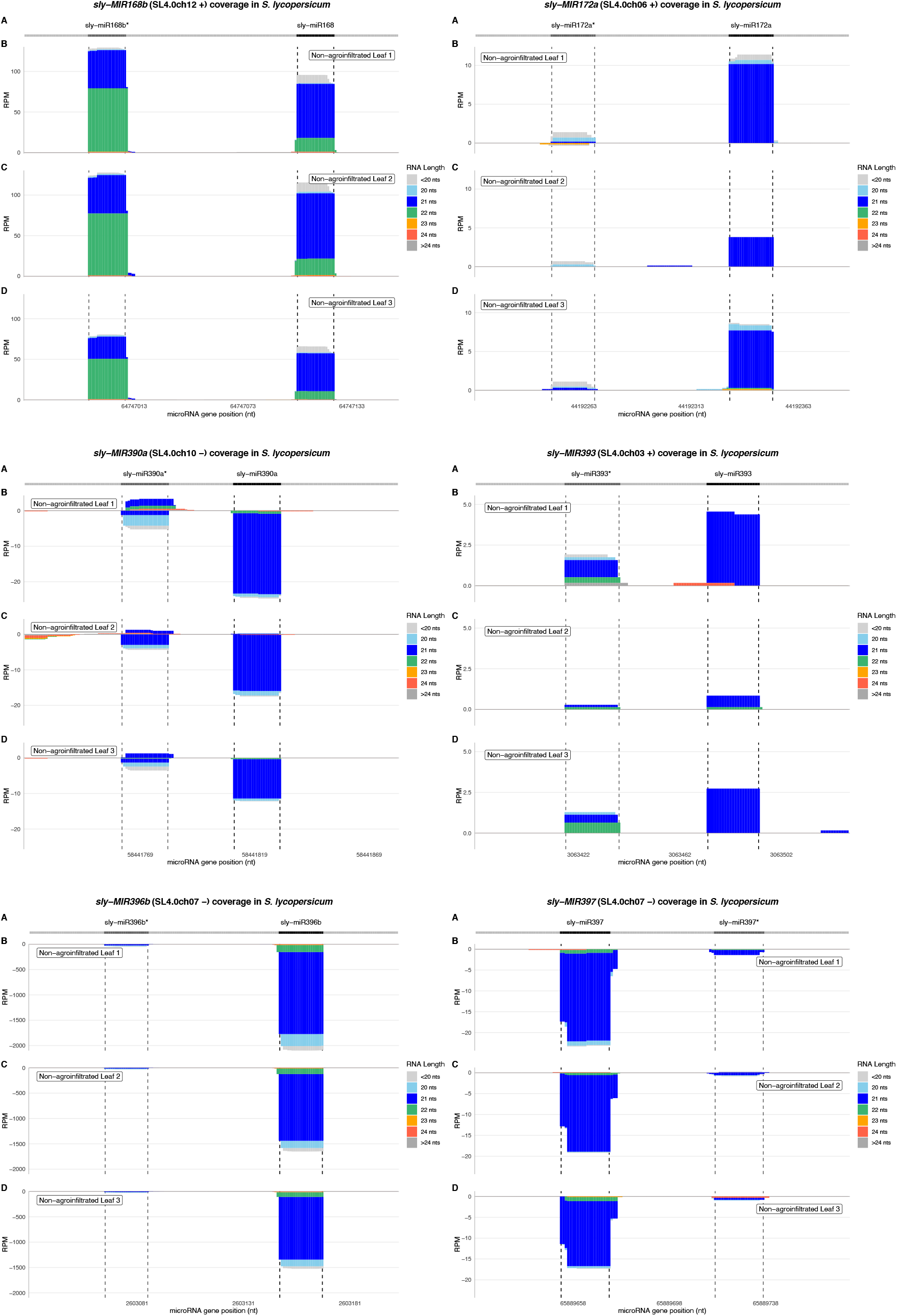
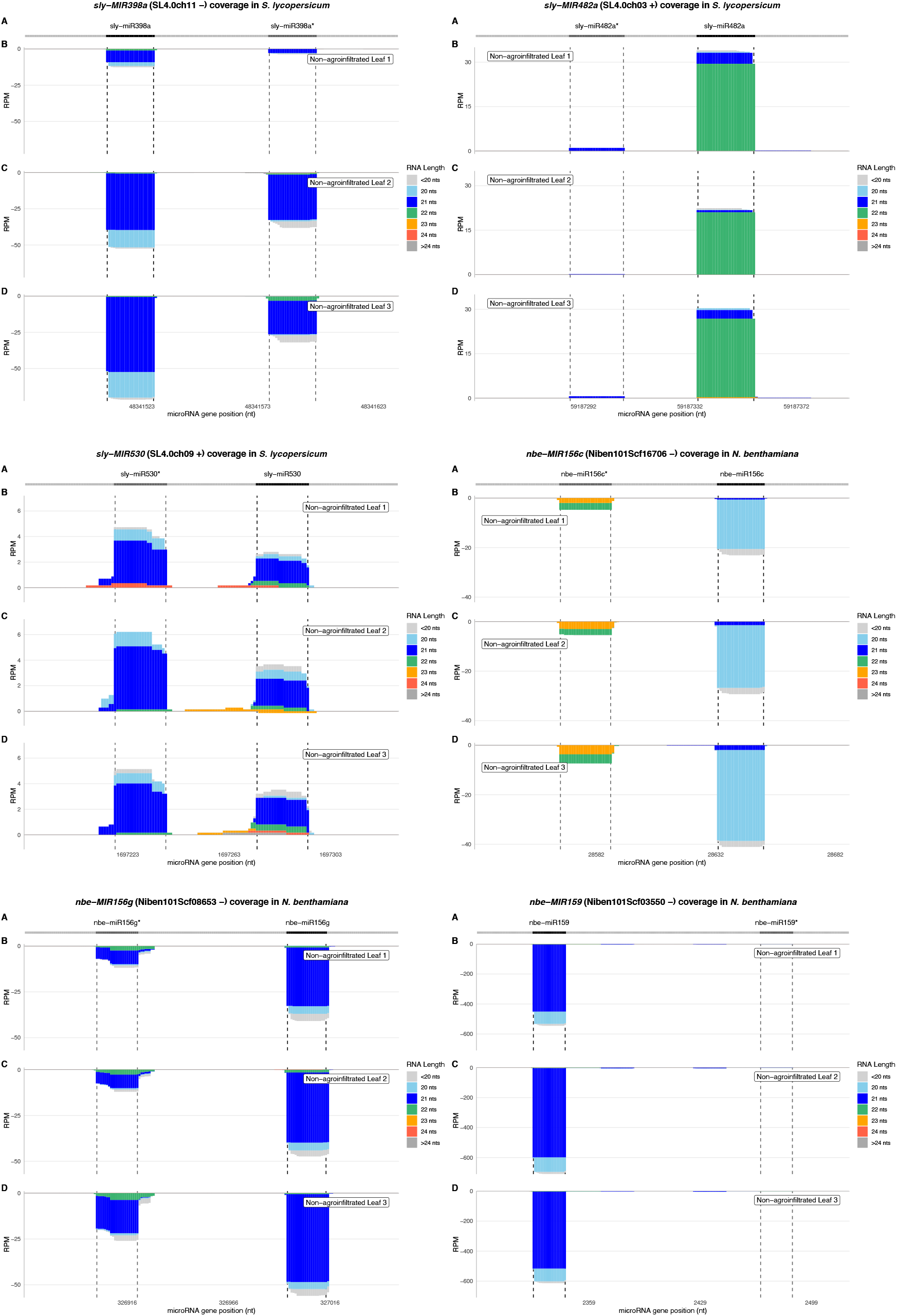
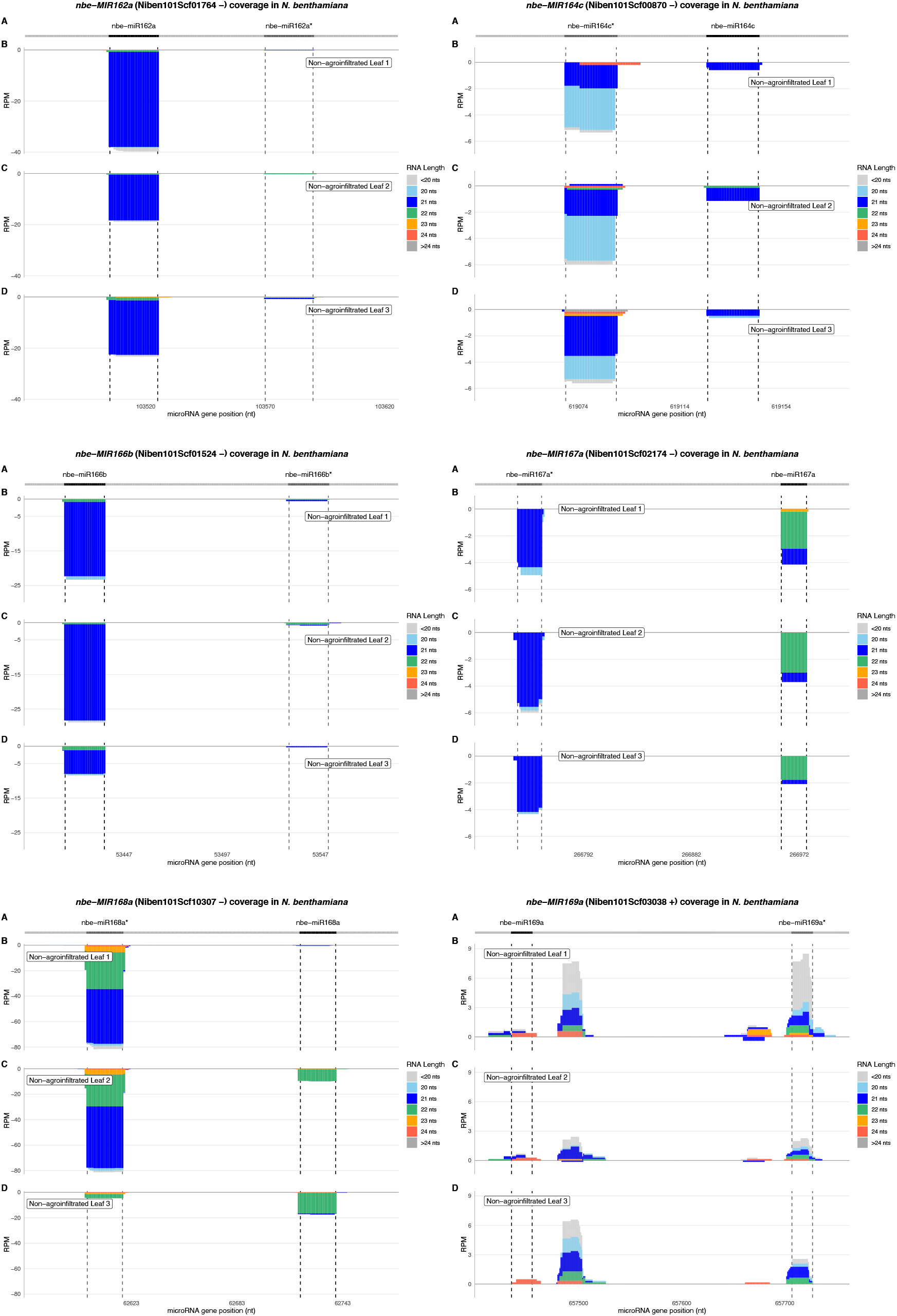
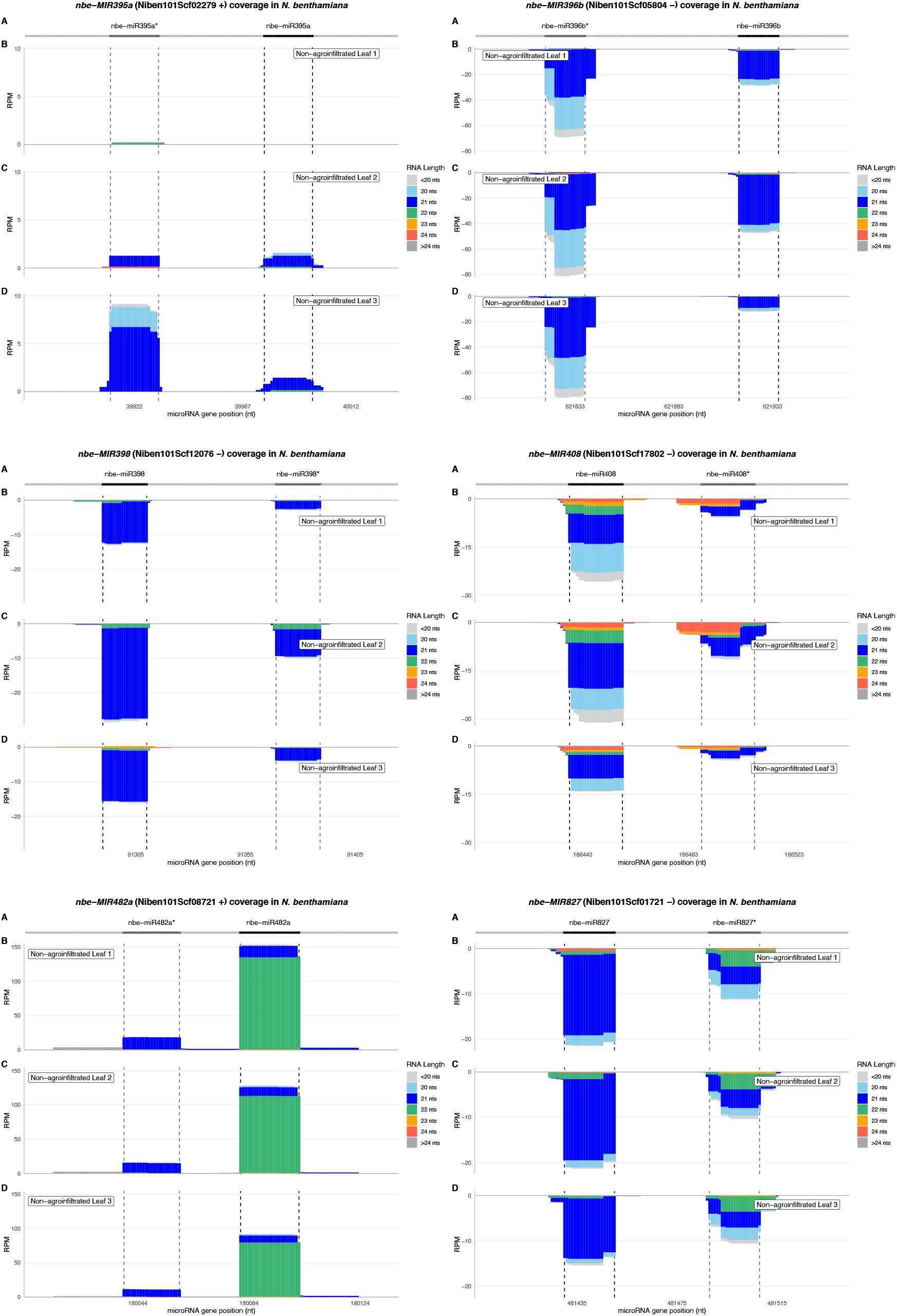
sRNA coverage of the endogenous microRNA genes in each control sample. Each plot represents the sRNA coverage of a specific *MIRNA* gene in tomato or *N. benthamiana*. In the title between parentheses, the chromosome and the DNA strand of the *MIRNA* genes are indicated. At the top (A), the light grey line corresponds to the precursors’ backbone; the position of the miRNA and miRNA* in the precursors are indicated in black and dark grey respectively. B, C and D show the coverage in the three control, non-agroinfiltrated leaves. The x axis indicates the position on the chromosomes in nucleotides, from 5’ to 3’. The y axis is the sRNA coverage in RPM for each nucleotide. Positive coverage means the reads align to the + DNA strand, negative coverage means the reads align to the - DNA strand. The coverage of reads with different lengths is represented separately with different colors, stacked from bottom to top according to the legend on the right. Coverages of all reads shorter than 20 nucleotides and longer than 24 nucleotides are represented in light grey and dark grey respectively.

**Supplemental Figure 5.**
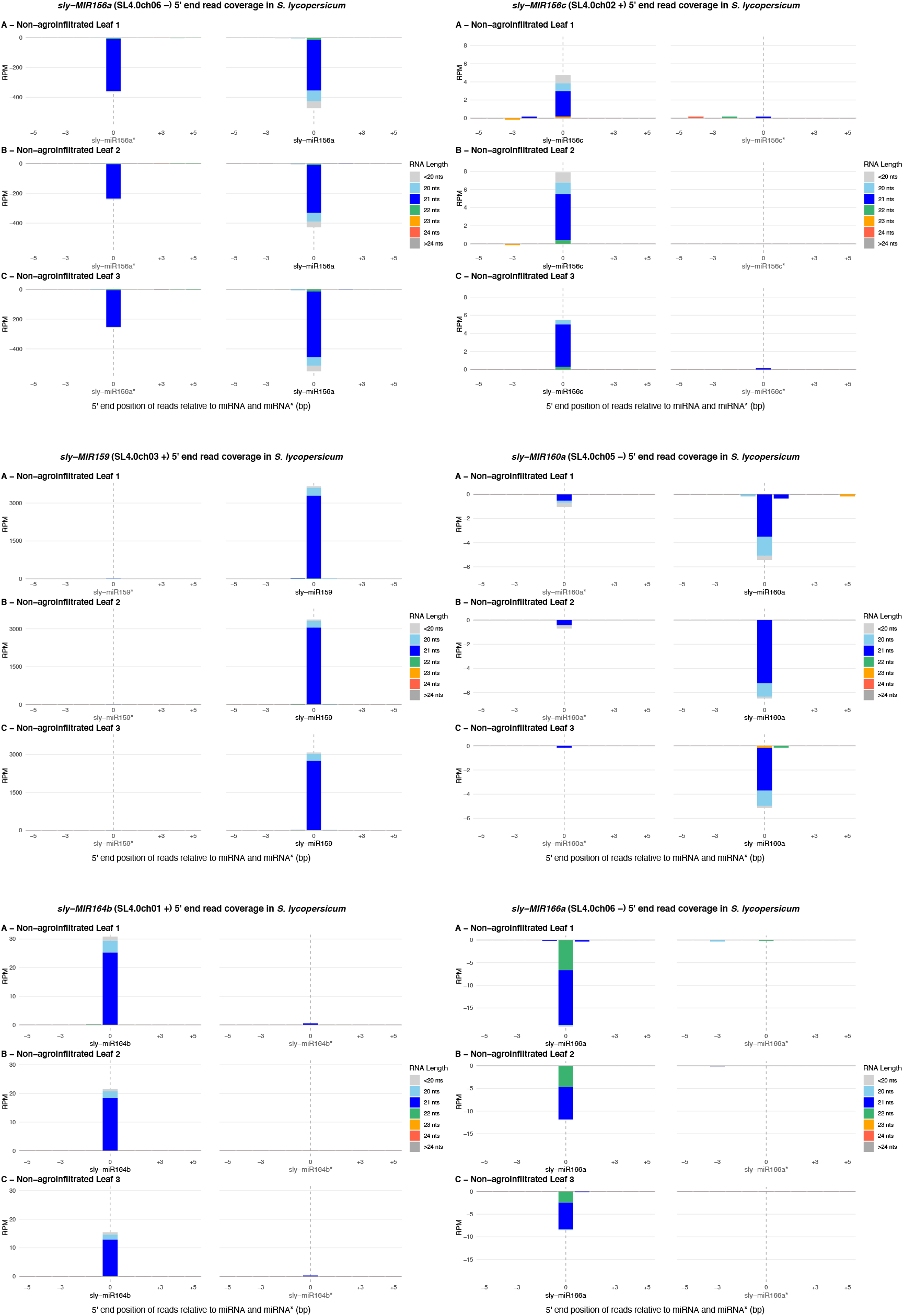
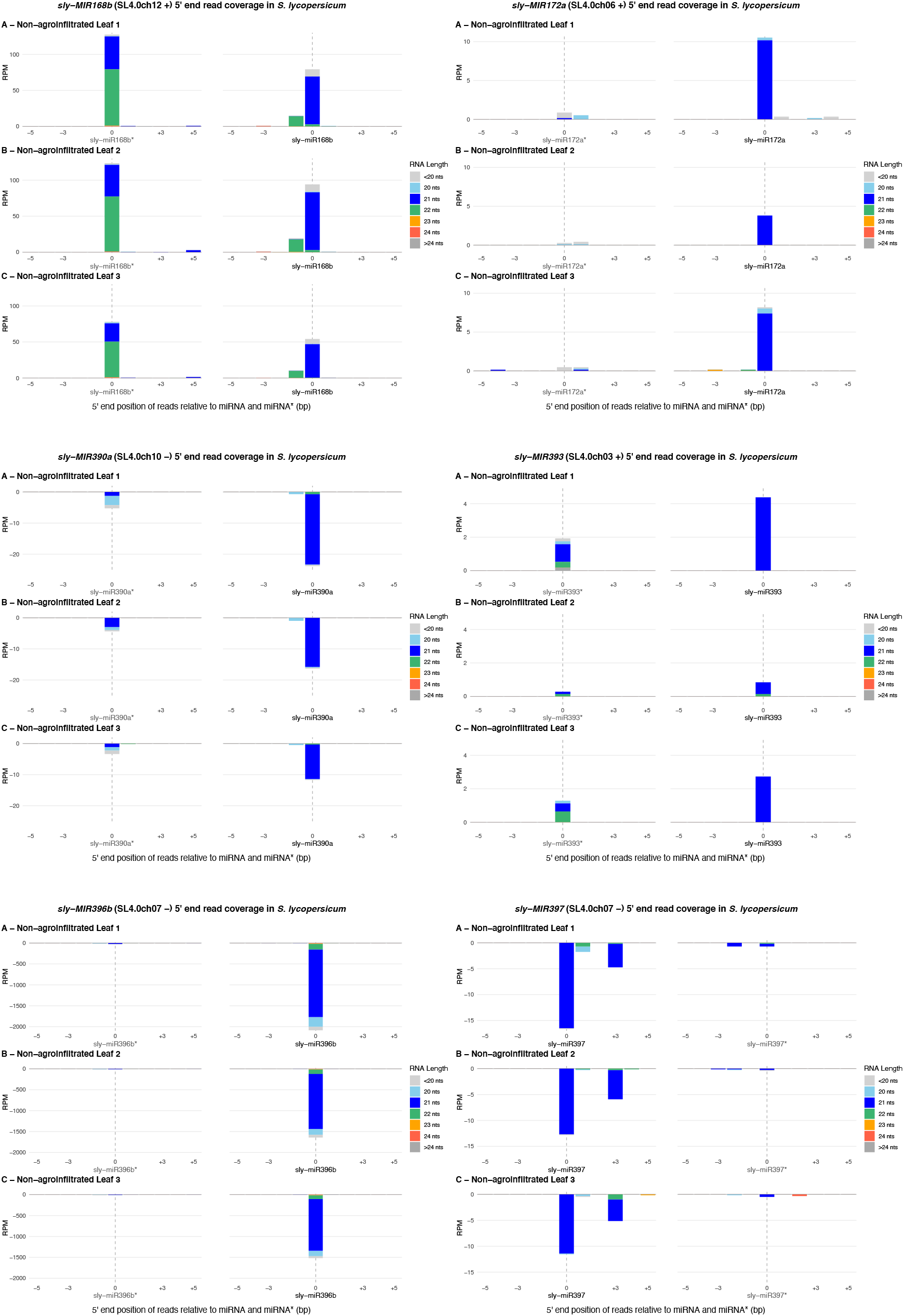
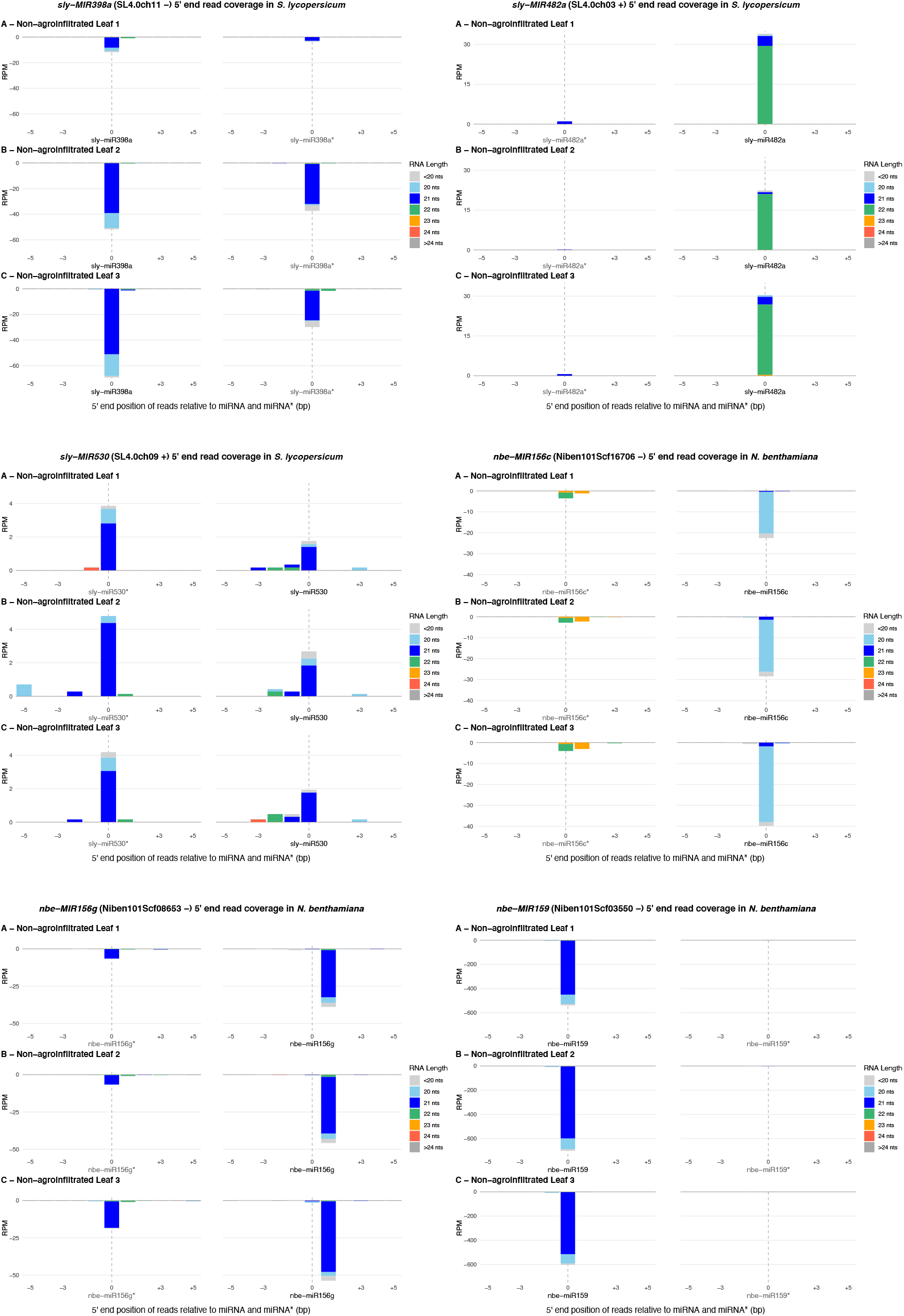
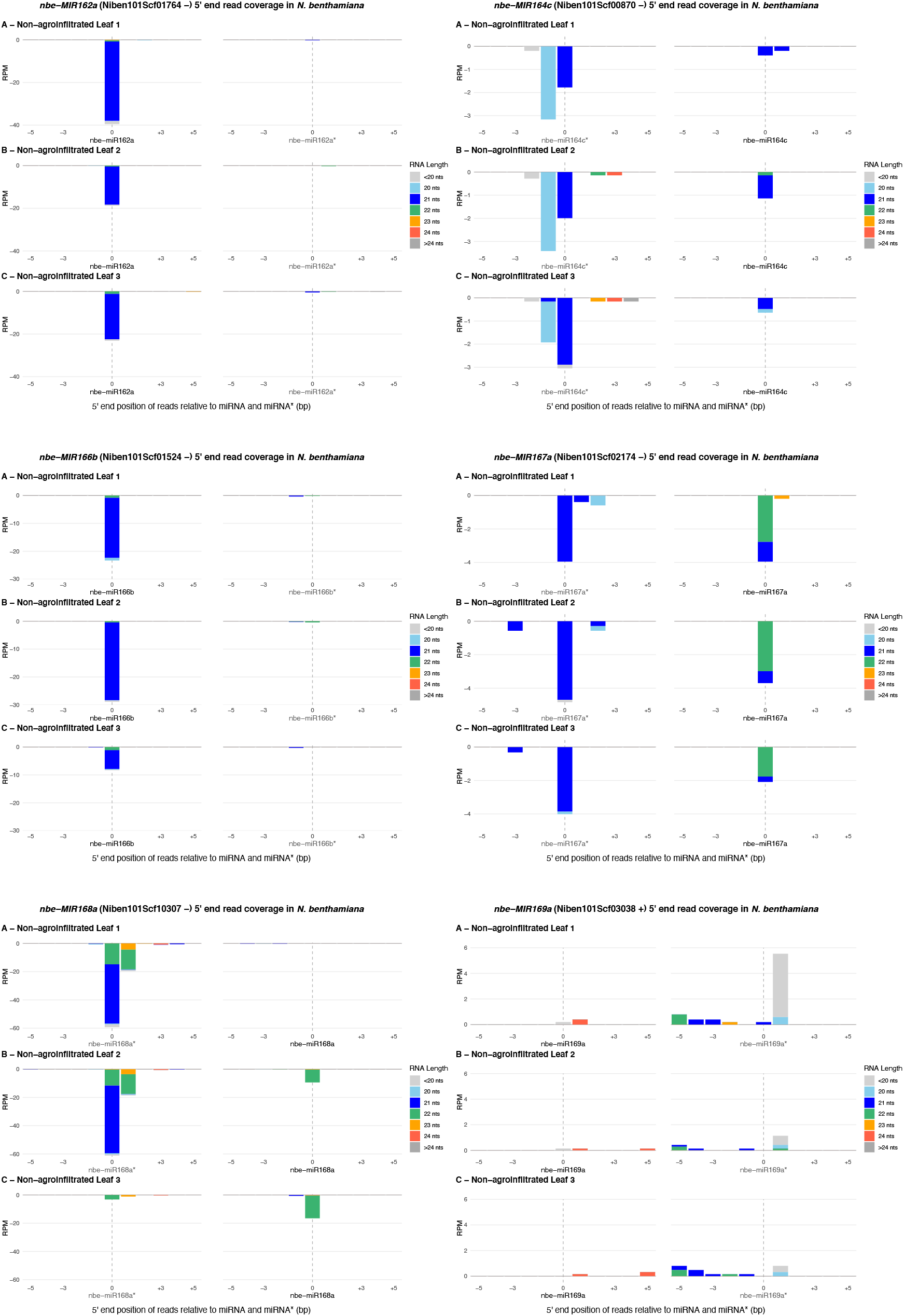
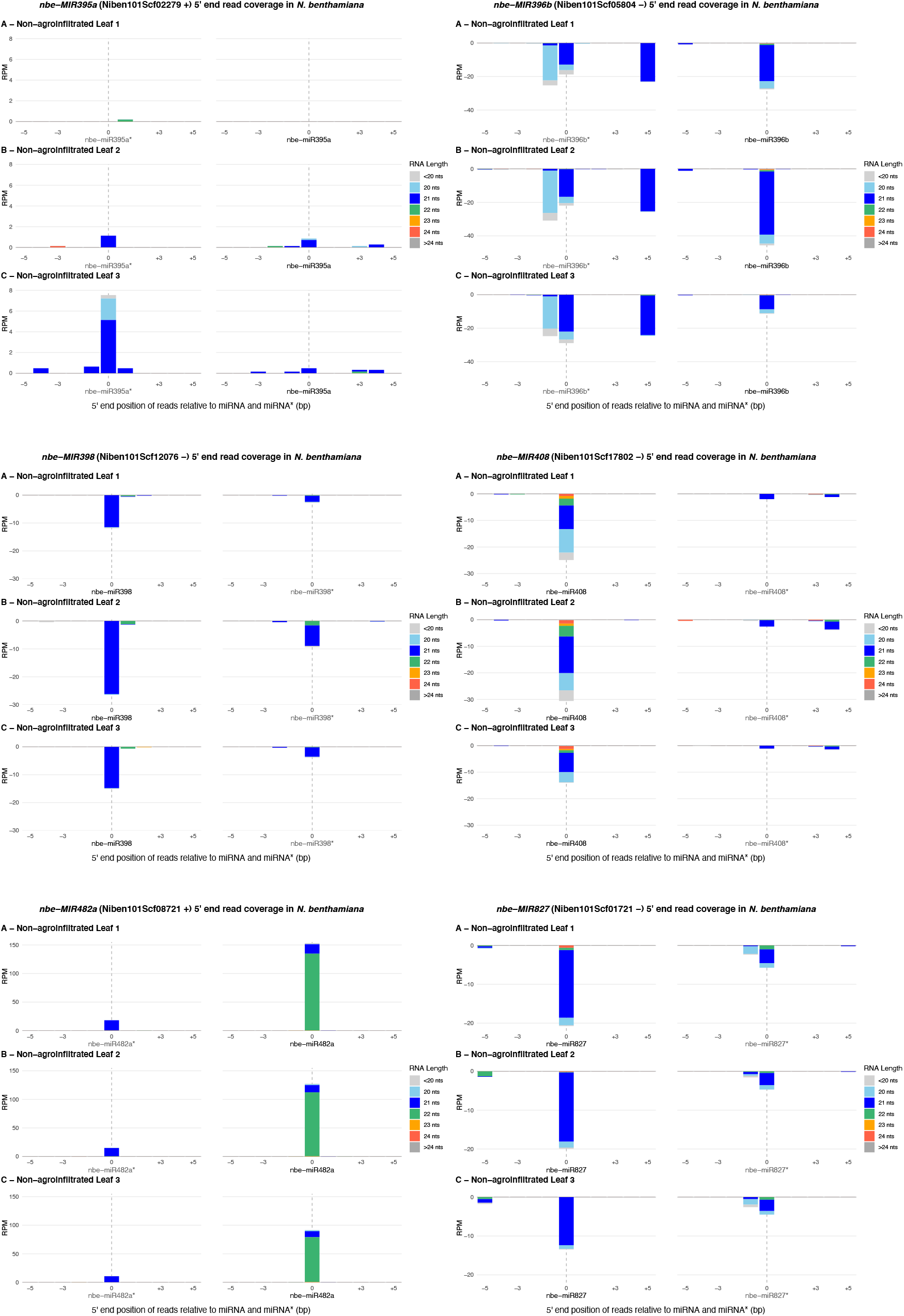
sRNA 5’ coverage around the microRNA and microRNA* 5’ ends in the endogenous microRNA genes. Each plot represents the sRNA 5’ coverage of a specific *MIRNA* gene in tomato or *N. benthamiana*. In the title between parentheses, the chromosome and the DNA strand of the *MIRNA* genes are indicated. A, B and C show the coverage in the three control, non-agroinfiltrated leaves. In the x axis, 0 indicates the 5’ end of the miRNAs and miRNA*s, −5 and +5 indicate five nucleotides upstream and downstream of them. The y axis is the sRNA 5’ coverage in RPM. Positive coverage means the reads align to the + DNA strand, negative coverage means the reads align to the - DNA strand. The sRNA 5’ coverage of reads with different lengths is represented separately with different colors, stacked from bottom to top according to the legend on the right. Coverages of all reads shorter than 20 nucleotides and longer than 24 nucleotides are represented in light grey and dark grey respectively.

**Supplemental Figure 6.**
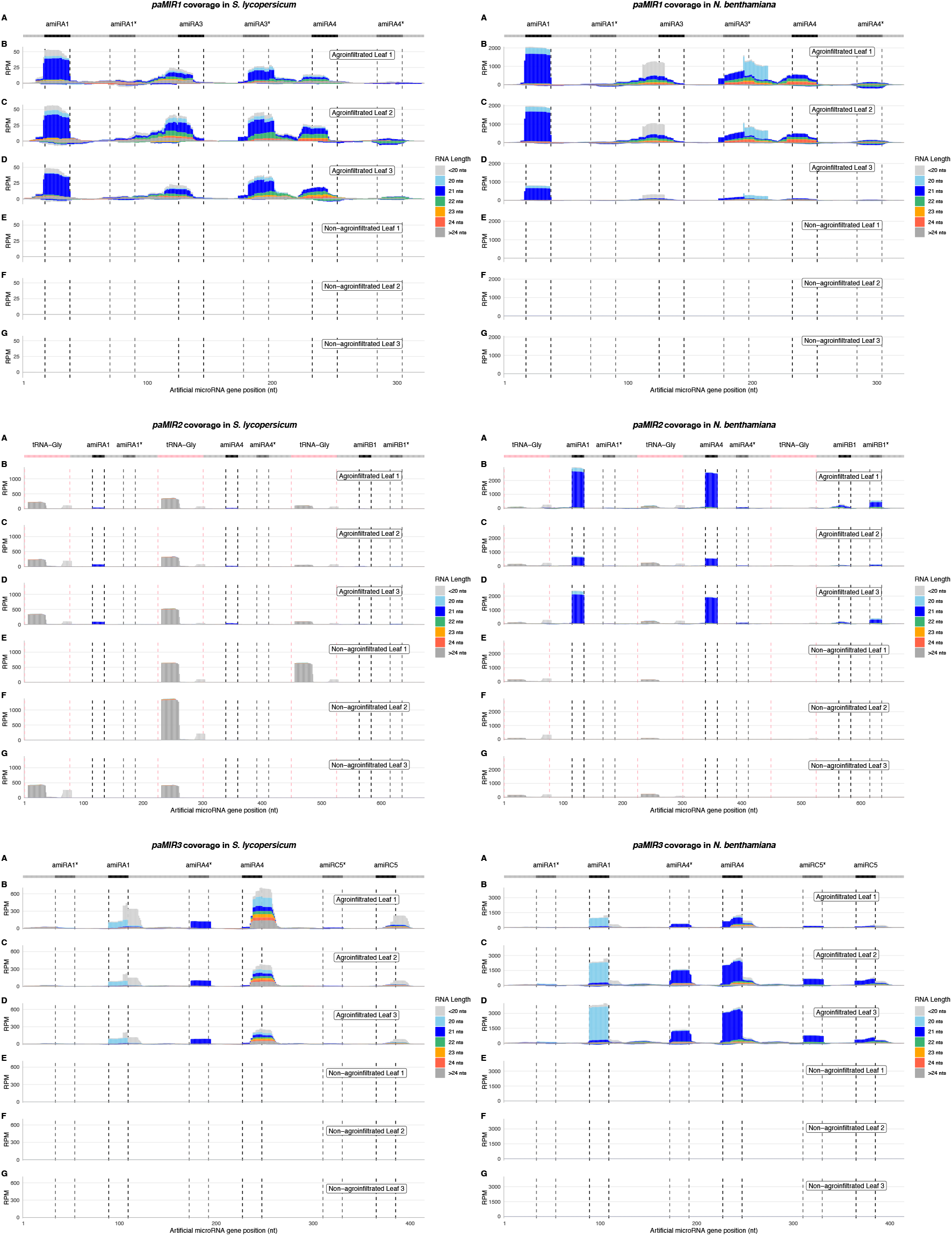
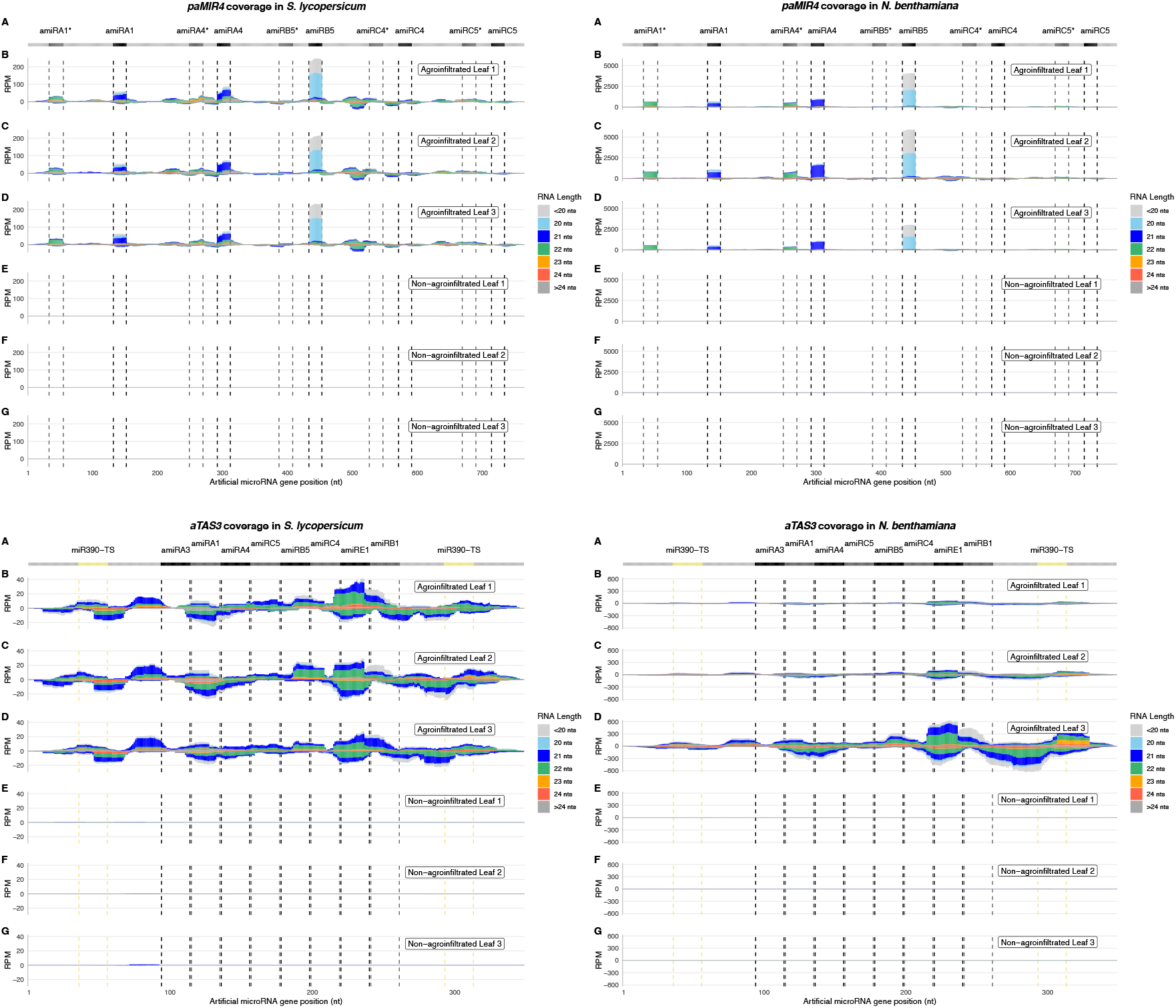
sRNA coverage of the artificial polycistronic microRNA and *trans*-acting siRNA genes in each individual sample. Each plot represents the sRNA coverage of a specific *aMIRNA* or *aTAS* gene in tomato, on the left, and *N. benthamiana*, on the right. At the top (A), the light grey line corresponds to the precursors’ backbone; the position of the amiRNA and amiRNA* in the precursors are indicated in black and dark grey respectively, except for *aTAS3* where these colors are used to distinguish adjacent amiRNAs; pink indicates the tRNA^Gly^ sequences; yellow indicates the miR390 target sites. The three top plots (B, C, D) show the coverage in the three leaves agroinfiltrated with the corresponding *aMIRNA* or *aTAS* construct; the three bottom plots (E, F, G) show the coverage in the three control, non-agroinfiltrated leaves. The x axis indicates the position on the gene in nucleotides, from 5’ to 3’. The y axis is the sRNA coverage in RPM for each nucleotide. Positive coverage means the reads align to the + DNA strand, negative coverage means the reads align to the - DNA strand. The coverage of reads with different lengths is represented separately with different colors, stacked from bottom to top according to the legend on the right. Coverages of all reads shorter than 20 nucleotides and longer than 24 nucleotides are represented in light grey and dark grey respectively.

**Supplemental Figure 7.**
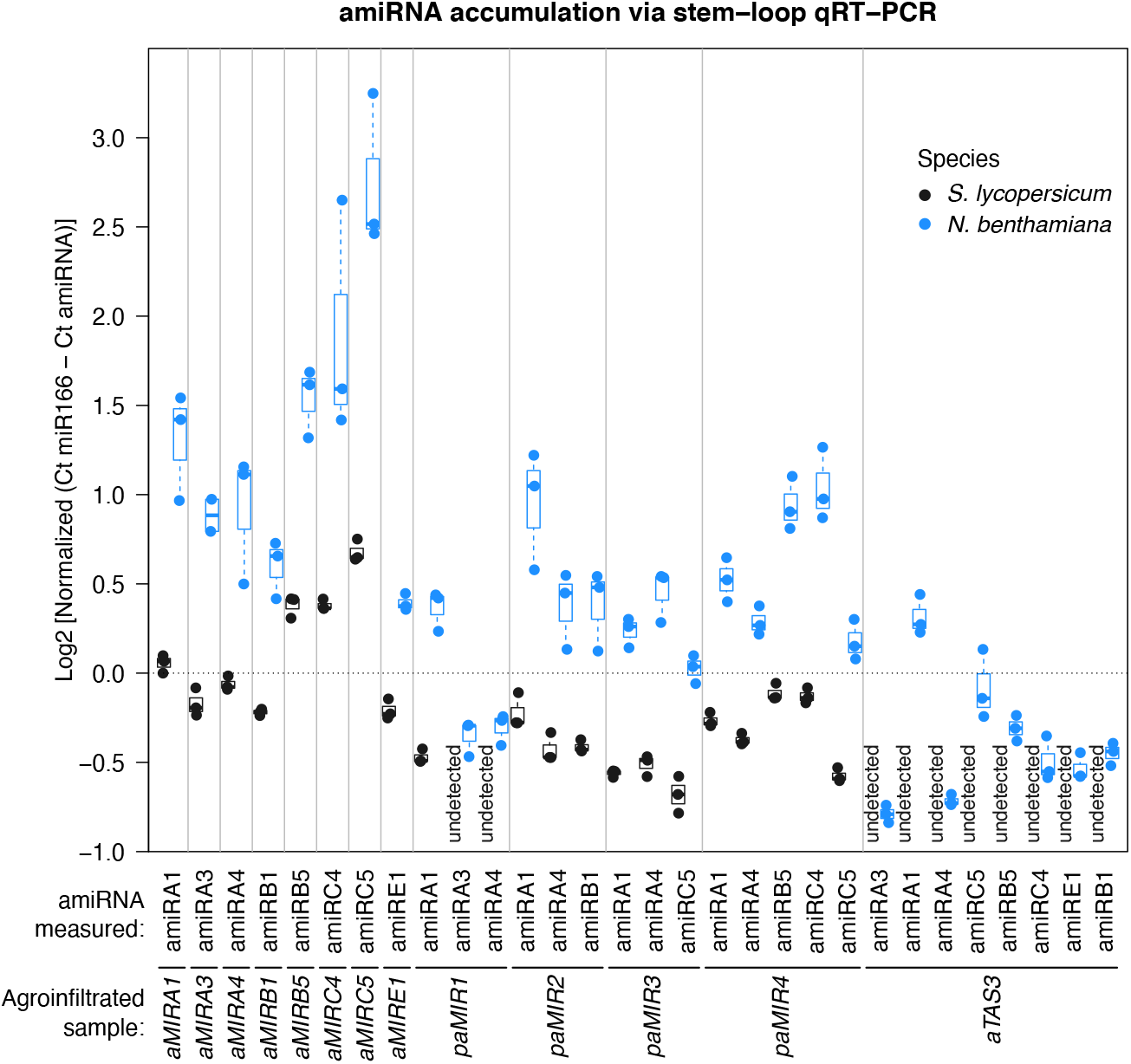
Stem-loop qRT-PCR detection of artificial microRNAs. See Supplemental Code 8 for generating this plot. The y axis shows the abundance of the mature amiRNAs measured through stem-loop qRT-PCR, as the binary logarithm of the difference between the cycle threshold (Ct) of the loading control miR166 and the amiRNA, normalized to the leftmost data point. The x axis shows which mature amiRNA was measured (top line) in which sample (bottom line). Different data points in the same column are replicates. Black data points are measured in tomato, blue data points are measured in *N. benthamiana*. With ‘undetected’, both amiRNAs without signal and false positives are indicated.

