## Supplemental Tables, Code, and Data for "Expression and processing of polycistronic artificial microRNAs and *trans*-acting siRNAs in *Solanum lycopersicum* and *Nicotiana benthamiana*": Supplemental_Code_1.R

```
# This code generates Figure 2

# Set the working directory by replacing "/your/path" with
# the complete path to the directory where this file is
setwd("/your/path")

# Load the data
tomato <- read.csv(file="20-24_tomato.csv", head=TRUE)
benthy <- read.csv(file="20-24_Nbenthamiana.csv", head=TRUE)

# Just for safety in case there is a lingering attachment
detach(tomato)
detach(benthy)

# Attach the namespace
attach(tomato)
attach(benthy)

tomato$miRNA = factor(tomato$miRNA, unique(tomato$miRNA))
tomato$miRNA

benthy$miRNA = factor(benthy$miRNA, unique(benthy$miRNA))
benthy$miRNA

# Define colors
# A1:firebrick1
# A3:dodgerblue
# A4:chartreuse3
# B1:purple1
# C5:orange
# B5:orchid1
# C4:saddlebrown
# E1:aquamarine2
# TAS3:gray
# endogenous:gray30

# Create a multi-paneled plot and define margins 
par(mfrow=c(2,1), mar=c(6, 4, 4, 10.5) + 0.1, xpd=TRUE)

# Plot tomato data
boxplot(tomato$precision~tomato$miRNA,
           ylim=c(0,1),
           vertical=TRUE,
           xaxt = "n",
           border=c("firebrick1","dodgerblue","chartreuse3","purple1","orchid1","saddlebrown","orange","aquamarine2",
                    "firebrick1","dodgerblue","chartreuse3",
                    "firebrick1","chartreuse3","purple1",
                    "firebrick1","chartreuse3","orange",
                    "firebrick1","chartreuse3","orchid1","saddlebrown","orange",
                    "gray",
                    "gray30","gray30","gray30","gray30","gray30","gray30","gray30","gray30","gray30","gray30","gray30","gray30","gray30","gray30","gray30"),
           cex=1,cex.axis=1.2, cex.lab=1.2,
           pch=20,outcex=0.75,
           ylab="Precision of processing",
           xlab = NA,
           las=2,
           main=expression(bolditalic("Solanum lycopersicum")))

stripchart(tomato$precision~tomato$miRNA,
        ylim=c(0,1),
        vertical=TRUE,
        xaxt = "n",
        pch=20,col=c("firebrick1","dodgerblue","chartreuse3","purple1","orchid1","saddlebrown","orange","aquamarine2",
                     "firebrick1","dodgerblue","chartreuse3",
                     "firebrick1","chartreuse3","purple1",
                     "firebrick1","chartreuse3","orange",
                     "firebrick1","chartreuse3","orchid1","saddlebrown","orange",
                     "gray",
                     "gray30","gray30","gray30","gray30","gray30","gray30","gray30","gray30","gray30","gray30","gray30","gray30","gray30","gray30","gray30"),
        cex=0.75,
        add=TRUE)

abline(v=c(1.5,2.5,3.5,4.5,5.5,6.5,7.5,8.5,11.5,14.5,17.5,22.5,23.5), lty=2, col="grey", xpd=FALSE)

text(x=c(1,2,3,4,5,6,7,8,10,13,16,20,23,24,25,26,27,28,29,30,31,32,33,34,35,36,37,38), y=-0.05, adj=1, srt = 90, cex =1.1,
     labels=c(expression(italic("aMIRA1")),expression(italic("aMIRA3")),expression(italic("aMIRA4")),expression(italic("aMIRB1")),expression(italic("aMIRB5")),expression(italic("aMIRC4")),expression(italic("aMIRC5")),expression(italic("aMIRE1")),
              expression(italic("paMIR1")),expression(italic("paMIR2")),expression(italic("paMIR3")),expression(italic("paMIR4")),expression(italic("aTAS3")),
              expression(italic("sly-MIR156a")),expression(italic("sly-MIR156c")),expression(italic("sly-MIR159")),expression(italic("sly-MIR160a")),expression(italic("sly-MIR164b")),
              expression(italic("sly-MIR166a")),expression(italic("sly-MIR168b")),expression(italic("sly-MIR172a")),expression(italic("sly-MIR390a")),expression(italic("sly-MIR393")),
              expression(italic("sly-MIR396b")),expression(italic("sly-MIR397")),expression(italic("sly-MIR398a")),expression(italic("sly-MIR482a")),expression(italic("sly-MIR530"))), xpd=TRUE,
     col="black")

text(x=-4, y=1.22, adj=1, srt = 0, cex =1.5,
     labels="A", xpd=TRUE, col="black", font=2)

legend("topright", c("amiRA1","amiRA3","amiRA4","amiRB1","amiRB5","amiRC4","amiRC5","amiRE1",expression("amiRs in"~italic("aTAS3")),"endogenous miRNAs","in control samples"),
       pch=15, col=c("firebrick1","dodgerblue","chartreuse3","purple1","orchid1","saddlebrown","orange","aquamarine2","gray","gray30","white"),
       xpd=TRUE, pt.cex = 1.8, cex = 1, inset=c(-0.25,0),
       title="miRNA measured", title.adj=0.1, box.lty=0) 

# Plot N. benthamiana data
##nbe-MIR395a: NB_Non_agro_1 is removed because this miRNA is not expressed,
##nbe-MIR168a: NB_Non_agro_3 is removed because this miRNA is not expressed
boxplot(benthy$precision~benthy$miRNA,
        ylim=c(0,1),
        vertical=TRUE,
        xaxt = "n",
        border=c("firebrick1","dodgerblue","chartreuse3","purple1","orchid1","saddlebrown","orange","aquamarine2",
                 "firebrick1","dodgerblue","chartreuse3",
                 "firebrick1","chartreuse3","purple1",
                 "firebrick1","chartreuse3","orange",
                 "firebrick1","chartreuse3","orchid1","saddlebrown","orange",
                 "gray",
                 "gray30","gray30","gray30","gray30","gray30","gray30","gray30","gray30","gray30","gray30","gray30","gray30","gray30","gray30","gray30"),
        cex=1,cex.axis=1.2, cex.lab=1.2,
        pch=20,outcex=0.75,
        ylab="Precision of processing",
        xlab = NA,
        las=2,
        main=expression(bolditalic("Nicotiana benthamiana")))

stripchart(benthy$precision~benthy$miRNA,
           ylim=c(0,1),
           vertical=TRUE,
           xaxt = "n",
           pch=20,col=c("firebrick1","dodgerblue","chartreuse3","purple1","orchid1","saddlebrown","orange","aquamarine2",
                        "firebrick1","dodgerblue","chartreuse3",
                        "firebrick1","chartreuse3","purple1",
                        "firebrick1","chartreuse3","orange",
                        "firebrick1","chartreuse3","orchid1","saddlebrown","orange",
                        "gray",
                        "gray30","gray30","gray30","gray30","gray30","gray30","gray30","gray30","gray30","gray30","gray30","gray30","gray30","gray30","gray30"),
           cex=0.75,
           add=TRUE)

abline(v=c(1.5,2.5,3.5,4.5,5.5,6.5,7.5,8.5,11.5,14.5,17.5,22.5,23.5), lty=2, col="grey", xpd=FALSE)

text(x=c(1,2,3,4,5,6,7,8,10,13,16,20,23,24,25,26,27,28,29,30,31,32,33,34,35,36,37,38), y=-0.05, adj=1, srt = 90, cex =1.1,
     labels=c(expression(italic("aMIRA1")),expression(italic("aMIRA3")),expression(italic("aMIRA4")),expression(italic("aMIRB1")),expression(italic("aMIRB5")),expression(italic("aMIRC4")),expression(italic("aMIRC5")),expression(italic("aMIRE1")),
              expression(italic("paMIR1")),expression(italic("paMIR2")),expression(italic("paMIR3")),expression(italic("paMIR4")),expression(italic("aTAS3")),
              expression(italic("nbe-MIR156c")),expression(italic("nbe-MIR156g")),expression(italic("nbe-MIR159")),expression(italic("nbe-MIR162a")),expression(italic("nbe-MIR164c")),
              expression(italic("nbe-MIR166b")),expression(italic("nbe-MIR167a")),expression(italic("nbe-MIR168a")),expression(italic("nbe-MIR169a")),expression(italic("nbe-MIR395a")),
              expression(italic("nbe-MIR396b")),expression(italic("nbe-MIR398")),expression(italic("nbe-MIR408")),expression(italic("nbe-MIR482a")),expression(italic("nbe-MIR827"))), xpd=TRUE,
     col="black")

text(x=-4, y=1.22, adj=1, srt = 0, cex =1.5,
     labels="B", xpd=TRUE, col="black", font=2)

legend("topright", c("amiRA1","amiRA3","amiRA4","amiRB1","amiRB5","amiRC4","amiRC5","amiRE1",expression("amiRs in"~italic("aTAS3")),"endogenous miRNAs","in control samples"),
       pch=15, col=c("firebrick1","dodgerblue","chartreuse3","purple1","orchid1","saddlebrown","orange","aquamarine2","gray","gray30","white"),
       xpd=TRUE, pt.cex = 1.8, cex = 1, inset=c(-0.25,0),
       title="miRNA measured", title.adj=0.1, box.lty=0) 
```
