## Supplemental Tables, Code, and Data for "Expression and processing of polycistronic artificial microRNAs and *trans*-acting siRNAs in *Solanum lycopersicum* and *Nicotiana benthamiana*": Supplemental_Code_2.R

```
# This code generates Figure 3A

# Activate required non-base packages
library(ggplot2)
library(dplyr)
library(cowplot)
library(ggpubr)

# Set the working directory by replacing "/your/path" with
# the complete path to the directory where this file is
setwd("/your/path")

# Read files made by "Supplemental Code 8.pl"
coverage1 <- read.table(file="amiRA1_tomato.txt",
                       sep="\t",head=TRUE)
coverage2 <- read.table(file="amiRA3_tomato.txt",
                       sep="\t",head=TRUE)
coverage3 <- read.table(file="amiRA4_tomato.txt",
                        sep="\t",head=TRUE)
coverage4 <- read.table(file="amiRB1_tomato.txt",
                        sep="\t",head=TRUE)
coverage5 <- read.table(file="amiRB5_tomato.txt",
                        sep="\t",head=TRUE)
coverage6 <- read.table(file="amiRC4_tomato.txt",
                        sep="\t",head=TRUE)
coverage7 <- read.table(file="amiRC5_tomato.txt",
                        sep="\t",head=TRUE)
coverage8 <- read.table(file="amiRE1_tomato.txt",
                        sep="\t",head=TRUE)

# Read file determining the gene features
gene <- read.table(file="gene_amiRNAs.tsv",
                        sep="\t",head=TRUE)

# Filter the plus and minus sets
cov_plus1 <- filter(coverage1, Strand=="plus")
cov_minus1 <- filter(coverage1, Strand=="minus")
cov_plus2 <- filter(coverage2, Strand=="plus")
cov_minus2 <- filter(coverage2, Strand=="minus")
cov_plus3 <- filter(coverage3, Strand=="plus")
cov_minus3 <- filter(coverage3, Strand=="minus")
cov_plus4 <- filter(coverage4, Strand=="plus")
cov_minus4 <- filter(coverage4, Strand=="minus")
cov_plus5 <- filter(coverage5, Strand=="plus")
cov_minus5 <- filter(coverage5, Strand=="minus")
cov_plus6 <- filter(coverage6, Strand=="plus")
cov_minus6 <- filter(coverage6, Strand=="minus")
cov_plus7 <- filter(coverage7, Strand=="plus")
cov_minus7 <- filter(coverage7, Strand=="minus")
cov_plus8 <- filter(coverage8, Strand=="plus")
cov_minus8 <- filter(coverage8, Strand=="minus")

# Define color pallette
sRNAcols = c("lightgray", # <20 nts
             "skyblue",   # 20 nts
             "blue",      # 21 nts
             "mediumseagreen", # 22nts
             "orange",    # 23 nts
             "tomato",    # 24 nts
             "darkgray")  # >24 nts

miRNAcols = c("grey", # gene
              "black", # amiRNA
              "grey40") # amiRNA*

# Define the order to list the Length categories
Lorder = c("<20 nts","20 nts","21 nts","22 nts","23 nts","24 nts",">24 nts")
Lordem = c("gene","amiRNA","amiRNA*")

# Make plots
title <- ggdraw() + 
  draw_label(expression(bolditalic("aMIRNA")~bold("coverage in")~bolditalic("S. lycopersicum")),
             fontface = 'bold', hjust = 0.5, size = 17)

zero <- ggplot() +
  geom_col(data=gene,
           aes(x=Position, y=RPM,
               fill=factor(Type,levels=Lordem)), width=1) +
  scale_fill_manual(values=miRNAcols) +
  theme_minimal() +
  theme(panel.grid.major.x=element_blank(), panel.grid.minor.x=element_blank(), panel.grid.major.y=element_blank(),panel.grid.minor.y=element_blank()) +
  coord_cartesian(ylim = c(0, 10), xlim = c(0,148)) +
  scale_x_continuous(expand = c(0, 0), breaks=c(1,40,80,120)) + scale_y_continuous(expand = c(0, 0)) +
  theme(axis.text.y = element_text(color="white"), axis.text.x=element_blank()) +
  theme(axis.title.y = element_text(colour="white")) + 
  theme(axis.title.x = element_text(colour="white")) +
  theme(legend.position="none") +
  theme(plot.margin = unit(c(0, 0, 0, 0), "cm")) +
  labs(fill="RNA Length") +
  annotate("text", x=c(48,100), y=c(5,5), label=c("amiRNA","amiRNA*"), size=5)

one <- ggplot() +
  geom_vline(xintercept = c(0,38,58,90,110),
             linetype=c("solid","dashed","dashed","dashed","dashed"),
             color=c("grey70","black","black","grey40","grey40"),
             size=c(0.8,0.6,0.6,0.6,0.6)) +
  geom_hline(yintercept = 0, color = "grey70", size = 0.4) +
  geom_col(data=cov_plus1,
           aes(x=Position, y=RPM,
               fill=factor(Length,levels=Lorder)), width=1) +
  geom_col(data=cov_minus1,
           aes(x=Position, y=RPM,
               fill=factor(Length, levels=Lorder)), width=1) +
  labs(x="Artificial microRNA gene position (nt)") +
  coord_cartesian(ylim = c(-11,357), xlim = c(0,148)) +
  scale_x_continuous(expand = c(0, 0), breaks=c(1,40,80,120)) +
  scale_y_continuous(expand = c(0, 0), breaks=c(0,100,200,300)) +
  scale_fill_manual(values=sRNAcols) +
  theme_minimal() + 
  theme(panel.grid.major.x=element_blank(), panel.grid.minor.x=element_blank(), panel.grid.minor.y=element_blank()) +
  theme(axis.text.x=element_text(colour = "white"), axis.title.x= element_text(colour = "white")) +
  theme(axis.text.y = element_text(color="black",size=12)) +
  theme(axis.title.y = element_text(vjust=0.5,hjust=0.5,size=12)) +
  theme(legend.position="none") +
  theme(plot.margin = unit(c(0, 0, 0, 0), "cm")) +
  labs(fill="RNA Length") +
  geom_label(aes(x=125, y=275, label="aMIRA1",fontface="italic",size=21), fill = "white")

two <- ggplot() +
  geom_vline(xintercept = c(0,38,58,90,110),
             linetype=c("solid","dashed","dashed","dashed","dashed"),
             color=c("grey70","black","black","grey40","grey40"),
             size=c(0.8,0.6,0.6,0.6,0.6)) +
  geom_hline(yintercept = 0, color = "grey70", size = 0.4) +
  geom_col(data=cov_plus2,
           aes(x=Position, y=RPM,
               fill=factor(Length,levels=Lorder)), width=1) +
  geom_col(data=cov_minus2,
           aes(x=Position, y=RPM,
               fill=factor(Length, levels=Lorder)), width=1) +
  labs(x="Artificial microRNA gene position (nt)") +
  coord_cartesian(ylim = c(-6,300), xlim = c(0,148)) +
  scale_x_continuous(expand = c(0, 0), breaks=c(1,40,80,120)) + scale_y_continuous(expand = c(0, 0), breaks=c(0,100,200,300)) +
  scale_fill_manual(values=sRNAcols) +
  theme_minimal() +
  theme(panel.grid.major.x=element_blank(), panel.grid.minor.x=element_blank(), panel.grid.minor.y=element_blank()) +
  theme(axis.text.x=element_text(colour = "white"), axis.title.x= element_text(colour = "white")) +
  theme(axis.text.y = element_text(color="black",size=12)) +
  theme(axis.title.y = element_text(vjust=0.5,hjust=0.5,size=12)) +
  theme(legend.position="none") +
  theme(plot.margin = unit(c(0, 0, 0, 0), "cm")) +
  labs(fill="RNA Length") +
  geom_label(aes(x=125, y=230, label="aMIRA3",fontface="italic",size=21), fill = "white")

three <- ggplot() +
  geom_vline(xintercept = c(0,38,58,90,110),
             linetype=c("solid","dashed","dashed","dashed","dashed"),
             color=c("grey70","black","black","grey40","grey40"),
             size=c(0.8,0.6,0.6,0.6,0.6)) +
  geom_hline(yintercept = 0, color = "grey70", size = 0.4) +
  geom_col(data=cov_plus3,
           aes(x=Position, y=RPM,
               fill=factor(Length,levels=Lorder)), width=1) +
  geom_col(data=cov_minus3,
           aes(x=Position, y=RPM,
               fill=factor(Length, levels=Lorder)), width=1) +
  labs(x="Artificial microRNA gene position (nt)") +
  coord_cartesian(ylim = c(-9,350), xlim = c(0,148)) +
  scale_x_continuous(expand = c(0, 0), breaks=c(1,40,80,120)) + scale_y_continuous(expand = c(0, 0), breaks=c(0,100,200,300)) +
  scale_fill_manual(values=sRNAcols) +
  theme_minimal() +
  theme(panel.grid.major.x=element_blank(), panel.grid.minor.x=element_blank(), panel.grid.minor.y=element_blank()) +
  theme(axis.text.x=element_text(colour = "white"), axis.title.x= element_text(colour = "white")) +
  theme(axis.text.y = element_text(color="black",size=12)) +
  theme(axis.title.y = element_text(vjust=0.5,hjust=0.5,size=12)) +
  theme(legend.position="none") +
  theme(plot.margin = unit(c(0, 0, 0, 0), "cm")) +
  labs(fill="RNA Length") +
  geom_label(aes(x=125, y=275, label="aMIRA4",fontface="italic",size=21), fill = "white")

four <- ggplot() +
  geom_vline(xintercept = c(0,38,58,90,110),
             linetype=c("solid","dashed","dashed","dashed","dashed"),
             color=c("grey70","black","black","grey40","grey40"),
             size=c(0.8,0.6,0.6,0.6,0.6)) +
  geom_hline(yintercept = 0, color = "grey70", size = 0.4) +
  geom_col(data=cov_plus4,
           aes(x=Position, y=RPM,
               fill=factor(Length,levels=Lorder)), width=1) +
  geom_col(data=cov_minus4,
           aes(x=Position, y=RPM,
               fill=factor(Length, levels=Lorder)), width=1) +
  labs(x="Artificial microRNA gene position (nt)") +
  coord_cartesian(ylim = c(-7,55), xlim = c(0,148)) +
  scale_x_continuous(expand = c(0, 0), breaks=c(1,40,80,120)) + scale_y_continuous(expand = c(0, 0), breaks=c(0,15,30,45)) +
  scale_fill_manual(values=sRNAcols) +
  theme_minimal() +
  theme(panel.grid.major.x=element_blank(), panel.grid.minor.x=element_blank(), panel.grid.minor.y=element_blank()) +
  theme(axis.text.x=element_text(colour = "white"), axis.title.x= element_text(colour = "white")) +
  theme(axis.text.y = element_text(color="black",size=12)) +
  theme(axis.title.y = element_text(vjust=0.5,hjust=0.5,size=12)) +
  theme(legend.position="none") +
  theme(plot.margin = unit(c(0, 0, 0, 0), "cm")) +
  labs(fill="RNA Length") +
  geom_label(aes(x=125, y=40, label="aMIRB1",fontface="italic",size=21), fill = "white")

five <- ggplot() +
  geom_vline(xintercept = c(0,38,58,90,110),
             linetype=c("solid","dashed","dashed","dashed","dashed"),
             color=c("grey70","black","black","grey40","grey40"),
             size=c(0.8,0.6,0.6,0.6,0.6)) +
  geom_hline(yintercept = 0, color = "grey70", size = 0.4) +
  geom_col(data=cov_plus5,
           aes(x=Position, y=RPM,
               fill=factor(Length,levels=Lorder)), width=1) +
  geom_col(data=cov_minus5,
           aes(x=Position, y=RPM,
               fill=factor(Length, levels=Lorder)), width=1) +
  labs(x="Artificial microRNA gene position (nt)") +
  coord_cartesian(ylim = c(-12,850), xlim = c(0,148)) +
  scale_x_continuous(expand = c(0, 0), breaks=c(1,40,80,120)) + scale_y_continuous(expand = c(0, 0), breaks=c(0,250,500,750)) +
  scale_fill_manual(values=sRNAcols) +
  theme_minimal() +
  theme(panel.grid.major.x=element_blank(), panel.grid.minor.x=element_blank(), panel.grid.minor.y=element_blank()) +
  theme(axis.text.x=element_text(colour = "white"), axis.title.x= element_text(colour = "white")) +
  theme(axis.text.y = element_text(color="black",size=12)) +
  theme(axis.title.y = element_text(vjust=0.5,hjust=0.5,size=12)) +
  theme(legend.position="none") +
  theme(plot.margin = unit(c(0, 0, 0, 0), "cm")) +
  labs(fill="RNA Length") +
  geom_label(aes(x=125, y=630, label="aMIRB5",fontface="italic",size=21), fill = "white")

six <- ggplot() +
  geom_vline(xintercept = c(0,38,58,90,110),
             linetype=c("solid","dashed","dashed","dashed","dashed"),
             color=c("grey70","black","black","grey40","grey40"),
             size=c(0.8,0.6,0.6,0.6,0.6)) +
  geom_hline(yintercept = 0, color = "grey70", size = 0.4) +
  geom_col(data=cov_plus6,
           aes(x=Position, y=RPM,
               fill=factor(Length,levels=Lorder)), width=1) +
  geom_col(data=cov_minus6,
           aes(x=Position, y=RPM,
               fill=factor(Length, levels=Lorder)), width=1) +
  labs(x="Artificial microRNA gene position (nt)") +
  coord_cartesian(ylim = c(-14,50), xlim = c(0,148)) +
  scale_x_continuous(expand = c(0, 0), breaks=c(1,40,80,120)) + scale_y_continuous(expand = c(0, 0), breaks=c(0,15,30,45)) +
  scale_fill_manual(values=sRNAcols) +
  theme_minimal() +
  theme(panel.grid.major.x=element_blank(), panel.grid.minor.x=element_blank(), panel.grid.minor.y=element_blank()) +
  theme(axis.text.x=element_text(colour = "white"), axis.title.x= element_text(colour = "white")) +
  theme(axis.text.y = element_text(color="black",size=12)) +
  theme(axis.title.y = element_text(vjust=0.5,hjust=0.5,size=12)) +
  theme(legend.position="none") +
  theme(plot.margin = unit(c(0, 0, 0, 0), "cm")) +
  labs(fill="RNA Length") +
  geom_label(aes(x=125, y=36, label="aMIRC4",fontface="italic",size=21), fill = "white")

seven <- ggplot() +
  geom_vline(xintercept = c(0,38,58,90,110),
             linetype=c("solid","dashed","dashed","dashed","dashed"),
             color=c("grey70","black","black","grey40","grey40"),
             size=c(0.8,0.6,0.6,0.6,0.6)) +
  geom_hline(yintercept = 0, color = "grey70", size = 0.4) +
  geom_col(data=cov_plus7,
           aes(x=Position, y=RPM,
               fill=factor(Length,levels=Lorder)), width=1) +
  geom_col(data=cov_minus7,
           aes(x=Position, y=RPM,
               fill=factor(Length, levels=Lorder)), width=1) +
  labs(x="Artificial microRNA gene position (nt)") +
  coord_cartesian(ylim = c(-6,81), xlim = c(0,148)) +
  scale_x_continuous(expand = c(0, 0), breaks=c(1,40,80,120)) + scale_y_continuous(expand = c(0, 0), breaks=c(0,25,50,75)) +
  scale_fill_manual(values=sRNAcols) +
  theme_minimal() +
  theme(panel.grid.major.x=element_blank(), panel.grid.minor.x=element_blank(), panel.grid.minor.y=element_blank()) +
  theme(axis.text.x=element_text(colour = "white"), axis.title.x= element_text(colour = "white")) +
  theme(axis.text.y = element_text(color="black",size=12)) +
  theme(axis.title.y = element_text(vjust=0.5,hjust=0.5,size=12)) +
  theme(legend.position="none") +
  theme(plot.margin = unit(c(0, 0, 0, 0), "cm")) +
  labs(fill="RNA Length") +
  geom_label(aes(x=125, y=60, label="aMIRC5",fontface="italic",size=21), fill = "white")

eight <- ggplot() +
  geom_vline(xintercept = c(0,38,58,90,110),
             linetype=c("solid","dashed","dashed","dashed","dashed"),
             color=c("grey70","black","black","grey40","grey40"),
             size=c(0.8,0.6,0.6,0.6,0.6)) +
  geom_hline(yintercept = 0, color = "grey70", size = 0.4) +
  geom_col(data=cov_plus8,
           aes(x=Position, y=RPM,
               fill=factor(Length,levels=Lorder)), width=1) +
  geom_col(data=cov_minus8,
           aes(x=Position, y=RPM,
               fill=factor(Length, levels=Lorder)), width=1) +
  labs(x="Artificial microRNA gene position (nt)") +
  coord_cartesian(ylim = c(-10,120), xlim = c(0,148)) +
  scale_x_continuous(expand = c(0, 0), breaks=c(1,40,80,120)) + scale_y_continuous(expand = c(0, 0), breaks=c(0,40,80,120)) +
  scale_fill_manual(values=sRNAcols) +
  theme_minimal() +
  theme(panel.grid.major.x=element_blank(), panel.grid.minor.x=element_blank(), panel.grid.minor.y=element_blank()) +
  theme(axis.text.x=element_text(colour = "black",size=12), axis.title.x= element_text(colour = "black",size=12)) +
  theme(axis.text.y = element_text(color="black",size=12)) +
  theme(axis.title.y = element_text(vjust=0.5,hjust=0.5,size=12)) +
  theme(legend.position="none") +
  theme(plot.margin = unit(c(0, 0, 0, 0), "cm")) +
  labs(fill="RNA Length") +
  geom_label(aes(x=125, y=90, label="aMIRE1",fontface="italic",size=21), fill = "white")

# Merge graphs in one plot
merge2 <- ggarrange(title, zero, one, two, three, four, five, six, seven, eight,
          heights = c(0.2,0.7,1,1,1,1,1,1,1,1,0.2),
          ncol = 1, nrow = 10, align = "v")

merge2

# Add legend
legend <- get_legend(one + theme(legend.position="right", legend.direction = "vertical") +
                       guides(fill = guide_legend(ncol = 1)))

plot_grid(merge2, legend, rel_widths = c(3, .4))
```
