## Supplemental Tables, Code, and Data for "Expression and processing of polycistronic artificial microRNAs and *trans*-acting siRNAs in *Solanum lycopersicum* and *Nicotiana benthamiana*": Supplemental_Code_3.R

```
# This code generates Figure 4A

# Activate required non-base packages
library(ggplot2)
library(dplyr)
library(cowplot)
library(ggpubr)

# Set the working directory by replacing "/your/path" with
# the complete path to the directory where this file is
setwd("/your/path")

# Read files made by "Supplemental Code 8.p"
coverage1 <- read.table(file="pamiR1_tomato.txt",
                       sep="\t",head=TRUE)
coverage2 <- read.table(file="pamiR2_tomato.txt",
                       sep="\t",head=TRUE)
coverage3 <- read.table(file="pamiR3_tomato.txt",
                        sep="\t",head=TRUE)
coverage4 <- read.table(file="pamiR4_tomato.txt",
                        sep="\t",head=TRUE)
coverage5 <- read.table(file="aTAS3_tomato.txt",
                        sep="\t",head=TRUE)

# Read files determining the gene features
gene1 <- read.table(file="gene_pamiR1.tsv",
                        sep="\t",head=TRUE)
gene2 <- read.table(file="gene_pamiR2.tsv",
                    sep="\t",head=TRUE)
gene3 <- read.table(file="gene_pamiR3.tsv",
                    sep="\t",head=TRUE)
gene4 <- read.table(file="gene_pamiR4.tsv",
                    sep="\t",head=TRUE)
gene5 <- read.table(file="gene_aTAS3.tsv",
                    sep="\t",head=TRUE)

# Filter the plus and minus sets
cov_plus1 <- filter(coverage1, Strand=="plus")
cov_minus1 <- filter(coverage1, Strand=="minus")
cov_plus2 <- filter(coverage2, Strand=="plus")
cov_minus2 <- filter(coverage2, Strand=="minus")
cov_plus3 <- filter(coverage3, Strand=="plus")
cov_minus3 <- filter(coverage3, Strand=="minus")
cov_plus4 <- filter(coverage4, Strand=="plus")
cov_minus4 <- filter(coverage4, Strand=="minus")
cov_plus5 <- filter(coverage5, Strand=="plus")
cov_minus5 <- filter(coverage5, Strand=="minus")

# Define color pallette
sRNAcols = c("lightgray", # <20 nts
             "skyblue",   # 20 nts
             "blue",      # 21 nts
             "mediumseagreen", # 22nts
             "orange",    # 23 nts
             "tomato",    # 24 nts
             "darkgray")  # >24 nts

miRNAcols = c("grey", # gene
              "black", # amiRNA
              "grey40", # amiRNA*
              "lightpink") # tRNA-Gly
miRNAcolsTAS = c("grey", # gene
              "black", # amiRNA
              "grey40", # amiRNA*
              "khaki") #miR390-TS 

# Define the order to list the Length categories
Lorder = c("<20 nts","20 nts","21 nts","22 nts","23 nts","24 nts",">24 nts")
Lordem = c("gene","amiRNA","amiRNA*","tRNA-Gly")
LordemTAS = c("gene","amiRNA","amiRNA*","miR390-TS")

# Title
title <- ggdraw() + 
  draw_label(expression(bolditalic("paMIRNA")~bold("coverage in")~bolditalic("S. lycopersicum")),
             fontface = 'bold', hjust = 0.5)

# Make plots
cartoon1 <- ggplot() +
  geom_col(data=gene1,aes(x=Position, y=RPM, fill=factor(Type,levels=Lordem)), width=1) +
  scale_fill_manual(values=miRNAcols) +
  theme_minimal() +
  theme(panel.grid.major.x=element_blank(), panel.grid.minor.x=element_blank(), panel.grid.major.y=element_blank(),panel.grid.minor.y=element_blank()) +
  coord_cartesian(ylim = c(0, 10), xlim = c(0,322)) +
  scale_x_continuous(expand = c(0, 0), breaks=c(1,100,200,300)) + scale_y_continuous(expand = c(0, 0)) +
  theme(axis.text.y = element_text(color="white"), axis.text.x=element_blank()) +
  theme(axis.title.y = element_text(colour="white")) + 
  theme(axis.title.x = element_text(colour="white")) +
  theme(legend.position="none") +
  theme(plot.margin = unit(c(0, 0, 0, 0), "cm")) +
  labs(fill="RNA Length") +
  annotate("text", x=c(28,81,135,188,242,295), y=c(5,5,5,5,5,5), label=c("amiRA1","amiRA1*","amiRA3","amiRA3*","amiRA4","amiRA4*"), size=4)

plot1 <- ggplot() +
  geom_vline(xintercept = c(0,18,38,70,90,125,145,177,197,232,252,284,304),
             linetype=c("solid","dashed","dashed","dashed","dashed","dashed","dashed","dashed","dashed","dashed","dashed","dashed","dashed"),
             color=c("grey70","black","black","grey40","grey40","black","black","grey40","grey40","black","black","grey40","grey40"),
             size=c(0.8,0.6,0.6,0.6,0.6,0.6,0.6,0.6,0.6,0.6,0.6,0.6,0.6)) +
  geom_hline(yintercept = 0, color = "grey70", size = 0.4) +
  geom_col(data=cov_plus1, aes(x=Position, y=RPM, fill=factor(Length,levels=Lorder)), width=1) +
  geom_col(data=cov_minus1, aes(x=Position, y=RPM, fill=factor(Length, levels=Lorder)), width=1) +
  labs(x="Artificial microRNA gene position (nt)") +
  coord_cartesian(ylim = c(-5,60), xlim = c(0,322)) +
  scale_x_continuous(expand = c(0, 0), breaks=c(1,100,200,300)) +
  scale_y_continuous(expand = c(0, 0), breaks=c(0,20,40,60)) +
  scale_fill_manual(values=sRNAcols) +
  theme_minimal() + 
  theme(panel.grid.major.x=element_blank(), panel.grid.minor.x=element_blank(), panel.grid.minor.y=element_blank()) +
  theme(axis.text.x=element_text(colour="black",size=12), axis.title.x= element_text(colour = "white")) +
  theme(axis.text.y=element_text(colour="black",size=12), axis.title.y = element_text(vjust=1,size=12)) +
  theme(legend.position="none") +
  theme(plot.margin = unit(c(0, 0, 0, 0), "cm")) +
  labs(fill="RNA Length")

cartoon2 <- ggplot() +
  geom_col(data=gene2, aes(x=Position, y=RPM, fill=factor(Type,levels=Lordem)), width=1) +
  scale_fill_manual(values=miRNAcols) +
  theme_minimal() +
  theme(panel.grid.major.x=element_blank(), panel.grid.minor.x=element_blank(), panel.grid.major.y=element_blank(),panel.grid.minor.y=element_blank()) +
  coord_cartesian(ylim = c(0, 10), xlim = c(0,673)) +
  scale_x_continuous(expand = c(0, 0), breaks=c(1,100,200,300,400,500,600)) + scale_y_continuous(expand = c(0, 0)) +
  theme(axis.text.y = element_text(color="white"), axis.text.x=element_blank()) +
  theme(axis.title.y = element_text(colour="white")) + 
  theme(axis.title.x = element_text(colour="white")) +
  theme(legend.position="none") +
  theme(plot.margin = unit(c(0, 0, 0, 0), "cm")) +
  labs(fill="RNA Length") +
  annotate("text", x=c(39,125,177,264,349,401,488,573,625), y=c(5,5,5,5,5,5,5,5,5), label=c("tRNA-Gly","amiRA1","amiRA1*","tRNA-Gly","amiRA4","amiRA4*","tRNA-Gly","amiRB1","amiRB1*"), size=4)

plot2 <- ggplot() +
  geom_vline(xintercept = c(0,1,77,115,135,167,187,225,301,339,359,391,411,449,525,563,583,615,635),
             linetype=c("solid","dashed","dashed","dashed","dashed","dashed","dashed","dashed","dashed","dashed","dashed","dashed","dashed","dashed","dashed","dashed","dashed","dashed","dashed"),
             color=c("grey70","lightpink","lightpink","black","black","grey40","grey40","lightpink","lightpink","black","black","grey40","grey40","lightpink","lightpink","black","black","grey40","grey40"),
             size=c(0.8,0.6,0.6,0.6,0.6,0.6,0.6,0.6,0.6,0.6,0.6,0.6,0.6,0.6,0.6,0.6,0.6,0.6,0.6)) +
  geom_hline(yintercept = 0, color = "grey70", size = 0.4) +
  geom_col(data=cov_plus2, aes(x=Position, y=RPM, fill=factor(Length,levels=Lorder)), width=1) +
  geom_col(data=cov_minus2, aes(x=Position, y=RPM, fill=factor(Length, levels=Lorder)), width=1) +
  labs(x="Artificial microRNA gene position (nt)") +
  coord_cartesian(ylim = c(-5,450), xlim = c(0,673)) +
  scale_x_continuous(expand = c(0, 0), breaks=c(1,100,200,300,400,500,600)) +
  scale_y_continuous(expand = c(0, 0), breaks=c(0,150,300,450)) +
  scale_fill_manual(values=sRNAcols) +
  theme_minimal() + 
  theme(panel.grid.major.x=element_blank(), panel.grid.minor.x=element_blank(), panel.grid.minor.y=element_blank()) +
  theme(axis.text.x=element_text(colour="black",size=12), axis.title.x= element_text(colour = "white")) +
  theme(axis.text.y=element_text(colour="black",size=12),axis.title.y = element_text(vjust=0.1,size=12)) +
  theme(legend.position="none") +
  theme(plot.margin = unit(c(0, 0, 0, 0), "cm")) +
  labs(fill="RNA Length") 

cartoon3 <- ggplot() +
  geom_col(data=gene3, aes(x=Position, y=RPM, fill=factor(Type,levels=Lordem)), width=1) +
  scale_fill_manual(values=miRNAcols) +
  theme_minimal() +
  theme(panel.grid.major.x=element_blank(), panel.grid.minor.x=element_blank(), panel.grid.major.y=element_blank(),panel.grid.minor.y=element_blank()) +
  coord_cartesian(ylim = c(0, 10), xlim = c(0,415)) +
  scale_x_continuous(expand = c(0, 0), breaks=c(1,100,200,300,400)) + scale_y_continuous(expand = c(0, 0)) +
  theme(axis.text.y = element_text(color="white"), axis.text.x=element_blank()) +
  theme(axis.title.y = element_text(colour="white")) + 
  theme(axis.title.x = element_text(colour="white")) +
  theme(legend.position="none") +
  theme(plot.margin = unit(c(0, 0, 0, 0), "cm")) +
  labs(fill="RNA Length") +
  annotate("text", x=c(44,99,182,237,320,375), y=c(5,5,5,5,5,5), label=c("amiRA1*","amiRA1","amiRA4*","amiRA4","amiRC5*","amiRC5"), size=4)

plot3 <- ggplot() +
  geom_vline(xintercept = c(0,34,54,89,109,172,192,227,247,310,330,365,385),
             linetype=c("solid","dashed","dashed","dashed","dashed","dashed","dashed","dashed","dashed","dashed","dashed","dashed","dashed"),
             color=c("grey70","grey40","grey40","black","black","grey40","grey40","black","black","grey40","grey40","black","black"),
             size=c(0.8,0.6,0.6,0.6,0.6,0.6,0.6,0.6,0.6,0.6,0.6,0.6,0.6)) +
  geom_hline(yintercept = 0, color = "grey70", size = 0.4) +
  geom_col(data=cov_plus3, aes(x=Position, y=RPM, fill=factor(Length,levels=Lorder)), width=1) +
  geom_col(data=cov_minus3, aes(x=Position, y=RPM, fill=factor(Length, levels=Lorder)), width=1) +
  labs(x="Artificial microRNA gene position (nt)") +
  coord_cartesian(ylim = c(-11,450), xlim = c(0,415)) +
  scale_x_continuous(expand = c(0, 0), breaks=c(1,100,200,300,400)) +
  scale_y_continuous(expand = c(0, 0), breaks=c(0,150,300,450)) +
  scale_fill_manual(values=sRNAcols) +
  theme_minimal() + 
  theme(panel.grid.major.x=element_blank(), panel.grid.minor.x=element_blank(), panel.grid.minor.y=element_blank()) +
  theme(axis.text.x=element_text(colour="black",size=12), axis.title.x= element_text(colour = "white")) +
  theme(axis.text.y=element_text(colour="black",size=12), axis.title.y = element_text(vjust=0.1,size=12)) +
  theme(legend.position="none") +
  theme(plot.margin = unit(c(0, 0, 0, 0), "cm")) +
  labs(fill="RNA Length") 

cartoon4 <- ggplot() +
  geom_col(data=gene4, aes(x=Position, y=RPM, fill=factor(Type,levels=Lordem)), width=1) +
  scale_fill_manual(values=miRNAcols) +
  theme_minimal() +
  theme(panel.grid.major.x=element_blank(), panel.grid.minor.x=element_blank(), panel.grid.major.y=element_blank(),panel.grid.minor.y=element_blank()) +
  coord_cartesian(ylim = c(0, 10), xlim = c(0,765)) +
  scale_x_continuous(expand = c(0, 0), breaks=c(1,100,200,300,400,500,600,700)) + scale_y_continuous(expand = c(0, 0)) +
  theme(axis.text.y = element_text(color="white"), axis.text.x=element_blank()) +
  theme(axis.title.y = element_text(colour="white")) + 
  theme(axis.title.x = element_text(colour="white")) +
  theme(legend.position="none") +
  theme(plot.margin = unit(c(0, 0, 0, 0), "cm")) +
  labs(fill="RNA Length") +
  annotate("text", x=c(43,142,254,309,392,450,531,588,674,731), y=c(5,5,5,5,5,5,5,5,5,5), label=c("amiRA1*","amiRA1","amiRA4*","amiRA4","amiRB5*","amiRB5","amiRC4*","amiRC4","amiRC5*","amiRC5"), size=4)

plot4 <- ggplot() +
  geom_vline(xintercept = c(0,33,55,132,152,249,270,292,312,387,408,433,453,526,547,571,591,669,690,714,734),
             linetype=c("solid","dashed","dashed","dashed","dashed","dashed","dashed","dashed","dashed","dashed","dashed","dashed","dashed","dashed","dashed","dashed","dashed","dashed","dashed","dashed","dashed"),
             color=c("grey70","grey40","grey40","black","black","grey40","grey40","black","black","grey40","grey40","black","black","grey40","grey40","black","black","grey40","grey40","black","black"),
             size=c(0.8,0.6,0.6,0.6,0.6,0.6,0.6,0.6,0.6,0.6,0.6,0.6,0.6,0.6,0.6,0.6,0.6,0.6,0.6,0.6,0.6)) +
  geom_hline(yintercept = 0, color = "grey70", size = 0.4) +
  geom_col(data=cov_plus4, aes(x=Position, y=RPM, fill=factor(Length,levels=Lorder)), width=1) +
  geom_col(data=cov_minus4, aes(x=Position, y=RPM, fill=factor(Length, levels=Lorder)), width=1) +
  labs(x="Artificial microRNA gene position (nt)") +
  coord_cartesian(ylim = c(-50,250), xlim = c(0,765)) +
  scale_x_continuous(expand = c(0, 0), breaks=c(1,100,200,300,400,500,600,700)) +
  scale_y_continuous(expand = c(0, 0), breaks=c(0,50,150,250)) +
  scale_fill_manual(values=sRNAcols) +
  theme_minimal() + 
  theme(panel.grid.major.x=element_blank(), panel.grid.minor.x=element_blank(), panel.grid.minor.y=element_blank()) +
  theme(axis.text.x=element_text(colour="black",size=12), axis.title.x= element_text(colour = "white")) +
  theme(axis.text.y=element_text(colour="black",size=12), axis.title.y = element_text(vjust=0.1,size=12)) +
  theme(legend.position="none") +
  theme(plot.margin = unit(c(0, 0, 0, 0), "cm")) +
  labs(fill="RNA Length") 

cartoon5 <- ggplot() +
  geom_col(data=gene5, aes(x=Position, y=RPM, fill=factor(Type,levels=LordemTAS)), width=1) +
  scale_fill_manual(values=miRNAcolsTAS) +
  theme_minimal() +
  theme(panel.grid.major.x=element_blank(), panel.grid.minor.x=element_blank(), panel.grid.major.y=element_blank(),panel.grid.minor.y=element_blank()) +
  coord_cartesian(ylim = c(0, 10), xlim = c(0,349)) +
  scale_x_continuous(expand = c(0, 0), breaks=c(1,100,200,300)) + scale_y_continuous(expand = c(0, 0)) +
  theme(axis.text.y = element_text(color="white"), axis.text.x=element_blank()) +
  theme(axis.title.y = element_text(colour="white")) + 
  theme(axis.title.x = element_text(colour="white")) +
  theme(legend.position="none") +
  theme(plot.margin = unit(c(0, 0, 0, 0), "cm")) +
  labs(fill="RNA Length") +
  annotate("text", x=c(46,104,125,146,167,188,209,230,251,303), y=c(5,5,7.5,5,7.5,5,7.5,5,7.5,5), label=c("miR390-TS","amiRA3","amiRA1","amiRA4","amiRC5","amiRB5","amiRC4","amiRE1","amiRB1","miR390-TS"), size=4)

plot5 <- ggplot() +
  geom_vline(xintercept = c(0,36,56,94,114,115,135,136,156,157,177,178,198,199,219,220,240,241,261,293,313),
             linetype=c("solid","dashed","dashed","dashed","dashed","dashed","dashed","dashed","dashed","dashed","dashed","dashed","dashed","dashed","dashed","dashed","dashed","dashed","dashed","dashed","dashed"),
             color=c("grey70","khaki","khaki","black","black","grey40","grey40","black","black","grey40","grey40","black","black","grey40","grey40","black","black","grey40","grey40","khaki","khaki"),
             size=c(0.8,0.6,0.6,0.6,0.6,0.6,0.6,0.6,0.6,0.6,0.6,0.6,0.6,0.6,0.6,0.6,0.6,0.6,0.6,0.6,0.6)) +
  geom_hline(yintercept = 0, color = "grey70", size = 0.4) +
  geom_col(data=cov_plus5, aes(x=Position, y=RPM, fill=factor(Length,levels=Lorder)), width=1) +
  geom_col(data=cov_minus5, aes(x=Position, y=RPM, fill=factor(Length, levels=Lorder)), width=1) +
  labs(x="Artificial microRNA gene position (nt)") +
  coord_cartesian(ylim = c(-25,40), xlim = c(0,349)) +
  scale_x_continuous(expand = c(0, 0), breaks=c(1,100,200,300)) +
  scale_y_continuous(expand = c(0, 0), breaks=c(-20,0,20,40)) +
  scale_fill_manual(values=sRNAcols) +
  theme_minimal() + 
  theme(panel.grid.major.x=element_blank(), panel.grid.minor.x=element_blank(), panel.grid.minor.y=element_blank()) +
  theme(axis.text.x=element_text(colour="black",size=12), axis.title.x= element_text(colour = "black",size=12)) +
  theme(axis.text.y=element_text(colour="black",size=12), axis.title.y = element_text(vjust=0.1,size=12)) +
  theme(legend.position="none") +
  theme(plot.margin = unit(c(0, 0, 0, 0), "cm")) +
  labs(fill="RNA Length") 

# Merge graphs in one plot
merge <- ggarrange(title,cartoon1,plot1,cartoon2,plot2,cartoon3,plot3,cartoon4,plot4,cartoon5,plot5,
          heights = c(0.2,0.7,1,0.7,1,0.7,1,0.7,1,0.7,1),
          ncol = 1, nrow = 11, align = "v",
          labels = c("","paMIR1","","paMIR2","","paMIR3","","paMIR4","","aTAS3"),
          font.label = list(size = 13, face="bold.italic"), hjust=0, vjust=0.7)

merge

# Add legend
legend <- get_legend(one + theme(legend.position="right", legend.direction = "vertical") +
                       guides(fill = guide_legend(ncol = 1)))

plot_grid(merge, legend, rel_widths = c(3, .4))
```
