## Supplemental Tables, Code, and Data for "Expression and processing of polycistronic artificial microRNAs and *trans*-acting siRNAs in *Solanum lycopersicum* and *Nicotiana benthamiana*": Supplemental_Code_4.R

```
# This code generates Figure 5A

# Activate required non-base packages
library(ggplot2)
library(dplyr)
library(cowplot)
library(ggpubr)

# Set the working directory by replacing "/your/path" with
# the complete path to the directory where this file is
setwd("/your/path")

# Read in a file made by sRNA_coverage.pl
coverage1 <- read.table(file="5-prime-end_pamiR1_tomato.txt", sep="\t", head=TRUE)
coverage2 <- read.table(file="5-prime-end_pamiR2_tomato.txt", sep="\t", head=TRUE)
coverage3 <- read.table(file="5-prime-end_pamiR3_tomato.txt", sep="\t", head=TRUE)
coverage4 <- read.table(file="5-prime-end_pamiR4_tomato.txt", sep="\t", head=TRUE)

# Define color pallette
sRNAcols = c("lightgray", # <20 nts
             "skyblue",   # 20 nts
             "blue",      # 21 nts
             "mediumseagreen", # 22nts
             "orange",    # 23 nts
             "tomato",    # 24 nts
             "darkgray")  # >24 nts

# Define the order to list the Length categories
Lorder = c("<20 nts","20 nts","21 nts","22 nts","23 nts","24 nts",">24 nts")

# Make plot paMIR1
paMIR1_1 <- ggplot() +
  geom_segment(aes(x = 18, y = -1, xend = 18, yend = 30), linetype = "dashed", colour="grey70") +
  geom_hline(yintercept = 0, color = "grey70", size = 0.4) +
  geom_col(data=coverage1,aes(x=Position, y=RPM, fill=factor(Length,levels=Lorder)),width=0.9) +
  scale_x_continuous(breaks=c(13,15,18,21,23), labels=c("-5","-3","0","+3","+5")) +
  scale_y_continuous(expand = c(0, 0), breaks=c(0,10,20,30)) +
  scale_fill_manual(values=sRNAcols) +
  labs(x="amiRA1", y="RPM") +
  coord_cartesian(xlim = c(13,23), ylim = c(-1,37)) +
  theme_minimal() +
  theme(panel.grid.major.x=element_blank(), panel.grid.minor.x=element_blank(), panel.grid.minor.y=element_blank()) +
  theme(axis.text.x=element_text(colour = "black",size=12), axis.title.x = element_text(color="black",size=12)) +
  theme(axis.text.y=element_text(colour = "black",size=12), axis.title.y = element_text(color="black",vjust=1,size=12)) +
  theme(legend.position="none") +
  theme(plot.margin = unit(c(0, 0, 0, 0), "cm")) +
  labs(fill="RNA Length")

paMIR1_2 <- ggplot() +
  geom_segment(aes(x = 70, y = -1, xend = 70, yend = 30), linetype = "dashed", colour="grey70") +
  geom_hline(yintercept = 0, color = "grey70", size = 0.4) +
  geom_col(data=coverage1, aes(x=Position, y=RPM, fill=factor(Length,levels=Lorder)),width=0.9) +
  scale_x_continuous(breaks=c(65,67,70,73,75), labels=c("-5","-3","0","+3","+5")) +
  scale_y_continuous(expand = c(0, 0), breaks=c(0,10,20,30)) +
  scale_fill_manual(values=sRNAcols) +
  labs(x="amiRA1*") +
  coord_cartesian(xlim = c(65,75), ylim = c(-1,37)) +
  theme_minimal() +
  theme(panel.grid.major.x=element_blank(), panel.grid.minor.x=element_blank(), panel.grid.minor.y=element_blank()) +
  theme(axis.text.x=element_text(colour = "black",size=12), axis.title.x = element_text(color="grey40",size=12)) +
  theme(axis.text.y=element_text(colour = "white",size=12), axis.title.y = element_text(color="white",vjust=1,size=12)) +
  theme(legend.position="none") +
  theme(plot.margin = unit(c(0, 0.1, 0, -0.1), "cm")) +
  labs(fill="RNA Length")

paMIR1_3 <- ggplot() +
  geom_segment(aes(x = 125, y = -1, xend = 125, yend = 30), linetype = "dashed", colour="grey70") +
  geom_hline(yintercept = 0, color = "grey70", size = 0.4) +
  geom_col(data=coverage1, aes(x=Position, y=RPM, fill=factor(Length,levels=Lorder)),width=0.9) +
  scale_x_continuous(breaks=c(120,122,125,128,130), labels=c("-5","-3","0","+3","+5")) +
  scale_y_continuous(expand = c(0, 0), breaks=c(0,10,20,30)) +
  scale_fill_manual(values=sRNAcols) +
  labs(x="amiRA3") +
  coord_cartesian(xlim = c(120,130), ylim = c(-1,37)) +
  theme_minimal() +
  theme(panel.grid.major.x=element_blank(), panel.grid.minor.x=element_blank(), panel.grid.minor.y=element_blank()) +
  theme(axis.text.x=element_text(colour = "black",size=12), axis.title.x = element_text(color="black",size=12)) +
  theme(axis.text.y=element_text(colour = "white",size=12), axis.title.y = element_text(color="white",vjust=1,size=12)) +
  theme(legend.position="none") +
  theme(plot.margin = unit(c(0, 0.1, 0, -0.1), "cm")) +
  labs(fill="RNA Length")

paMIR1_4 <- ggplot() +
  geom_segment(aes(x = 177, y = -1, xend = 177, yend = 30), linetype = "dashed", colour="grey70") +
  geom_hline(yintercept = 0, color = "grey70", size = 0.4) +
  geom_col(data=coverage1, aes(x=Position, y=RPM, fill=factor(Length,levels=Lorder)),width=0.9) +
  scale_x_continuous(breaks=c(172,174,177,180,182), labels=c("-5","-3","0","+3","+5")) +
  scale_y_continuous(expand = c(0, 0), breaks=c(0,10,20,30)) +
  scale_fill_manual(values=sRNAcols) +
  labs(x="amiRA3*") +
  coord_cartesian(xlim = c(172,182), ylim = c(-1,37)) +
  theme_minimal() +
  theme(panel.grid.major.x=element_blank(), panel.grid.minor.x=element_blank(), panel.grid.minor.y=element_blank()) +
  theme(axis.text.x=element_text(colour = "black",size=12), axis.title.x = element_text(color="grey40",size=12)) +
  theme(axis.text.y=element_text(colour = "white",size=12), axis.title.y = element_text(color="white",vjust=1,size=12)) +
  theme(legend.position="none") +
  theme(plot.margin = unit(c(0, 0.1, 0, -0.1), "cm")) +
  labs(fill="RNA Length")

paMIR1_5 <- ggplot() +
  geom_segment(aes(x = 232, y = -1, xend = 232, yend = 30), linetype = "dashed", colour="grey70") +
  geom_hline(yintercept = 0, color = "grey70", size = 0.4) +
  geom_col(data=coverage1, aes(x=Position, y=RPM, fill=factor(Length,levels=Lorder)),width=0.9) +
  scale_x_continuous(breaks=c(227,229,232,235,237), labels=c("-5","-3","0","+3","+5")) +
  scale_y_continuous(expand = c(0, 0), breaks=c(0,10,20,30)) +
  scale_fill_manual(values=sRNAcols) +
  labs(x="amiRA4") +
  coord_cartesian(xlim = c(227,237), ylim = c(-1,37)) +
  theme_minimal() +
  theme(panel.grid.major.x=element_blank(), panel.grid.minor.x=element_blank(), panel.grid.minor.y=element_blank()) +
  theme(axis.text.x=element_text(colour = "black",size=12), axis.title.x = element_text(color="black",size=12)) +
  theme(axis.text.y=element_text(colour = "white",size=12), axis.title.y = element_text(color="white",vjust=1,size=12)) +
  theme(legend.position="none") +
  theme(plot.margin = unit(c(0, 0.1, 0, -0.1), "cm")) +
  labs(fill="RNA Length")

paMIR1_6 <- ggplot() +
  geom_segment(aes(x = 284, y = -1, xend = 284, yend = 30), linetype = "dashed", colour="grey70") +
  geom_hline(yintercept = 0, color = "grey70", size = 0.4) +
  geom_col(data=coverage1, aes(x=Position, y=RPM, fill=factor(Length,levels=Lorder)),width=0.9) +
  scale_x_continuous(breaks=c(279,281,284,287,289), labels=c("-5","-3","0","+3","+5")) +
  scale_y_continuous(expand = c(0, 0), breaks=c(0,10,20,30)) +
  scale_fill_manual(values=sRNAcols) +
  labs(x="amiRA4*") +
  coord_cartesian(xlim = c(279,289), ylim = c(-1,37)) +
  theme_minimal() +
  theme(panel.grid.major.x=element_blank(), panel.grid.minor.x=element_blank(), panel.grid.minor.y=element_blank()) +
  theme(axis.text.x=element_text(colour = "black",size=12), axis.title.x = element_text(color="grey40",size=12)) +
  theme(axis.text.y=element_text(colour = "white",size=12), axis.title.y = element_text(color="white",vjust=1,size=12)) +
  theme(legend.position="none") +
  theme(plot.margin = unit(c(0, 0.1, 0, -0.1), "cm")) +
  labs(fill="RNA Length")

merge1 <- ggarrange(paMIR1_1,paMIR1_2,paMIR1_3,paMIR1_4,paMIR1_5,paMIR1_6,
                    widths = c(1,1,1,1,1,1),
                    ncol = 6, nrow = 1, align = "h")

# Make plot paMIR2
paMIR2_1 <- ggplot() +
  geom_segment(aes(x = 115, y = -2, xend = 115, yend = 75), linetype = "dashed", colour="grey70") +
  geom_hline(yintercept = 0, color = "grey70", size = 0.4) +
  geom_col(data=coverage2, aes(x=Position, y=RPM, fill=factor(Length,levels=Lorder)),width=0.9) +
  scale_x_continuous(breaks=c(110,112,115,118,120), labels=c("-5","-3","0","+3","+5")) +
  scale_y_continuous(expand = c(0, 0), breaks=c(0,25,50,75)) +
  scale_fill_manual(values=sRNAcols) +
  labs(x="amiRA1", y="RPM") +
  coord_cartesian(xlim = c(110,120), ylim = c(-2,95)) +
  theme_minimal() +
  theme(panel.grid.major.x=element_blank(), panel.grid.minor.x=element_blank(), panel.grid.minor.y=element_blank()) +
  theme(axis.text.x=element_text(colour = "black",size=12), axis.title.x = element_text(color="black",size=12)) +
  theme(axis.text.y=element_text(colour = "black",size=12), axis.title.y = element_text(color="black",vjust=1,size=12)) +
  theme(legend.position="none") +
  theme(plot.margin = unit(c(0, 0, 0, 0), "cm")) +
  labs(fill="RNA Length")

paMIR2_2 <- ggplot() +
  geom_segment(aes(x = 167, y = -2, xend = 167, yend = 75), linetype = "dashed", colour="grey70") +
  geom_hline(yintercept = 0, color = "grey70", size = 0.4) +
  geom_col(data=coverage2, aes(x=Position, y=RPM, fill=factor(Length,levels=Lorder)),width=0.9) +
  scale_x_continuous(breaks=c(162,164,167,170,172), labels=c("-5","-3","0","+3","+5")) +
  scale_y_continuous(expand = c(0, 0), breaks=c(0,25,50,75)) +
  scale_fill_manual(values=sRNAcols) +
  labs(x="amiRA1*") +
  coord_cartesian(xlim = c(162,172), ylim = c(-2,95)) +
  theme_minimal() +
  theme(panel.grid.major.x=element_blank(), panel.grid.minor.x=element_blank(), panel.grid.minor.y=element_blank()) +
  theme(axis.text.x=element_text(colour = "black",size=12), axis.title.x = element_text(color="grey40",size=12)) +
  theme(axis.text.y=element_text(colour = "white",size=12), axis.title.y = element_text(color="white",vjust=1,size=12)) +
  theme(legend.position="none") +
  theme(plot.margin = unit(c(0, 0.1, 0, -0.1), "cm")) +
  labs(fill="RNA Length")

paMIR2_3 <- ggplot() +
  geom_segment(aes(x = 339, y = -2, xend = 339, yend = 75), linetype = "dashed", colour="grey70") +
  geom_hline(yintercept = 0, color = "grey70", size = 0.4) +
  geom_col(data=coverage2, aes(x=Position, y=RPM, fill=factor(Length,levels=Lorder)),width=0.9) +
  scale_x_continuous(breaks=c(334,336,339,342,344), labels=c("-5","-3","0","+3","+5")) +
  scale_y_continuous(expand = c(0, 0), breaks=c(0,25,50,75)) +
  scale_fill_manual(values=sRNAcols) +
  labs(x="amiRA4") +
  coord_cartesian(xlim = c(334,344), ylim = c(-2,95)) +
  theme_minimal() +
  theme(panel.grid.major.x=element_blank(), panel.grid.minor.x=element_blank(), panel.grid.minor.y=element_blank()) +
  theme(axis.text.x=element_text(colour = "black",size=12), axis.title.x = element_text(color="black",size=12)) +
  theme(axis.text.y=element_text(colour = "white",size=12), axis.title.y = element_text(color="white",vjust=1,size=12)) +
  theme(legend.position="none") +
  theme(plot.margin = unit(c(0, 0.1, 0, -0.1), "cm")) +
  labs(fill="RNA Length")

paMIR2_4 <- ggplot() +
  geom_segment(aes(x = 391, y = -2, xend = 391, yend = 75), linetype = "dashed", colour="grey70") +
  geom_hline(yintercept = 0, color = "grey70", size = 0.4) +
  geom_col(data=coverage2, aes(x=Position, y=RPM, fill=factor(Length,levels=Lorder)),width=0.9) +
  scale_x_continuous(breaks=c(386,388,391,394,396), labels=c("-5","-3","0","+3","+5")) +
  scale_y_continuous(expand = c(0, 0), breaks=c(0,25,50,75)) +
  scale_fill_manual(values=sRNAcols) +
  labs(x="amiRA4*") +
  coord_cartesian(xlim = c(386,396), ylim = c(-2,95)) +
  theme_minimal() +
  theme(panel.grid.major.x=element_blank(), panel.grid.minor.x=element_blank(), panel.grid.minor.y=element_blank()) +
  theme(axis.text.x=element_text(colour = "black",size=12), axis.title.x = element_text(color="grey40",size=12)) +
  theme(axis.text.y=element_text(colour = "white",size=12), axis.title.y = element_text(color="white",vjust=1,size=12)) +
  theme(legend.position="none") +
  theme(plot.margin = unit(c(0, 0.1, 0, -0.1), "cm")) +
  labs(fill="RNA Length")

paMIR2_5 <- ggplot() +
  geom_segment(aes(x = 563, y = -2, xend = 563, yend = 75), linetype = "dashed", colour="grey70") +
  geom_hline(yintercept = 0, color = "grey70", size = 0.4) +
  geom_col(data=coverage2, aes(x=Position, y=RPM, fill=factor(Length,levels=Lorder)),width=0.9) +
  scale_x_continuous(breaks=c(558,560,563,566,568), labels=c("-5","-3","0","+3","+5")) +
  scale_y_continuous(expand = c(0, 0), breaks=c(0,25,50,75)) +
  scale_fill_manual(values=sRNAcols) +
  labs(x="amiRB1") +
  coord_cartesian(xlim = c(558,568), ylim = c(-2,95)) +
  theme_minimal() +
  theme(panel.grid.major.x=element_blank(), panel.grid.minor.x=element_blank(), panel.grid.minor.y=element_blank()) +
  theme(axis.text.x=element_text(colour = "black",size=12), axis.title.x = element_text(color="black",size=12)) +
  theme(axis.text.y=element_text(colour = "white",size=12), axis.title.y = element_text(color="white",vjust=1,size=12)) +
  theme(legend.position="none") +
  theme(plot.margin = unit(c(0, 0.1, 0, -0.1), "cm")) +
  labs(fill="RNA Length")

paMIR2_6 <- ggplot() +
  geom_segment(aes(x = 615, y = -2, xend = 615, yend = 75), linetype = "dashed", colour="grey70") +
  geom_hline(yintercept = 0, color = "grey70", size = 0.4) +
  geom_col(data=coverage2, aes(x=Position, y=RPM, fill=factor(Length,levels=Lorder)),width=0.9) +
  scale_x_continuous(breaks=c(610,612,615,618,620), labels=c("-5","-3","0","+3","+5")) +
  scale_y_continuous(expand = c(0, 0), breaks=c(0,25,50,75)) +
  scale_fill_manual(values=sRNAcols) +
  labs(x="amiRB1*") +
  coord_cartesian(xlim = c(610,620), ylim = c(-2,95)) +
  theme_minimal() +
  theme(panel.grid.major.x=element_blank(), panel.grid.minor.x=element_blank(), panel.grid.minor.y=element_blank()) +
  theme(axis.text.x=element_text(colour = "black",size=12), axis.title.x = element_text(color="grey40",size=12)) +
  theme(axis.text.y=element_text(colour = "white",size=12), axis.title.y = element_text(color="white",vjust=1,size=12)) +
  theme(legend.position="none") +
  theme(plot.margin = unit(c(0, 0.1, 0, -0.1), "cm")) +
  labs(fill="RNA Length")

merge2 <- ggarrange(paMIR2_1,paMIR2_2,paMIR2_3,paMIR2_4,paMIR2_5,paMIR2_6,
                    widths = c(1,1,1,1,1,1),
                    ncol = 6, nrow = 1, align = "h")

# Make plot paMIR3
paMIR3_1 <- ggplot() +
  geom_segment(aes(x = 34, y = -4, xend = 34, yend = 100), linetype = "dashed", colour="grey70") +
  geom_hline(yintercept = 0, color = "grey70", size = 0.4) +
  geom_col(data=coverage3, aes(x=Position, y=RPM, fill=factor(Length,levels=Lorder)),width=0.9) +
  scale_x_continuous(breaks=c(29,31,34,37,39), labels=c("-5","-3","0","+3","+5")) +
  scale_y_continuous(expand = c(0, 0), breaks=c(0,30,60,90)) +
  scale_fill_manual(values=sRNAcols) +
  labs(x="amiRA1*", y="RPM") +
  coord_cartesian(xlim = c(29,39), ylim = c(-4,120)) +
  theme_minimal() +
  theme(panel.grid.major.x=element_blank(), panel.grid.minor.x=element_blank(), panel.grid.minor.y=element_blank()) +
  theme(axis.text.x=element_text(colour = "black",size=12), axis.title.x = element_text(color="grey40",size=12)) +
  theme(axis.text.y=element_text(colour = "black",size=12), axis.title.y = element_text(color="black",vjust=1,size=12)) +
  theme(legend.position="none") +
  theme(plot.margin = unit(c(0, 0, 0, 0), "cm")) +
  labs(fill="RNA Length")

paMIR3_2 <- ggplot() +
  geom_segment(aes(x = 89, y = -4, xend = 89, yend = 100), linetype = "dashed", colour="grey70") +
  geom_hline(yintercept = 0, color = "grey70", size = 0.4) +
  geom_col(data=coverage3, aes(x=Position, y=RPM, fill=factor(Length,levels=Lorder)),width=0.9) +
  scale_x_continuous(breaks=c(84,86,89,92,94), labels=c("-5","-3","0","+3","+5")) +
  scale_y_continuous(expand = c(0, 0), breaks=c(0,30,60,90)) +
  scale_fill_manual(values=sRNAcols) +
  labs(x="amiRA1") +
  coord_cartesian(xlim = c(84,94), ylim = c(-4,120)) +
  theme_minimal() +
  theme(panel.grid.major.x=element_blank(), panel.grid.minor.x=element_blank(), panel.grid.minor.y=element_blank()) +
  theme(axis.text.x=element_text(colour = "black",size=12), axis.title.x = element_text(color="black",size=12)) +
  theme(axis.text.y=element_text(colour = "white",size=12), axis.title.y = element_text(color="white",vjust=1,size=12)) +
  theme(legend.position="none") +
  theme(plot.margin = unit(c(0, 0.1, 0, -0.1), "cm")) +
  labs(fill="RNA Length")

paMIR3_3 <- ggplot() +
  geom_segment(aes(x = 172, y = -4, xend = 172, yend = 100), linetype = "dashed", colour="grey70") +
  geom_hline(yintercept = 0, color = "grey70", size = 0.4) +
  geom_col(data=coverage3, aes(x=Position, y=RPM, fill=factor(Length,levels=Lorder)),width=0.9) +
  scale_x_continuous(breaks=c(167,169,172,175,177), labels=c("-5","-3","0","+3","+5")) +
  scale_y_continuous(expand = c(0, 0), breaks=c(0,30,60,90)) +
  scale_fill_manual(values=sRNAcols) +
  labs(x="amiRA4*") +
  coord_cartesian(xlim = c(167,177), ylim = c(-4,120)) +
  theme_minimal() +
  theme(panel.grid.major.x=element_blank(), panel.grid.minor.x=element_blank(), panel.grid.minor.y=element_blank()) +
  theme(axis.text.x=element_text(colour = "black",size=12), axis.title.x = element_text(color="grey40",size=12)) +
  theme(axis.text.y=element_text(colour = "white",size=12), axis.title.y = element_text(color="white",vjust=1,size=12)) +
  theme(legend.position="none") +
  theme(plot.margin = unit(c(0, 0.1, 0, -0.1), "cm")) +
  labs(fill="RNA Length")

paMIR3_4 <- ggplot() +
  geom_segment(aes(x = 227, y = -4, xend = 227, yend = 100), linetype = "dashed", colour="grey70") +
  geom_hline(yintercept = 0, color = "grey70", size = 0.4) +
  geom_col(data=coverage3, aes(x=Position, y=RPM, fill=factor(Length,levels=Lorder)),width=0.9) +
  scale_x_continuous(breaks=c(222,224,227,230,232), labels=c("-5","-3","0","+3","+5")) +
  scale_y_continuous(expand = c(0, 0), breaks=c(0,30,60,90)) +
  scale_fill_manual(values=sRNAcols) +
  labs(x="amiRA4") +
  coord_cartesian(xlim = c(222,232), ylim = c(-4,120)) +
  theme_minimal() +
  theme(panel.grid.major.x=element_blank(), panel.grid.minor.x=element_blank(), panel.grid.minor.y=element_blank()) +
  theme(axis.text.x=element_text(colour = "black",size=12), axis.title.x = element_text(color="black",size=12)) +
  theme(axis.text.y=element_text(colour = "white",size=12), axis.title.y = element_text(color="white",vjust=1,size=12)) +
  theme(legend.position="none") +
  theme(plot.margin = unit(c(0, 0.1, 0, -0.1), "cm")) +
  labs(fill="RNA Length")

paMIR3_5 <- ggplot() +
  geom_segment(aes(x = 310, y = -4, xend = 310, yend = 100), linetype = "dashed", colour="grey70") +
  geom_hline(yintercept = 0, color = "grey70", size = 0.4) +
  geom_col(data=coverage3, aes(x=Position, y=RPM, fill=factor(Length,levels=Lorder)),width=0.9) +
  scale_x_continuous(breaks=c(305,307,310,313,315), labels=c("-5","-3","0","+3","+5")) +
  scale_y_continuous(expand = c(0, 0), breaks=c(0,30,60,90)) +
  scale_fill_manual(values=sRNAcols) +
  labs(x="amiRC5*") +
  coord_cartesian(xlim = c(305,315), ylim = c(-4,120)) +
  theme_minimal() +
  theme(panel.grid.major.x=element_blank(), panel.grid.minor.x=element_blank(), panel.grid.minor.y=element_blank()) +
  theme(axis.text.x=element_text(colour = "black",size=12), axis.title.x = element_text(color="grey40",size=12)) +
  theme(axis.text.y=element_text(colour = "white",size=12), axis.title.y = element_text(color="white",vjust=1,size=12)) +
  theme(legend.position="none") +
  theme(plot.margin = unit(c(0, 0.1, 0, -0.1), "cm")) +
  labs(fill="RNA Length")

paMIR3_6 <- ggplot() +
  geom_segment(aes(x = 365, y = -4, xend = 365, yend = 100), linetype = "dashed", colour="grey70") +
  geom_hline(yintercept = 0, color = "grey70", size = 0.4) +
  geom_col(data=coverage3, aes(x=Position, y=RPM, fill=factor(Length,levels=Lorder)),width=0.9) +
  scale_x_continuous(breaks=c(360,362,365,368,370), labels=c("-5","-3","0","+3","+5")) +
  scale_y_continuous(expand = c(0, 0), breaks=c(0,30,60,90)) +
  scale_fill_manual(values=sRNAcols) +
  labs(x="amiRC5") +
  coord_cartesian(xlim = c(360,370), ylim = c(-4,120)) +
  theme_minimal() +
  theme(panel.grid.major.x=element_blank(), panel.grid.minor.x=element_blank(), panel.grid.minor.y=element_blank()) +
  theme(axis.text.x=element_text(colour = "black",size=12), axis.title.x = element_text(color="black",size=12)) +
  theme(axis.text.y=element_text(colour = "white",size=12), axis.title.y = element_text(color="white",vjust=1,size=12)) +
  theme(legend.position="none") +
  theme(plot.margin = unit(c(0, 0.1, 0, -0.1), "cm")) +
  labs(fill="RNA Length")

merge3 <- ggarrange(paMIR3_1,paMIR3_2,paMIR3_3,paMIR3_4,paMIR3_5,paMIR3_6,
                    widths = c(1,1,1,1,1,1),
                    ncol = 6, nrow = 1, align = "h")

# Make plot paMIR4
paMIR4_1 <- ggplot() +
  geom_segment(aes(x = 33, y = -6, xend = 33, yend = 205), linetype = "dashed", colour="grey40") +
  geom_hline(yintercept = 0, color = "grey40", size = 0.4) +
  geom_col(data=coverage4, aes(x=Position, y=RPM, fill=factor(Length,levels=Lorder)),width=0.9) +
  scale_x_continuous(breaks=c(28,33,38), labels=c("-5","0","+5")) +
  scale_y_continuous(expand = c(0, 0), breaks=c(0,65,130,195)) +
  scale_fill_manual(values=sRNAcols) +
  labs(x="amiRA1*", y="RPM") +
  coord_cartesian(xlim = c(28,38), ylim = c(-6,265)) +
  theme_minimal() +
  theme(panel.grid.major.x=element_blank(), panel.grid.minor.x=element_blank(), panel.grid.minor.y=element_blank()) +
  theme(axis.text.x=element_text(colour = "black",size=12), axis.title.x = element_text(color="grey40",size=12)) +
  theme(axis.text.y=element_text(colour = "black",size=12), axis.title.y = element_text(color="black",vjust=1,size=12)) +
  theme(legend.position="none") +
  theme(plot.margin = unit(c(0, 0, 0, 0), "cm")) +
  labs(fill="RNA Length")

paMIR4_2 <- ggplot() +
  geom_segment(aes(x = 132, y = -6, xend = 132, yend = 205), linetype = "dashed", colour="grey40") +
  geom_hline(yintercept = 0, color = "grey40", size = 0.4) +
  geom_col(data=coverage4, aes(x=Position, y=RPM, fill=factor(Length,levels=Lorder)),width=0.9) +
  scale_x_continuous(breaks=c(127,132,137), labels=c("-5","0","+5")) +
  scale_y_continuous(expand = c(0, 0), breaks=c(0,65,130,195)) +
  scale_fill_manual(values=sRNAcols) +
  labs(x="amiRA1") +
  coord_cartesian(xlim = c(127,137), ylim = c(-6,265)) +
  theme_minimal() +
  theme(panel.grid.major.x=element_blank(), panel.grid.minor.x=element_blank(), panel.grid.minor.y=element_blank()) +
  theme(axis.text.x=element_text(colour = "black",size=12), axis.title.x = element_text(color="black",size=12)) +
  theme(axis.text.y=element_text(colour = "white",size=12), axis.title.y = element_text(color="white",vjust=1,size=12)) +
  theme(legend.position="none") +
  theme(plot.margin = unit(c(0, 0.1, 0, -0.1), "cm")) +
  labs(fill="RNA Length")

paMIR4_3 <- ggplot() +
  geom_segment(aes(x = 249, y = -6, xend = 249, yend = 205), linetype = "dashed", colour="grey40") +
  geom_hline(yintercept = 0, color = "grey40", size = 0.4) +
  geom_col(data=coverage4, aes(x=Position, y=RPM, fill=factor(Length,levels=Lorder)),width=0.9) +
  scale_x_continuous(breaks=c(244,249,254), labels=c("-5","0","+5")) +
  scale_y_continuous(expand = c(0, 0), breaks=c(0,65,130,195)) +
  scale_fill_manual(values=sRNAcols) +
  labs(x="amiRA4*") +
  coord_cartesian(xlim = c(244,254), ylim = c(-6,265)) +
  theme_minimal() +
  theme(panel.grid.major.x=element_blank(), panel.grid.minor.x=element_blank(), panel.grid.minor.y=element_blank()) +
  theme(axis.text.x=element_text(colour = "black",size=12), axis.title.x = element_text(color="grey40",size=12)) +
  theme(axis.text.y=element_text(colour = "white",size=12), axis.title.y = element_text(color="white",vjust=1,size=12)) +
  theme(legend.position="none") +
  theme(plot.margin = unit(c(0, 0.1, 0, -0.1), "cm")) +
  labs(fill="RNA Length")

paMIR4_4 <- ggplot() +
  geom_segment(aes(x = 292, y = -6, xend = 292, yend = 205), linetype = "dashed", colour="grey40") +
  geom_hline(yintercept = 0, color = "grey40", size = 0.4) +
  geom_col(data=coverage4, aes(x=Position, y=RPM, fill=factor(Length,levels=Lorder)),width=0.9) +
  scale_x_continuous(breaks=c(287,292,297), labels=c("-5","0","+5")) +
  scale_y_continuous(expand = c(0, 0), breaks=c(0,65,130,195)) +
  scale_fill_manual(values=sRNAcols) +
  labs(x="amiRA4") +
  coord_cartesian(xlim = c(287,297), ylim = c(-6,265)) +
  theme_minimal() +
  theme(panel.grid.major.x=element_blank(), panel.grid.minor.x=element_blank(), panel.grid.minor.y=element_blank()) +
  theme(axis.text.x=element_text(colour = "black",size=12), axis.title.x = element_text(color="black",size=12)) +
  theme(axis.text.y=element_text(colour = "white",size=12), axis.title.y = element_text(color="white",vjust=1,size=12)) +
  theme(legend.position="none") +
  theme(plot.margin = unit(c(0, 0.1, 0, -0.1), "cm")) +
  labs(fill="RNA Length")

paMIR4_5 <- ggplot() +
  geom_segment(aes(x = 387, y = -6, xend = 387, yend = 205), linetype = "dashed", colour="grey40") +
  geom_hline(yintercept = 0, color = "grey40", size = 0.4) +
  geom_col(data=coverage4, aes(x=Position, y=RPM, fill=factor(Length,levels=Lorder)),width=0.9) +
  scale_x_continuous(breaks=c(382,387,392), labels=c("-5","0","+5")) +
  scale_y_continuous(expand = c(0, 0), breaks=c(0,65,130,195)) +
  scale_fill_manual(values=sRNAcols) +
  labs(x="amiRB5*") +
  coord_cartesian(xlim = c(382,392), ylim = c(-6,265)) +
  theme_minimal() +
  theme(panel.grid.major.x=element_blank(), panel.grid.minor.x=element_blank(), panel.grid.minor.y=element_blank()) +
  theme(axis.text.x=element_text(colour = "black",size=12), axis.title.x = element_text(color="grey40",size=12)) +
  theme(axis.text.y=element_text(colour = "white",size=12), axis.title.y = element_text(color="white",vjust=1,size=12)) +
  theme(legend.position="none") +
  theme(plot.margin = unit(c(0, 0.1, 0, -0.1), "cm")) +
  labs(fill="RNA Length")

paMIR4_6 <- ggplot() +
  geom_segment(aes(x = 433, y = -6, xend = 433, yend = 205), linetype = "dashed", colour="grey40") +
  geom_hline(yintercept = 0, color = "grey40", size = 0.4) +
  geom_col(data=coverage4, aes(x=Position, y=RPM, fill=factor(Length,levels=Lorder)),width=0.9) +
  scale_x_continuous(breaks=c(428,433,438), labels=c("-5","0","+5")) +
  scale_y_continuous(expand = c(0, 0), breaks=c(0,65,130,195)) +
  scale_fill_manual(values=sRNAcols) +
  labs(x="amiRB5") +
  coord_cartesian(xlim = c(428,438), ylim = c(-6,265)) +
  theme_minimal() +
  theme(panel.grid.major.x=element_blank(), panel.grid.minor.x=element_blank(), panel.grid.minor.y=element_blank()) +
  theme(axis.text.x=element_text(colour = "black",size=12), axis.title.x = element_text(color="black",size=12)) +
  theme(axis.text.y=element_text(colour = "white",size=12), axis.title.y = element_text(color="white",vjust=1,size=12)) +
  theme(legend.position="none") +
  theme(plot.margin = unit(c(0, 0.1, 0, -0.1), "cm")) +
  labs(fill="RNA Length")

paMIR4_7 <- ggplot() +
  geom_segment(aes(x = 526, y = -6, xend = 526, yend = 205), linetype = "dashed", colour="grey40") +
  geom_hline(yintercept = 0, color = "grey40", size = 0.4) +
  geom_col(data=coverage4, aes(x=Position, y=RPM, fill=factor(Length,levels=Lorder)),width=0.9) +
  scale_x_continuous(breaks=c(521,526,531), labels=c("-5","0","+5")) +
  scale_y_continuous(expand = c(0, 0), breaks=c(0,65,130,195)) +
  scale_fill_manual(values=sRNAcols) +
  labs(x="amiRC4*") +
  coord_cartesian(xlim = c(521,531), ylim = c(-6,265)) +
  theme_minimal() +
  theme(panel.grid.major.x=element_blank(), panel.grid.minor.x=element_blank(), panel.grid.minor.y=element_blank()) +
  theme(axis.text.x=element_text(colour = "black",size=12), axis.title.x = element_text(color="grey40",size=12)) +
  theme(axis.text.y=element_text(colour = "white",size=12), axis.title.y = element_text(color="white",vjust=1,size=12)) +
  theme(legend.position="none") +
  theme(plot.margin = unit(c(0, 0.1, 0, -0.1), "cm")) +
  labs(fill="RNA Length")

paMIR4_8 <- ggplot() +
  geom_segment(aes(x = 571, y = -6, xend = 571, yend = 205), linetype = "dashed", colour="grey40") +
  geom_hline(yintercept = 0, color = "grey40", size = 0.4) +
  geom_col(data=coverage4, aes(x=Position, y=RPM, fill=factor(Length,levels=Lorder)),width=0.9) +
  scale_x_continuous(breaks=c(566,571,576), labels=c("-5","0","+5")) +
  scale_y_continuous(expand = c(0, 0), breaks=c(0,65,130,195)) +
  scale_fill_manual(values=sRNAcols) +
  labs(x="amiRC4") +
  coord_cartesian(xlim = c(566,576), ylim = c(-6,265)) +
  theme_minimal() +
  theme(panel.grid.major.x=element_blank(), panel.grid.minor.x=element_blank(), panel.grid.minor.y=element_blank()) +
  theme(axis.text.x=element_text(colour = "black",size=12), axis.title.x = element_text(color="black",size=12)) +
  theme(axis.text.y=element_text(colour = "white",size=12), axis.title.y = element_text(color="white",vjust=1,size=12)) +
  theme(legend.position="none") +
  theme(plot.margin = unit(c(0, 0.1, 0, -0.1), "cm")) +
  labs(fill="RNA Length")

paMIR4_9 <- ggplot() +
  geom_segment(aes(x = 669, y = -6, xend = 669, yend = 205), linetype = "dashed", colour="grey40") +
  geom_hline(yintercept = 0, color = "grey40", size = 0.4) +
  geom_col(data=coverage4, aes(x=Position, y=RPM, fill=factor(Length,levels=Lorder)),width=0.9) +
  scale_x_continuous(breaks=c(664,669,674), labels=c("-5","0","+5")) +
  scale_y_continuous(expand = c(0, 0), breaks=c(0,65,130,195)) +
  scale_fill_manual(values=sRNAcols) +
  labs(x="amiRC5*") +
  coord_cartesian(xlim = c(664,674), ylim = c(-6,265)) +
  theme_minimal() +
  theme(panel.grid.major.x=element_blank(), panel.grid.minor.x=element_blank(), panel.grid.minor.y=element_blank()) +
  theme(axis.text.x=element_text(colour = "black",size=12), axis.title.x = element_text(color="grey40",size=12)) +
  theme(axis.text.y=element_text(colour = "white",size=12), axis.title.y = element_text(color="white",vjust=1,size=12)) +
  theme(legend.position="none") +
  theme(plot.margin = unit(c(0, 0.1, 0, -0.1), "cm")) +
  labs(fill="RNA Length")

paMIR4_10 <- ggplot() +
  geom_segment(aes(x = 714, y = -6, xend = 714, yend = 205), linetype = "dashed", colour="grey40") +
  geom_hline(yintercept = 0, color = "grey40", size = 0.4) +
  geom_col(data=coverage4, aes(x=Position, y=RPM, fill=factor(Length,levels=Lorder)),width=0.9) +
  scale_x_continuous(breaks=c(709,714,719), labels=c("-5","0","+5")) +
  scale_y_continuous(expand = c(0, 0), breaks=c(0,65,130,195)) +
  scale_fill_manual(values=sRNAcols) +
  labs(x="amiRC5") +
  coord_cartesian(xlim = c(709,719), ylim = c(-6,265)) +
  theme_minimal() +
  theme(panel.grid.major.x=element_blank(), panel.grid.minor.x=element_blank(), panel.grid.minor.y=element_blank()) +
  theme(axis.text.x=element_text(colour = "black",size=12), axis.title.x = element_text(color="black",hjust=0.8,size=12)) +
  theme(axis.text.y=element_text(colour = "white",size=12), axis.title.y = element_text(color="white",vjust=1,size=12)) +
  theme(legend.position="none") +
  theme(plot.margin = unit(c(0, 0.1, 0, -0.1), "cm")) +
  labs(fill="RNA Length")

merge4 <- ggarrange(paMIR4_1,paMIR4_2,paMIR4_3,paMIR4_4,paMIR4_5,paMIR4_6,paMIR4_7,paMIR4_8,paMIR4_9,paMIR4_10,
                    widths = c(1,1,1,1,1,1,1,1,1,1),
                    ncol = 10, nrow = 1, align = "h")

# Merge plots in one single graph
title <- ggdraw() + 
  draw_label(expression(bolditalic("paMIRNA")~bold("5' end coverage in")~bolditalic("S. lycopersicum")),
             fontface = 'bold', hjust = 0.5, size = 16)

xlabel <- ggdraw() + 
  draw_label("5' end position of reads relative to amiRNA and amiRNA* (bp)",
             hjust = 0.5, size = 13)

merge5 <- ggarrange(title,merge1,merge2,merge3,merge4,xlabel,
                    heights = c(1/25,1/5,1/5,1/5,1/5,1/30),
                    ncol = 1, nrow = 6, align = "v",
                    labels = c("","paMIR1","paMIR2","paMIR3","paMIR4",""),
                    font.label = list(size = 16, face="bold.italic"),
                    hjust=0, vjust=c(1,1,1.3,1.5,1.3,1))

merge5

# Add legend
legend <- get_legend(one + theme(legend.position="right", legend.direction = "vertical") +
                       guides(fill = guide_legend(ncol = 1)))

plot_grid(merge5, legend, rel_widths = c(3, .4))
```
