## Supplemental Tables, Code, and Data for "Expression and processing of polycistronic artificial microRNAs and *trans*-acting siRNAs in *Solanum lycopersicum* and *Nicotiana benthamiana*": Supplemental_Code_5.R

```
# This code generates Figure 6

# Activate required non-base packages
library(ggplot2)
library(cowplot)
library(ggpmisc)

# Set the working directory by replacing "/your/path" with
# the complete path to the directory where this file is
setwd("/your/path")

# Load the data
correlation <- read.table(file="stemloopPCR_correlation.txt", head=TRUE)

# Define colors
#A1:firebrick1
#A3:dodgerblue
#A4:chartreuse3
#B1:purple1
#B5:orchid1
#C4:saddlebrown
#C5:orange
#E1:aquamarine2

# Make plots
title <- ggdraw() + 
  draw_label(expression(bold("sRNA-seq vs stem-loop RT-PCR")),
             fontface = 'bold', hjust = 0.5, size = 15.5)

correlation$Gene <- factor(correlation$Gene,
                           levels=c("single","paMIR1","paMIR2","paMIR3","paMIR4","aTAS3"),
                           labels=c("monocistronic\ngenes","paMIR1","paMIR2","paMIR3","paMIR4","aTAS3"))

tomato <- ggplot(subset(correlation, Species == "tomato"),
                 aes(x=Seq, y=PCR)) +
  geom_point(aes(shape=Gene,color=Sample),size=3) +
  scale_shape_manual(values=c(16,10,17,9,18,8))+ 
  scale_color_manual(values=c('firebrick1','dodgerblue','chartreuse3',
                              'purple1','orchid1','saddlebrown','orange',
                              'aquamarine2'))+
  geom_smooth(method=lm, se=FALSE, color='black', formula = y ~ x) +
  stat_poly_eq(formula = y ~ x, parse = TRUE, size = 5,
               label.x = 0.9, label.y = 0.4) +
  labs(title="S. lycopersicum",
       x="Log2[RPM]", y = "Ct(miR166)-Ct(amiRNA)",
       shape="aMIRNA gene", colour="amiRNA measured") +
  guides(color=guide_legend(order = 2,override.aes = list(shape=15, size=5)),
         shape=guide_legend(order = 1)) +
  coord_cartesian(xlim = c(-2,14), ylim = c(-20,-1)) +
  theme_classic() +
  theme(legend.text=element_text(size=13),legend.title=element_text(size=13)) +
  theme(axis.text=element_text(size=13, color="black"),axis.title=element_text(size=13)) +
  theme(plot.title = element_text(size=13, face=4, vjust=-10, hjust=0.1)) +
  theme(legend.position = "none")

Nbenthamiana <- ggplot(subset(correlation, Species == "Nbenthamiana"),
                       aes(x=Seq, y=PCR)) +
  geom_point(aes(shape=Gene,color=Sample),size=3) +
  scale_shape_manual(values=c(16,10,17,9,18,8))+ 
  scale_color_manual(values=c('firebrick1','dodgerblue','chartreuse3',
                              'purple1','orchid1','saddlebrown','orange',
                              'aquamarine2'))+
  geom_smooth(method=lm, se=FALSE, color='black', formula = y ~ x) +
  stat_poly_eq(formula = y ~ x, parse = TRUE, size = 5,
               label.x = 0.9, label.y = 0.4) +
  labs(title="N. benthamiana",
       x="Log2[RPM]", y = "Ct(miR166)-Ct(amiRNA)",
       shape="aMIRNA gene", colour="amiRNA measured") +
  guides(color=guide_legend(order = 2,override.aes = list(shape=15, size=5)),
         shape=guide_legend(order = 1)) +
  coord_cartesian(xlim = c(-2,14), ylim = c(-20,-1)) +
  theme_classic() +
  theme(legend.text=element_text(size=13),legend.title=element_text(size=13)) +
  theme(axis.text=element_text(size=13, color="black"),axis.title=element_text(size=13)) +
  theme(plot.title = element_text(size=13, face=4, vjust=-10, hjust=0.1)) +
  theme(legend.position = "none")

# Merge plots in one graph
merge1 <- plot_grid(title,tomato,Nbenthamiana,
                     labels = c("","A","B"),
                     label_size = 18,
                     hjust = -0.1, vjust = c(1.5,1.5,1.5),
                     align = "v", axis = "l",
                     ncol= 1, nrow = 3,
                     rel_heights = c( 1/40,1/3,1/3))

to_isolate_legend <- ggplot(subset(correlation, Species == "Nbenthamiana"),
                       aes(x=Seq, y=PCR)) +
  geom_point(aes(shape=Gene,color=Sample),size=3) +
  scale_shape_manual(values=c(16,10,17,9,18,8), labels=c("monocistronic\ngenes",
                    expression(italic("paMIR1")),expression(italic("paMIR2")),expression(italic("paMIR3")),expression(italic("paMIR4")),expression(italic("aTAS3"))))+ 
  scale_color_manual(values=c('firebrick1','dodgerblue','chartreuse3',
                              'purple1','orchid1','saddlebrown','orange',
                              'aquamarine2'))+
  geom_smooth(method=lm, se=FALSE, color='black', formula = y ~ x) +
  stat_poly_eq(aes(label = paste(..eq.label.., ..rr.label.., sep = "~~~")), 
               label.x.npc = "right", label.y.npc = 0.1,
               formula = y ~ x, parse = TRUE, size = 5) +
  labs(title="N. benthamiana",
       x="Log2[RPM]", y = "Ct(miR166)-Ct(amiRNA)",
       shape=expression(italic("aMIRNA")~"gene"), colour="amiRNA measured") +
  guides(color=guide_legend(order = 2,override.aes = list(shape=15, size=5)),
         shape=guide_legend(order = 1)) +
  coord_cartesian(xlim = c(-2,14), ylim = c(-20,-1)) +
  theme_classic() +
  theme(legend.text=element_text(size=13),legend.text.align = 0, legend.title=element_text(size=13)) +
  theme(axis.text=element_text(size=13, color="black"),axis.title=element_text(size=13)) +
  theme(plot.title = element_text(size=13, face=4, vjust=-10, hjust=0.1))

legend <- get_legend(to_isolate_legend)

plot_grid(merge1, legend, rel_widths = c(2.8, .8))
```
