## Supplemental Tables, Code, and Data for "Expression and processing of polycistronic artificial microRNAs and *trans*-acting siRNAs in *Solanum lycopersicum* and *Nicotiana benthamiana*": Supplemental_Code_6.R

```
# This code generates Figure 7

# Set the working directory by replacing "/your/path" with
# the complete path to the directory where this file is
setwd("/your/path")

# Load the data
qRTPCR <- read.table(file="2020_05_17_all_qRT-PCRs.txt", head=TRUE)

qRTPCR$sample = factor(qRTPCR$sample, unique(qRTPCR$sample))
qRTPCR$sample

# Define colors
# PP2C1: blue
# PP2C2: deepskyblue
# PP2C3: cyan
# SAHH1: darkred
# SAHH2: red
# SAHH3: orange
# HyPRP: green
# UGT75C1: magenta

# Make plot
par(mar = c(6, 6, 3, 2) + 0.1,xpd=TRUE)

my_lab <- expression(paste("Relative Accumulation (normalized to ", italic("SlSAND"), ")"))

boxplot(qRTPCR$value~qRTPCR$sample,
        ylim=c(0,3.1), yaxs="i",
        xlim=c(1,42), xaxs="i",
        vertical=TRUE,
        xaxt = "n",
        border=c("blue","blue","blue","deepskyblue","cyan","darkred","red","orange",
                 "darkred","red","orange","green","green","magenta","blue","deepskyblue","cyan",
                 "blue","deepskyblue","cyan","darkred","red","orange",
                 "blue","deepskyblue","cyan","green","blue","deepskyblue","cyan",
                 "darkred","red","orange","green","blue","deepskyblue","cyan",
                 "darkred","red","orange","green","magenta"),
        cex=1,cex.axis=1.2, cex.lab=1.2,
        outcex=0.75,
        ylab=my_lab,
        xlab = NA,
        las=2,
        main=expression(bold("Target gene accumulation in")~bolditalic("S. lycopersicum")), cex.main=1.3)

stripchart(qRTPCR$value~qRTPCR$sample,
           ylim=c(0,3.1), yaxs="i",
           xlim=c(1,42), xaxs="i",
           vertical=TRUE,
           xaxt = "n",
           col= c("blue","blue","blue","deepskyblue","cyan","darkred","red","orange",
                  "darkred","red","orange","green","green","magenta","blue","deepskyblue","cyan",
                  "blue","deepskyblue","cyan","darkred","red","orange",
                  "blue","deepskyblue","cyan","green","blue","deepskyblue","cyan",
                  "darkred","red","orange","green","blue","deepskyblue","cyan",
                  "darkred","red","orange","green","magenta"),
           pch=16,
           cex=1,cex.axis = 1.2, cex.lab = 1.2,
           method="jitter",
           ylab= NA, xlab = NA,
           add=TRUE)

text(x=32.5, y=-0.45, adj=1.5, srt = 0, cex =1.2,
     labels="Agroinfiltrated sample", xpd=TRUE,
     col="black")

abline(v=c(1.5,2.5,5.5,8.5,11.5,12.5,13.5,14.5,17.5,23.5,27.5,34.5), lty=1, col="grey", xpd=FALSE)
abline(h=1, lty=3, xpd=FALSE)

text(x=c(1,2,4,7,10,12,13,14,16,20.5,25.5,31,38.5), y=-0.05, adj=1, srt = 90, cex =1.2,
     labels=c(expression(italic("aMIRA1")),expression(italic("aMIRA3")),
              expression(italic("aMIRA4")),expression(italic("aMIRB1")),
              expression(italic("aMIRB5")),expression(italic("aMIRC4")),
              expression(italic("aMIRC5")),expression(italic("aMIRE1")),
              expression(italic("paMIR1")),expression(italic("paMIR2")),
              expression(italic("paMIR3")),expression(italic("paMIR4")),
              expression(italic("aTAS3"))), xpd=TRUE, col="black")

legend(36,3, c(expression(italic("SlPP2C1")),expression(italic("SlPP2C2")),
                   expression(italic("SlPP2C3")),expression(italic("SlSAHH1")),
                   expression(italic("SlSAHH2")),expression(italic("SlSAHH3")),
                   expression(italic("SlHyPRP1")),expression(italic("SlUGT75C1"))),
       pch=16, col=c("blue","deepskyblue","cyan","darkred","red","orange","green","magenta"),
       xpd=TRUE, pt.cex = 1.5, cex = 1, inset=0.03,
       title="Target gene", title.adj=0.2, box.lty=0) 
```
