## Supplemental Tables, Code, and Data for "Expression and processing of polycistronic artificial microRNAs and *trans*-acting siRNAs in *Solanum lycopersicum* and *Nicotiana benthamiana*": Supplemental_Code_7.pl

```
#!/usr/bin/perl -w
$version = 0.2;
$usage = "sRNA_coverage.pl version $version

USAGE: sRNA_coverage.pl [bam] {[chr] | [chr:start-stop]}

REQUIRES: samtools , indexed BAM file";

unless(($ARGV[0]) and ($ARGV[1])) {
    die "$usage";
}

$temp_bam = $ARGV[0];
$temp_bam =~ s/\.bam/_temp\.bam/;

# Get total reads in file
open(COUNT, "samtools view -c $ARGV[0] |");
$count = <COUNT>;
close COUNT;
chomp $count;
$Mcount = $count / 1E6;

# Header
print "Chromosome\tPosition\tReads\tRPM\tLength\tStrand\n";

for($i = 19; $i <= 25; ++$i) {
# sense
    open(SAM, "samtools view -h -F 16 $ARGV[0] $ARGV[1] |");
    open(BAMOUT, "| samtools view -b - > $temp_bam");
    while(<SAM>) {
	if($_ =~ /^\@/) {
	    print BAMOUT $_;
	} else {
	    @fields = split ("\t", $_);
	    if($i == 19) {
		if((length $fields[9]) <= $i) {
		    print BAMOUT $_;
		}
	    } elsif ($i == 25) {
		if((length $fields[9]) >= $i) {
		    print BAMOUT $_;
		}
	    } else {
		if((length $fields[9]) == $i) {
		    print BAMOUT $_;
		}
	    }
	}
    }
    close SAM;
    close BAMOUT;
    system ("samtools index $temp_bam");
    open(DEPTH, "samtools depth -a -d 0 $temp_bam -r $ARGV[1] |");
    while (<DEPTH>) {
	chomp;
	print $_;
	@fields = split ("\t", $_);
	$rpm = $fields[2] / $Mcount;
	print "\t$rpm";
	if($i == 19) {
	    print "\t" . '<20 nts' . "\tplus\n";
	} elsif ($i == 25) {
	    print "\t" . '>24 nts' . "\tplus\n";
	} else {
	    print "\t" . "$i nts" . "\tplus\n";
	}
    }
    close DEPTH;
    system ("rm -f $temp_bam*");

# antisense
    open(SAM, "samtools view -h -f 16 $ARGV[0] $ARGV[1] |");
    open(BAMOUT, "| samtools view -b - > $temp_bam");
    while(<SAM>) {
	if($_ =~ /^\@/) {
	    print BAMOUT $_;
	} else {
	    @fields = split ("\t", $_);
	    if($i == 19) {
		if((length $fields[9]) <= $i) {
		    print BAMOUT $_;
		}
	    }
	    elsif ($i == 25) {
		if((length $fields[9]) >= $i) {
		    print BAMOUT $_;
		}
	    } else {
		if((length $fields[9]) == $i) {
		    print BAMOUT $_;
		}
	    }
	}
    }
    close SAM;
    close BAMOUT;
    system("samtools index $temp_bam");
    open(DEPTH, "samtools depth -a -d 0 $temp_bam -r $ARGV[1] |");
    while (<DEPTH>) {
	chomp;
	@fields = split ("\t", $_);
	$minus = 0 - $fields[2];
	$fields[2] = $minus;
	$out = join("\t", @fields);
	print $out;
	$rpm = $minus / $Mcount;
	print "\t$rpm";
	if($i == 19) {
	    print "\t" . '<20 nts' . "\tminus\n";
	} elsif ($i == 25) {
	    print "\t" . '>24 nts' . "\tminus\n";
	} else {
	    print "\t" . "$i nts" . "\tminus\n";
	}
    }
    close DEPTH;
    system ("rm -f $temp_bam*");
}
```
