## Supplemental Tables, Code, and Data for "Expression and processing of polycistronic artificial microRNAs and *trans*-acting siRNAs in *Solanum lycopersicum* and *Nicotiana benthamiana*": Supplemental_Code_7.R

```
# This code generates Supplemental Figure 7

# Set the working directory by replacing "/your/path" with
# the complete path to the directory where this file is
setwd("/your/path")

# Load the data
stemloop <- read.table(file="tot_selected.txt", head=TRUE)

# Just for safety in case there is a lingering attachment
detach(stemloop)

# Attach the namespace
attach(stemloop)

stemloop$amiRNA = factor(stemloop$amiRNA, unique(stemloop$amiRNA))
stemloop$amiRNA

# Make plot
par(mar = c(9, 7, 3, 1) + 0.1,xpd=TRUE)

boxplot(stemloop$accumulation~stemloop$amiRNA,
        ylim=c(-1,3.5), yaxs="i",
        xlim=c(2.3,58.7), xaxs="i",
        vertical=TRUE,
        xaxt = "n",
        yaxt = "n",
        border=c("grey10","dodgerblue1","grey10","dodgerblue1","grey10","dodgerblue1","grey10","dodgerblue1","grey10","dodgerblue1",
                 "grey10","dodgerblue1","grey10","dodgerblue1","grey10","dodgerblue1","grey10","dodgerblue1","grey10","dodgerblue1",
                 "grey10","dodgerblue1","grey10","dodgerblue1","grey10","dodgerblue1","grey10","dodgerblue1","grey10","dodgerblue1",
                 "grey10","dodgerblue1","grey10","dodgerblue1","grey10","dodgerblue1","grey10","dodgerblue1","grey10","dodgerblue1",
                 "grey10","dodgerblue1","grey10","dodgerblue1","grey10","dodgerblue1","grey10","dodgerblue1","grey10","dodgerblue1",
                 "grey10","dodgerblue1","grey10","dodgerblue1","grey10","dodgerblue1","grey10","dodgerblue1","grey10","dodgerblue1"),
        cex=1.2,cex.axis=1.2, cex.lab=1.2,
        outcex=0.75,
        ylab="Log2 [Normalized (Ct miR166 - Ct amiRNA)]",
        xlab = NA,
        main="amiRNA accumulation via stem-loop qRT-PCR", cex.main=1.3)
axis(2, at = seq(-1, 3, by = 0.5), las=2,cex=1,cex.axis=1.2, cex.lab=1.2,col = "black")

stripchart(stemloop$accumulation~stemloop$amiRNA,
           ylim=c(-1,3.5), yaxs="i",
           xlim=c(2.3,58.7), xaxs="i",
           vertical=TRUE,
           xaxt = "n",
           col= c("grey10","dodgerblue1","grey10","dodgerblue1","grey10","dodgerblue1","grey10","dodgerblue1","grey10","dodgerblue1",
                  "grey10","dodgerblue1","grey10","dodgerblue1","grey10","dodgerblue1","grey10","dodgerblue1","grey10","dodgerblue1",
                  "grey10","dodgerblue1","grey10","dodgerblue1","grey10","dodgerblue1","grey10","dodgerblue1","grey10","dodgerblue1",
                  "grey10","dodgerblue1","grey10","dodgerblue1","grey10","dodgerblue1","grey10","dodgerblue1","grey10","dodgerblue1",
                  "grey10","dodgerblue1","grey10","dodgerblue1","grey10","dodgerblue1","grey10","dodgerblue1","grey10","dodgerblue1",
                  "grey10","dodgerblue1","grey10","dodgerblue1","grey10","dodgerblue1","grey10","dodgerblue1","grey10","dodgerblue1"),
           pch=16,
           cex=1.2,cex.axis = 1.2, cex.lab = 1.2,
           method="jitter",
           ylab= NA, xlab = NA,
           add=TRUE)

abline(v=c(2.5,4.5,6.5,8.5,10.5,12.5,14.5,16.5,22.5,28.5,34.5,44.5), lty=1, col="grey", xpd=FALSE)
abline(h=0, lty=3, xpd=FALSE)

text(x=c(1.5,3.5,5.5,7.5,9.5,11.5,13.5,15.5,19.5,25.5,31.5,39.5,52.5),
     y=-1.7, adj=1, srt = 90, cex =1.2,
     labels=c(expression(italic("aMIRA1")),expression(italic("aMIRA3")),expression(italic("aMIRA4")),
              expression(italic("aMIRB1")),expression(italic("aMIRB5")),expression(italic("aMIRC4")),
              expression(italic("aMIRC5")),expression(italic("aMIRE1")),expression(italic("paMIR1")),
              expression(italic("paMIR2")),expression(italic("paMIR3")),expression(italic("paMIR4")),
              expression(italic("aTAS3"))), xpd=TRUE, col="black")

text(x=-0.2, y=-1.9, adj=1, srt = 0, cex =1.2,
     labels="Agroinfiltrated\nsample:", xpd=TRUE,
     col="black")

text(x=c(1.5,3.5,5.5,7.5,9.5,11.5,13.5,15.5,17.5,19.5,21.5,23.5,25.5,27.5,29.5,
         31.5,33.5,35.5,37.5,39.5,41.5,43.5,45.5,47.5,49.5,51.5,53.5,55.5,57.5,59.5),
     y=-1.1, adj=1, srt = 90, cex =1.2,
     labels=c("amiRA1","amiRA3","amiRA4","amiRB1","amiRB5","amiRC4","amiRC5","amiRE1",
              "amiRA1","amiRA3","amiRA4","amiRA1","amiRA4","amiRB1","amiRA1","amiRA4","amiRC5",
              "amiRA1","amiRA4","amiRB5","amiRC4","amiRC5",
              "amiRA3","amiRA1","amiRA4","amiRC5","amiRB5","amiRC4","amiRE1","amiRB1"), xpd=TRUE,
     col="black")

text(x=-0.2, y=-1.35, adj=1, srt = 0, cex =1.2,
     labels="amiRNA\nmeasured:", xpd=TRUE,
     col="black")

text(x=c(19,21,45,47,49,51,53,55,57,59), y=-0.4,
     adj=1, srt = 90, cex =1,
     labels="undetected", xpd=TRUE,
     col="grey10")

segments(c(0.7,2.7,4.7,6.7,8.7,10.7,12.7,14.7,16.7,22.7,28.7,34.7,44.7),
         c(-1.65,-1.65,-1.65,-1.65,-1.65,-1.65,-1.65,-1.65,-1.65,-1.65,
           -1.65,-1.65,-1.65),
         c(2.3,4.3,6.3,8.3,10.3,12.3,14.3,16.3,22.3,28.3,34.3,44.3,60.3),
         c(-1.65,-1.65,-1.65,-1.65,-1.65,-1.65,-1.65,-1.65,-1.65,-1.65,
           -1.65,-1.65,-1.65),
         col="black",
         lwd=c(1.5,1.5,1.5,1.5,1.5,1.5,1.5,1.5,1.5,1.5,
               1.5,1.5,1.5),
         xpd=TRUE)

legend(47,3, c(expression(italic("S. lycopersicum")),expression(italic("N. benthamiana"))),
       pch=16, col=c("grey10","dodgerblue1"),
       xpd=TRUE, pt.cex = 1.3, cex = 1.2, inset=0.03,
       title="Species", title.adj=0.1, title.col="black",box.lty=0) 
```
